# The Na^+^-pumping mechanism driven by redox reactions in the NADH-quinone oxidoreductase from *Vibrio cholerae* relies on dynamic conformational changes

**DOI:** 10.1101/2025.06.01.656757

**Authors:** Moe Ishikawa-Fukuda, Takehito Seki, Jun-ichi Kishikawa, Masuya Takahiro, Kei-ichi Okazaki, Takayuki Kato, Blanca Barquera, Hideto Miyoshi, Masatoshi Murai

## Abstract

The Na^+^-pumping NADH-quinone oxidoreductase (Na^+^-NQR) is a key respiratory enzyme in many marine and pathogenic bacteria that couples electron transfer to Na^+^-pumping across the membrane. Earlier X-ray and cryo-EM structures of Na^+^-NQR from *Vibrio cholerae* suggested that the subunits harboring redox cofactors undergo conformational changes during catalytic turnover. However, these proposed rearrangements have not yet been confirmed. Here, we have identified at least five distinct conformational states of Na^+^-NQR using: mutants that lack specific cofactors, specific inhibitors or low-sodium conditions. Molecular dynamics simulations based on these structural insights indicate that 2Fe-2S reduction in NqrD/E plays a crucial role in triggering Na^+^ translocation by driving structural rearrangements in the NqrD/E subunits, which subsequently influence NqrC and NqrF positioning. This study provides the first structural insights into the mechanism of Na^+^ translocation coupled to electron transfer in Na⁺-NQR.

## INTRODUCTION

The Na^+^-pumping NADH-quinone oxidoreductase (Na^+^-NQR) is the first enzyme in the respiratory chain of numerous marine and pathogenic bacteria, including *Vibrio cholerae*, *Neisseria gonorrhoeae,* and *Haemophilus influenzae*. Na^+^-NQR catalyzes the transfer of electrons from NADH to quinone, coupled to the translocation of Na^+^, generating an electrochemical Na^+^ gradient across the inner membrane. This gradient powers essential energy-dependent processes such as ATP synthesis, rotation of the flagellar motor, transport of nutrients, and operation of multi-drug transporters^1−3^. As originally revealed by biochemical and biophysical studies, Na^+^-NQR is an integral membrane complex composed of six subunits (NqrA-F) encoded by the *nqr* operon, with five spectroscopically visible redox cofactors (FAD, 2Fe-2S, 2FMNs, and riboflavin) and a total molecular mass or approximately 200 kDa Na^+^-NQR is exclusively found in prokaryotes and is structurally unrelated to the mitochondrial H^+^-pumping NADH-quinone oxidoreductase (respiratory complex I), making it a promising target for highly selective antibiotics^3,4^.

X-ray crystallographic^5,6^ and single-particle cryo-EM analyses^6,7^ have revealed the entire architecture of Na^+^-NQR (Fig. 1a). The cytoplasmic domain of the NqrF subunit includes a Rossmann nucleotide binding motif, where NADH binds, and contains one non-covalently bound FAD (FAD^NqrF^) that is the initial acceptor of electrons, and one 2Fe-2S cluster (2Fe-2S^NqrF^)^8,9^. The membrane-embedded NqrD and NqrE subunits form an inverted repeat structure that coordinates a second 2Fe-2S (2Fe-2S^NqrD/E^) ^2,10−12^, originally assigned as a single iron atom in the initial x-ray crystallographic model^7^ and subsequently identified as a 2Fe-2S cluster by cryo-EM, although it still lacks any spectroscopic or kinetic signature. The periplasmic NqrC has one covalently bound FMN (FMN^NqrC^) in its hydrophilic region^13,14^. The NqrB subunit is an integral membrane protein with a covalently bound FMN (FMN^NqrB^) and a unique riboflavin cofactor (RBF^NqrB^) that is the final donor of electrons to the ubiquinone (UQ) substrate^15,16^. The hydrophilic NqrA subunit is attached to the membrane through a tight interaction with the protruding part of the N-terminal region of NqrB. This cytoplasmic interfacial area has a significant influence on the binding of UQ^17,18^. These structural studies, along with earlier spectroscopic^19−23^, kinetic^10,24^, and mutagenetic investigations^8,9,12−14,25^, have established a consensus electron transfer pathway: NADH → FAD^NqrF^ → 2Fe-2S^NqrF^ → 2Fe-2S^NqrD/E^ → FMN^NqrC^ → FMN^NqrB^ → RBF^NqrB^ → UQ.

**Fig. 1:**
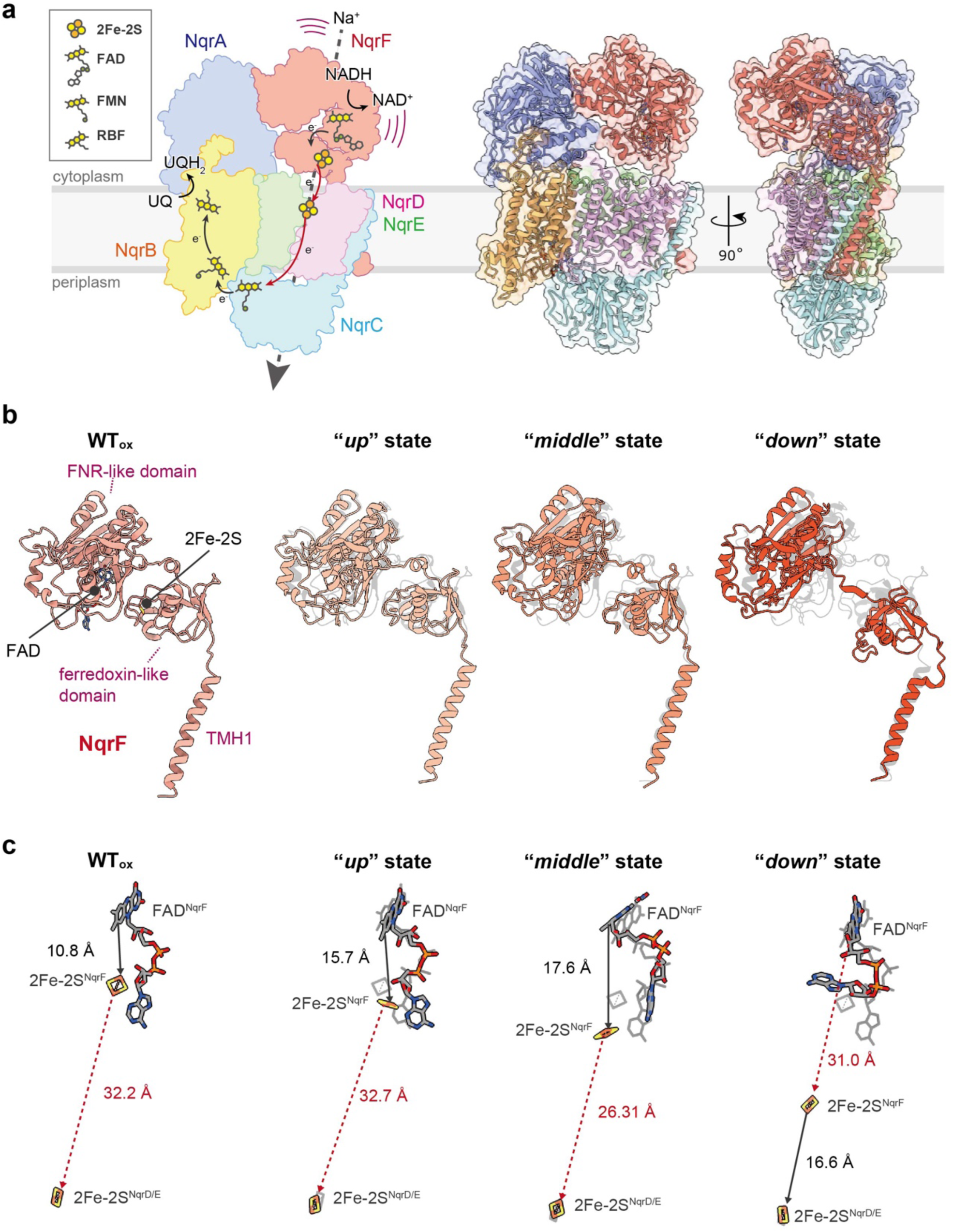
Architecture of Na^+^-NQR. ***a,*** Location of redox cofactors and electron transfer steps in the overall architecture of Na+-NQR (**WT_ox_**: pdb: 7XK3). *Left* hand panel: simplified structure of Na^+^-NQR showing the locations of redox cofactors and depicting electron transfer steps as arrows. Distances consistent with electron transfer are colored in *gray,* whereas long distances in this structural conformation, inconsistent with electron transfer (2Fe-2S^NqrF^ → 2Fe-2S^NqrD/E^, 2Fe-2S^NqrD/E^ → FMN^NqrC^), are colored in *red*. *Middle* and *right*-hand panels: the Na^+^-NQR assembly from front and side views, respectively. ***b,*** The “***up***”, “***middle***”, and “***down***” conformational states: The *left* panel shows the FNR domain containing the FAD cofactor, ferredoxin-like domain containing 2Fe-2S, and TMH1 of NqrF (side view). The remaining panels show the structure of NqrF in each conformational state illustrating the pivotal motion of NqrF in **WT_red_/ - Na^+^**. To compare **WT_ox_** and all states in **WT_red_/ - Na^+^**, the images of NqrF in **WT_ox_** are shown as a *gray silhouette*. ***c,*** Changes in distances between cofactors that accompany these conformational changes. The changes in distance between 2Fe-2S and FAD in NqrF, **WT_red_/ - Na^+^**. For comparison of **WT_ox_** with all states in **WT_red_/ - Na^+^**, the relative locations of **WT_ox_** are overlapped as *gray silhouette*. *Black continuous arrows* indicate feasible distances for electron transfer, and *red dashed arrows* indicate distances that are not consistent with electron transfer.

However, the distances between two of the donor-acceptor pairs in two of these electron transfer steps: 2Fe-2S^NqrF^ → 2Fe-2S^NqrD/E^, and 2Fe-2S^NqrD/E^ → FMN^NqrC^, shown as red arrows in Fig. 1a are too large (26-33 Å, edge-to-edge) for physiologically relevant electron transfer^7,25^. This indicates that the subunits containing these cofactors must undergo significant conformational rearrangements to reduce these spatial gaps during catalytic turnover^7,19^. Nevertheless, there is as yet no structural evidence to support these proposed rearrangements. In the case of the 2Fe-2S^NqrF^ → 2Fe-2S^NqrD/E^ electron transfer step, our previous cryo-EM study of wild-type Na^+^-NQR suggested that the cytosolic domain of the NqrF subunit is highly flexible, but the inferred range of motion is still not large enough to account for the electron transfer ^7^. In the case of 2Fe-2S^NqrD/E^ → FMN^NqrC^, an apparent discrepancy between X-ray and cryo-EM results may actually indicate a crucial conformational change. In the X-ray crystallographic structures of *V. cholerae* Na^+^-NQR^5,6^, the periplasmic domain of NqrC containing FMN^NqrC^ is positioned near the NqrD/E subunits, whereas the cryo-EM structure places this domain near NqrB. These results suggest that repositioning of the NqrC subunit during turnover could allow FMN^NqrC^ to shuttle electrons from 2Fe-2S^NqrD/E^ to FMN^NqrB^. However, there is no evidence to link this apparent rearrangement of NqrC to different intermediates in the catalytic mechanism.

Another enigma involves in the pathway(s) for Na^+^ translocation in the enzyme. Based on the biochemical characterization of conserved acidic residues within the TMHs, Juarez *et al.* identified critical acidic residues necessary for Na⁺ translocation (e.g. NqrB-D397, NqrD-D133, and NqrE-E95)^26^. Hau *et al.* observed two bound Na⁺ ions on the periplasmic side of NqrB in their cryo-EM map and proposed a Na⁺ channel within NqrB^6^. More recently, Kumar et al. reported a cryo-EM structure of the “Rhodobacter nitrogen fixing” protein (RNF) complex, an evolutionary ancestor of Na⁺-NQR^27^. Combined with atomic molecular dynamics (MD) simulations, they showed that the reduction of a 2Fe-2S cluster coordinated by the symmetric RnfA/E subunits (homologues of NqrD/E), induces capture of Na⁺ and subsequently triggers an *inward/outward* transition of the subunits. This model aligns with the alternating-access mechanism proposed by Bogachev and colleagues for Na^+^-NQR^10^. Despite numerous attempts to elucidate the mechanism of the redox-mediated Na⁺ translocation in Na⁺-NQR, no consensus has been reached. To address this, Na⁺-NQR structures of different reaction states (or different reaction conditions) are needed.

In this study, we have combined cryo-EM and atomic-level MD simulations to show how each subunit undergoes conformational rearrangements that facilitate electron transfer within the enzyme and how these rearrangements drive Na⁺ translocation. Our synergistic approach demonstrates that the reduction of the 2Fe-2S^NqrD/E^ triggers Na^+^ translocation across the membrane by coordinating with structural rearrangements in the NqrD/E subunits, which subsequently influence positioning of NqrC and NqrF. This study provides the first structural insights into the mechanism of Na^+^ translocation coupled to electron transfer in Na⁺-NQR.

## RESULTS

### Strategy for the preparation of cryo-EM grids

As described in the *Introduction*, structural studies on Na^+^-NQR to date suggest that redox driven Na^+^-translocation likely requires structural rearrangements in three different parts of the enzyme: the cytoplasmic domain of NqrF, the membrane embedded helices of NqrD/E, and the periplasmic region of NqrC, in order to modulate the distances between key pairs of redox cofactors: 2Fe-2S^NqrF^ to 2Fe-2S^NqrD/E^ and 2Fe-2S^NqrD/E^ to FMN^NqrC^ ^7^. To capture snapshots of Na^+^-NQR from *V. cholerae* in other putative conformational states, cryo-EM grids were prepared under nine different conditions that alter the catalytic cycle of the enzyme (Table 1): wild-type enzyme reduced by NADH in the presence (1) or absence (2) of Na^+^ (**WT_red_**, **WT_red_/-Na^+^**, respectively), (3) wild-type reduced by NADH in the presence of the inhibitor korormicin A (**WT_red_+KR**), (4) wild-type reduced by NADH in the presence of the inhibitor aurachin D-42 (**WT_red_+AD-42**), (5) NqrB-G141A reduced by NADH in the presence of korormicin A (**NqrB-G141A_red_+KR**), NqrC-T225Y *as isolated* (6) or reduced (7) by NADH (**NqrC-T225Y_ox_**, **NqrC-T225Y_red_**, respectively), and NqrB-T236Y *as isolated* (8) or reduced (9) by NADH (**NqrB-T236Y_ox_**, **NqrB-T236Y_red_**, respectively).

**Table 1:** The list of samples for cryo-EM analysis.

| **Sample name**<br>**(PDB ID)** | **variant** | **NADH** | **inhibitor** | **Na<sup>+</sup>** | **trait** | **TMH<sup>NqrCDEF</sup>**<br>**conformation** |
| --- | --- | --- | --- | --- | --- | --- |
| **WT<sub>red</sub> (9UUU)** | wild-type | + | – | + | – | Inward-opened |
| **WT<sub>red</sub>/ -Na<sup>+</sup>**<br>**“up” state (9UD6)**<br>**“middle” state (9UD8)**<br>**“down” state (9UD9)** | wild-type | + | – | – | – | Inward-opened<br>**(all state)** |
| **WT<sub>red</sub> + KR (9UD5)** | wild-type | + | KR | + | – | Inward-opened |
| **WT<sub>red</sub> + AD-42 (9UDG)** | wild-type | + | AD-42 | + | – | Inward-opened |
| **NqrB-G141A<sub>red</sub> + KR**<br>**“stable” state (9UDA)**<br>**“shifted” state (9UDF)** | NqrB-G141A | + | KR | + | KR<br>resistant | Inward-opened<br>**(“stable” state)**<br>Outward-opened<br>**(“shifted” state)** |
| **NqrB-T236Y<sub>ox</sub> (9UD3)** | NqrC-T225Y | – | – | + | FMN<sup>NqrB</sup><br>missing | Inward-opened |
| **NqrB-T236Y<sub>red</sub> (9UD4)** | NqrC-T225Y | + | – | + | FMN<sup>NqrB</sup><br>missing | Inward-opened |
| **NqrC-T225Y<sub>ox</sub> (9U5G)** | NqrC-T225Y | – | – | + | FMN<sup>NqrC</sup><br>missing | Outward-opened |
| **NqrC-T225Y<sub>red</sub> (9UD2)** | NqrC-T225Y | + | – | + | FMN<sup>NqrC</sup><br>missing | Outward-opened |

The NqrC-T225Y and NqrB-T236Y mutants lack FMN^NqrC^ and FMN^NqrB^, respectively, disrupting electron flow within the enzyme^19^. Korormicin A binds to the enzyme and allosterically inhibits the final electron transfer step in which ubiquinone becomes reduced^7,28^. The NqrB-G141A mutant exhibits significant resistance to korormicin A, though the inhibitor still binds to the enzyme^28^. These three mutations all interfere with the redox cycle of the enzyme and may thus be expected to influence the distributions in its conformational states. After the 3D reconstruction of particle images of the nine samples, each dataset was processed using the following analytical tools in the cryosparc soft^29^ toolbox to isolate intermediate states in molecular motion: 3D variability analysis (3DV)^30^, and 3DFlex (3DF)^31^ (Extended Data Fig. 1, Supplementary Fig. 1,2, and Supplementary Table 1).

### Structure of Na^+^-NQR reduced by NADH

We determined the structure of wild-type enzyme reduced by 5 mM NADH (**WT_red_**) at a resolution of 3.01 Å (Extended Data Fig. 1a and Supplementary Fig. 1a). As all the spectroscopically visible cofactors in Na^+^-NQR are reported to be fully reduced under these conditions^19^, this structure is thought to represent the fully reduced structure of the enzyme. The overall structure is very close to that of the enzyme *as isolated* (**WT_ox_,** PDB ID: 7XK3), with a single, well-resolved state observed for the NqrC subunit (Extended Data Fig. 2a). In the structure of **WT_red_**, NADH is bound in a long flat density around the Rossmann nucleotide binding motif of NqrF^8^ (Supplementary Fig. 3). Focusing on the membrane domain, the TMH bundle formed by NqrF, NqrD/E, and NqrC (TMH^NqrCDEF^ bundle) has a structure consistent with an *“inward-open”* state. Notably, TMHs 3-4 of NqrE on the cytoplasmic side have shifted slightly toward 2Fe-2S^NqrD/E^ compared to the **WT_ox_** structure (Extended Data Fig. 2a).

NqrF is a cytoplasmic subunit anchored to the membrane by a single transmembrane helix (TMH1^NqrF^). It is also tightly bound to the neighboring hydrophilic NqrA subunit through electrostatic interactions. The hydrophilic part of NqrF is highly flexible and comprises two distinct domains: a ferredoxin-<u>N</u>ADPH reductase (FNR)-like domain (K132‒G408) containing the FAD cofactor, and a ferredoxin-like domain (G33‒K131) harboring the 2Fe-2S^NqrF^ as its cofactor^8,9^ (Fig. 1b).

In the currently obtained cryo-EM structures, including that of fully reduced state (**WT_red_**), the NqrF subunit displays high conformational flexibility, adopting multiple distinct states. Nevertheless, the extent of flexibility does not extend into the membrane region as would be necessary for electron transfer. The distance between 2Fe-2S^NqrF^ and the 2Fe-2S^NqrD/E^, even at their closest proximity, is 30 Å (edge to edge), which is inconsistent with catalytic turnover rates. Therefore, during the catalytic cycle, NqrF likely undergoes a pivotal motion of its ferredoxin-like domain between FAD and 2Fe-2S^NqrD/E^, while the TMH1^NqrF^ remains firmly anchored to the membrane. However, the electron transfer in NqrF is too fast (below milliseconds) to capture the snapshot in motion^21^.

### Resolution of three conformations of the cytoplasmic NqrF subunit

To capture such rapid motions of NqrF, Na^+^-NQR was reduced by 5 mM NADH in the absence of Na^+^ (**WT_red_/-Na^+^**). In these conditions, the rate constant for FMN^NqrC^ reduction is slowed approximately eightfold^19^, which may allow the identification of previously unobserved conformational states of NqrF. By focusing on the position of the ferredoxin-like domain, three distinct states of NqrF–***up*, *middle***, and ***dow*n**–were identified as separate classes (Fig. 1b,c, Extended Data Fig. 1b, and Supplementary Fig. 1b). These continuous density maps were further refined using 3DF (Supplementary Fig. 2a).

The *“**up**”* state comprises over 87% of the particle population in the dataset, providing a high-resolution map at 2.65 Å, where the FNR-like and ferredoxin-like domains are found in similar positions to those in the enzyme *as isolated* (**WT_ox_**). The *“**down**”* state, reconstructed from the smallest population of the particles (5%), gave a final density map at 3.11 Å. Remarkably, in the “***down***” state, the ferredoxin-like domain fits into the pocket formed by the NqrD/E subunits, whereas this pocket is exposed to the cytoplasm in the other two states. Consequently, the distance between 2Fe-2S^NqrF^ and 2Fe-2S^NqrD/E^ is reduced to 16.6 Å, consistent with electron transfer between them, while the distance between FAD^NqrF^ and 2Fe-2S^NqrF^ is increased to 31.0 Å (Fig. 1c). Although the resolution of the *“**middle**”* state is relatively low (3.75 Å), the density map is sufficient to position the ferredoxin-like domain in an intermediate location between the *“**down**”* and *“**up**”* states (Fig. 1b,c).

Notably, in all three states, the structures of the membrane domains, including TMH^NqrF^, remain almost identical to those in **WT_red_**, with the TMH^NqrCDEF^ bundle in an *“inward-open”* state (Fig. 2 and Extended Data Fig. 2b,c). This shows that the reduction of the NqrF cofactors allows the ferredoxin-like domain harboring 2Fe-2S^NqrF^ to bridge transport of electrons from FAD^NqrF^ to 2Fe-2S^NqrD/E^, without altering the conformation of the TMH^NqrCDEF^ bundle.

**Fig 2:**
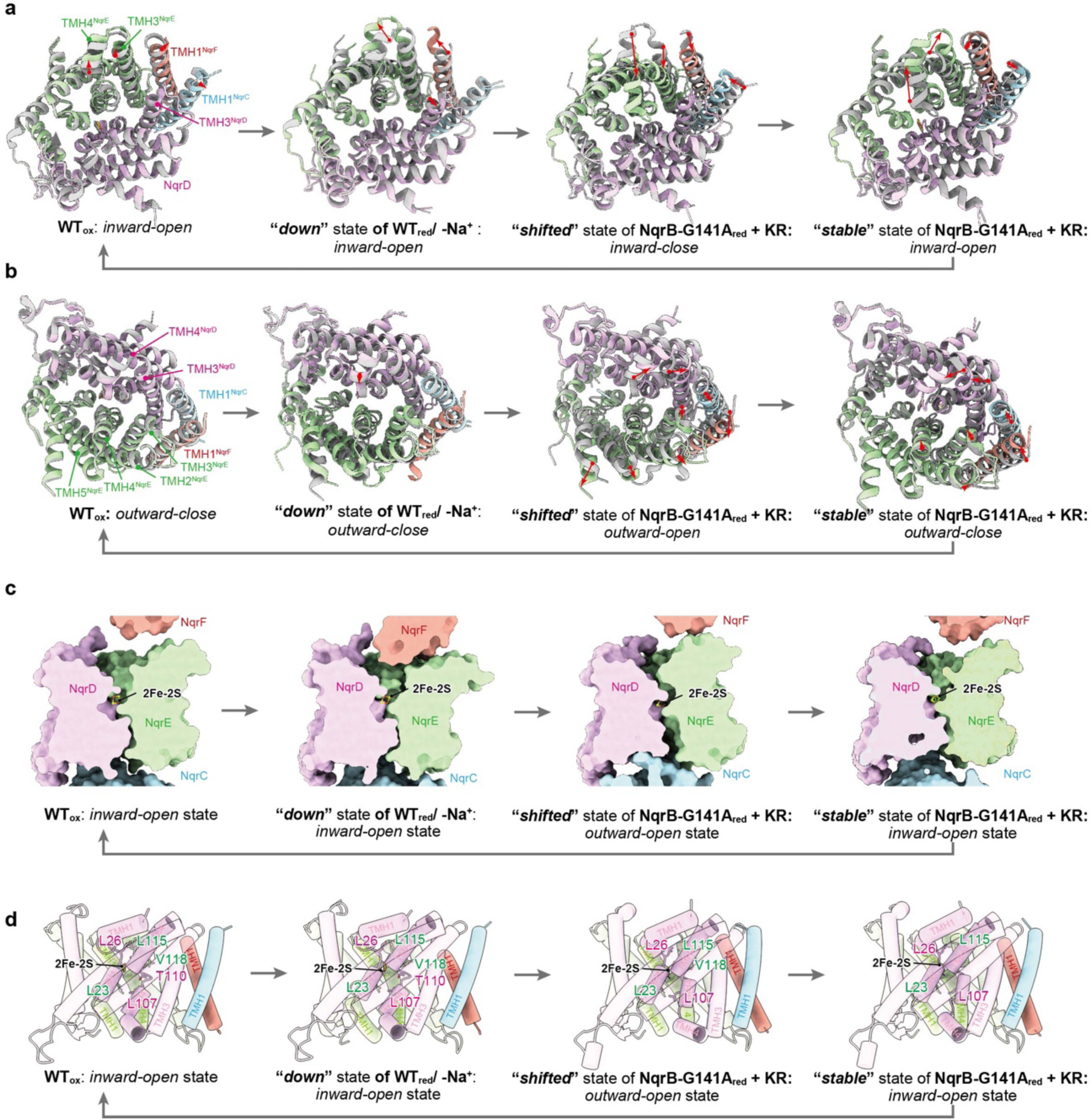
Conformations of the transmembrane helices of NqrCDEF as the catalytic cycle progresses from four different views. Four steps in the catalytic cycle are represented by the following structures: **WT_ox_**, “*down*” state of **WT_red_/ - Na^+^**, “*shifted*” state, and “*stable*” state of **NqrB-G141A_red_ + KR**. *Gray* images show the structure of the previous state; red arrows show the direction of movement from the previous state to the current state. *a,* Cytoplasmic view. *b,* Periplasmic view. *c,* Sectional side views of NqrCDEF. *d,* Tube models from front view. The critical amino acid residues to coordinate Na^+^ in the inward and outward gates in the MD simulation are shown as stick models. Supplementary Videos 2, 3, and 4 show how the arrangement of the helices morphs during the catalytic cycle from the complete view of the enzyme, cytoplasmic, and periplasmic views.

### The conformational shift of the NqrC subunit between NqrB and NqrD/E

Korormicin A (KR) and aurachin D-42 (AD-42) are specific inhibitors of Na⁺-NQR that can serve as tools for capturing key reaction intermediates of the enzyme^32−37^. However, wild-type enzyme with korormicin A or aurachin D-42 bound (**WT_red_ + KR**, **WT_red_ + AD-42**), treated with NADH, did not show any distinct reaction intermediates (Extended Data Fig 1c,d and 2h,i and Supplementary Fig 1c,d, and 4), as the inhibitor completely terminates UQ reduction at the final step of the catalytic cycle, mimicking **WT_red_**. To decelerate the catalytic cycle of Na^+^-NQR, we prepared a cryo-EM grid of the NADH-treated NqrB-G141A mutant with korormicin A bound (**NqrB-G141A_red_ + KR**). This alanine substitution excludes the alkyl side chain of korormicin A from the binding cavity (Supplementary Fig 5), resulting in a decrease in the binding affinity (∼160-fold increase in IC_50_ values) although the inhibitor molecule still binds to the mutated enzyme^28,38−40^. This unique combination of mutation and inhibitor could potentially influence the catalytic cycle of the enzyme, and lead to the identification of multiple conformational states of the enzyme.

The image particles were processed through iterative 3DV with a mask covering the entire Na⁺-NQR complex, followed by focused refinement on the NqrC density. This approach identified two distinct states of Na^+^-NQR, one in which the hydrophilic domain of the NqrC subunit approaches the NqrD/E and one in which it approaches NqrB, likely reflecting its motion during catalytic turnover (Extended Data Fig. 1e, Supplementary Fig. 1e, Supplementary Video 1). These continuous density maps were further refined by 3DF (Supplementary Fig. 2b) to reconstitute two conformational states: *“**stable**”* and “***shifted***” with particle populations of 88 and 12%, respectively (Fig. 3).

**Fig. 3:**
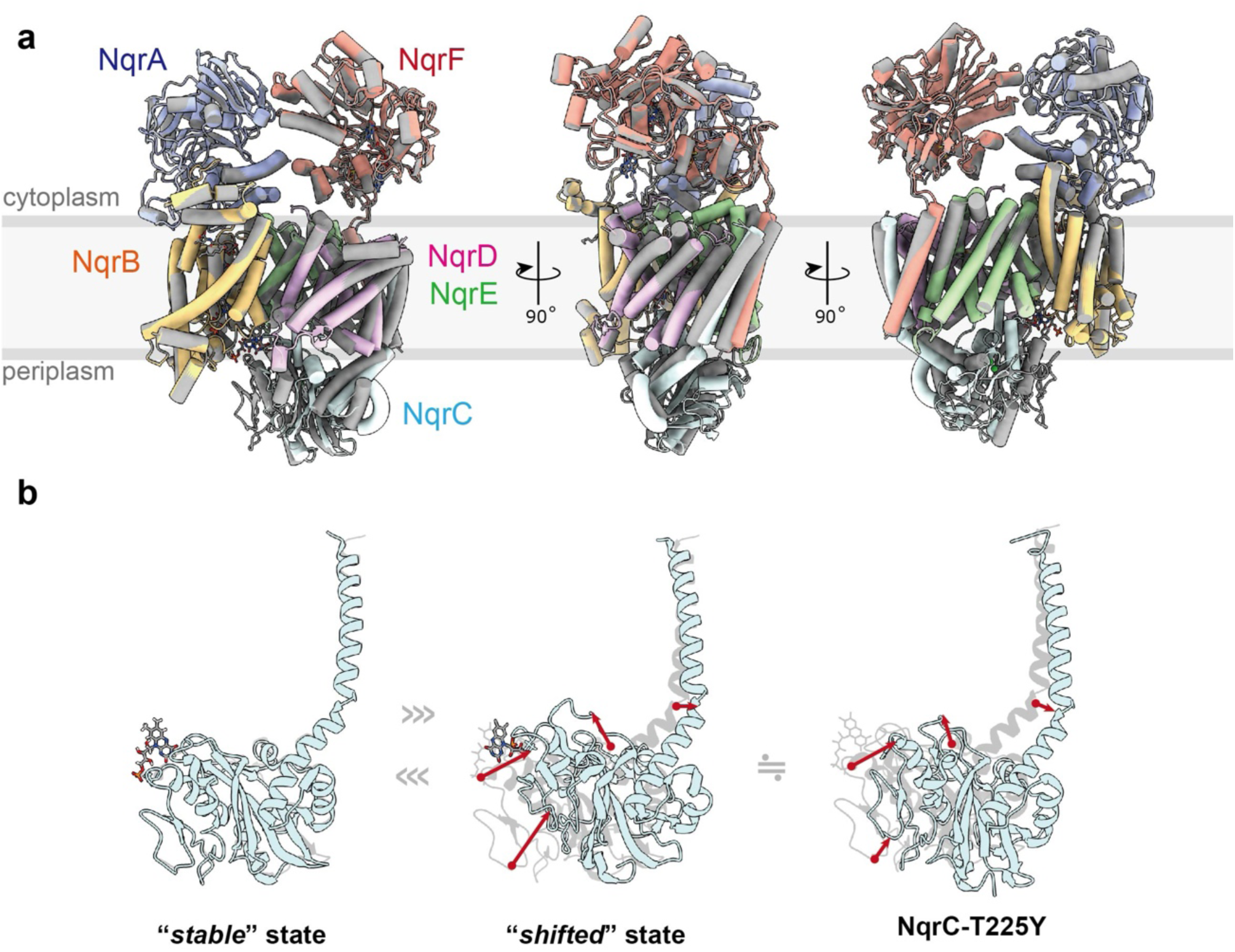
The shift of NqrC: Transition of NqrC from “*stable*” to “*shifted*” conformation. ***a,*** Comparison of two states in **NqrB-G141A_red_ + KR** sample, complete structure from three different views; “***stable***” state (*gray*) and “***shifted***” state (*color*). In the “***stable***” state (88% of particles) the hydrophilic region of NqrC is located close to NqrB; in the “***shifted***” state (12% of particles) NqrC is close to NqrD (Supplementary Fig. 1e). ***b,*** NqrC is shown alone. *Left*, “***stable***” state from **NqrB-G141A_red_ + KR**; center, “***shifted***” state from **NqrB-G141A_red_ + KR**; *right*, NqrC from **NqrC-T225Y_red_**. For comparison, the *gray silhouettes* in the center and right-hand images show the “***stable***” state.

In the “***stable***” state (Fig. 3b), resolved at 2.93 Å, the NqrC subunit has moved closer to NqrB, consistent with rapid electron transfer between FMN^NqrC^ and FMN^NqrB^ (7.8 Å). This structure aligns with that of the enzyme treated with NADH (**WT_red_**). In contrast, the “***shifted***” state, reconstructed at 2.61 Å with improved local alignment of particle images, exhibits a weaker density for the hydrophilic domain of NqrC, suggesting increased flexibility in this region. This flexibility makes full *de novo* modeling challenging; however, a structural model with rigid body fitting indicates that the hydrophilic domain of NqrC subunit has shifted toward the NqrD/E subunits.

In the ***shifted*** state (Fig. 3b), while FMN^NqrC^ remains distant from 2Fe-2S^NqrD/E^ (∼27 Å), the loop of Glu169-Gly177^NqrC^, which includes Thr173 and Leu176 that stabilize the isoalloxazine ring of this FMN^9^, is embedded in the pocket formed by the NqrD/E subunits. The existence of these two distinct conformations shows that the periplasmic domain of NqrC, which houses FMN^NqrC^, switches its position between NqrB and NqrD/E in a way that allows it to mediate electron transfer from 2Fe-2S^NqrD/E^ to FMN^NqrB^ (Fig. 1a, 3). Notably, the TMH^NqrCDEF^ bundle adopts the *“inward-open”* conformation in the “***stable****”* state but transitions to the *“outward-open”* conformation in the “***shifted****”* state (Fig. 2). This *inward-to-outward* transition of the bundle appears to be linked to the relocation of the periplasmic domain of NqrC from the NqrB side to the NqrD/E side.

### The rearrangement of NqrC is coordinated with the inward/outward transition of the TMH^NqrCDEF^ bundle

To understand the factors that could cause NqrC to move from the NqrB side to the NqrD/E side, we analyzed the structure of mutants lacking either FMN^NqrB^ (NqrB-T236Y) or FMN^NqrC^ (NqrC-T225Y). In the mutant lacking FMN^NqrB^ (Extended Data Fig 3a)^34−36^, both “*as isolated*” and “NADH-treated” enzymes (**NqrB-T236Y_ox_** and **NqrB-T236Y_red_**) showed *“inward open”* structures with the NqrC subunits positioned near NqrB at resolutions of 3.83- and 3.31-Å, respectively (Extended Data Fig. 1f,g and 2d,e and Supplementary Fig 1f,g). These structures closely resemble **WT_ox_** and **WT_red_** (Supplementary Fig. 6), suggesting that this mutant adopts a similar conformation in which NqrC has shifted fully from the NqrD/E side to the NqrB side. In contrast, the mutant lacking FMN^NqrC^ (Extended Data Fig. 3b) show identical *outward-open* structures with NqrC shifted toward NqrD/E, irrespective of NADH treatment (**NqrC-T225Y_ox_** and **NqrC-T225Y_red_**) with resolutions of 2.66- and 2.59-Å, respectively (Extended Data Fig 1h,i and 2f,g, Supplementary Fig. 1h,i). In these conformations, as in the “***shifted***” state in **NqrB-G141A_red_ + KR**, the periplasmic domain of NqrC is positioned closest to the dimetric NqrD/E subunits (Fig. 3b and Extended Data Fig 3c).

These findings, including the structure of the korormicin A-bound NqrB-G141A mutant (**NqrB-G141A_red_ + KR**), suggest that the rearrangement of NqrC is coupled to the *inward-to-outward* transition of the NqrD/E subunits, which bind 2Fe-2S^NqrD/E^. In the structure of Na⁺-NQR reduced by NADH in the absence of Na⁺ (**WT_red_/-Na^+^**), all observed NqrF states (***up****, **middle**,* and ***down***) corresponded to the *inward-open* state of the TMH^NqrCDEF^ bundle (Fig. 2, Extended Data Fig. 2a-c), suggesting that the position of 2Fe-2S^NqrF^ does not influence the membrane domain structure. Considering that electron transfer from 2Fe-2S^NqrD/E^ to FMN^NqrC^ requires conformational rearrangements of both NqrC and NqrD/E, it is plausible that, the redox state of 2Fe-2S^NqrD/E^ may drive the *inward-to-outward* transition of the TMH^NqrCDEF^ bundle. This transition likely triggers the relocation of the periplasmic domain of NqrC, thereby facilitating the reduction of FMN^NqrC^.

### Shift in local resolution map reflects redox driven conformational changes of Na^+^-NQR

During single-particle analysis, we noticed that some of the redox-induced rearrangements of Na^+^-NQR subunits correlate with shifts in the local resolution map. A notable example of this is the rearrangement of NqrF during electron transfer from FAD to 2Fe-2S^NqrD/E^ (Fig. 1b,c). In the “***down***” and “***middle***” states, where 2Fe-2S^NqrF^ is positioned close enough to 2Fe-2S^NqrD/E^ to allow facile electron transfer, the ferredoxin-like domain of NqrF shows lower resolution, while the FNR-like domain containing FAD remains highly resolved (Supplementary Fig. 2a). This suggests that reduction of 2Fe-2S^NqrF^ increases the flexibility of the ferredoxin-like domain, enabling its movement towards 2Fe-2S^NqrD/E^, while the FNR-like domain remains stable, preventing electron backflow.

Interestingly, this observation is even more pronounced in the structure of the NqrB-G141A mutant with bound korormicin A (**NqrB-G141A_red_ + KR**), which exhibits two distinct states of the TMH^NqrCDEF^ bundle. In the “***stable***” state, where the bundle adopts an “*inward-open*” conformation, the cytoplasmic domain of NqrF appears blurred (flexible), whereas the periplasmic domain of NqrC exhibits sharp density (rigid). Conversely, in the “***shifted***” state, where the bundle adopts an “*outward-open*” conformation, NqrF displays sharp density (rigid), while NqrC appears blurred (flexible) (Extended Data Fig. 1e, Supplementary Fig 1e, and 2b).

A similar pattern appears in the NqrC-T225Y mutant, which lacks FMN^NqrC^. This mutant adopts the “*outward-open*” conformation of its TMH^NqrCDEF^ bundle, similar to the “***shifted***” state, and shows the highest resolution for NqrF, indicating that it remains in a single, stable “***up***” conformation (Extended Data Fig 1h,i, Supplementary Fig 1h,i). Together, these results suggest that the “*inward-to-outward*” transition alternates the flexibility of NqrF and NqrC, coordinating electron transfer from 2Fe-2S^NqrF^ to FMN^NqrC^ via 2Fe-2S^NqrD/E^. The shifting resolution patterns highlight how structural flexibility and stability work together to control electron flow.

### A proposed mechanism for Na^+^ translocation

Recently, Müller and coworkers published a cryo-EM structure of the RNF complex from *A. woodii*, where the RnfA/E subunits coordinating a 2Fe-2S cluster show high homology to the NqrD/E subunits of Na⁺-NQR^27^. Long-range molecular dynamics (MD) simulations predicted that the dimeric RnfA/E subunits can adopt two distinct conformations—*inward-* and *outward-open*—depending on the redox state of the 2Fe-2S cluster. This rearrangement appears to be crucial for both Na⁺ translocation and electron transfer within the enzyme. Their report supports the hypothesis that, in Na⁺-NQR, the redox state of the 2Fe-2S^NqrD/E^ not only drives the *inward-to-outward* transition of TMHs^NqrCDEF^ bundle but also facilitates Na⁺ translocation.

We have performed MD simulations beginning from the cryo-EM structure with an *inward-open* state of TMHs^NqrCDEF^ bundle, which provide some insight into the mechanism of Na^+^ uptake from the cytoplasm (Supplementary Table 2). Spatial density analysis of MD simulations results showed that Na^+^ ions and water can access the cytoplasmic pocket near 2Fe-2S^NqrD/E^ in the wild-type Na^+^-NQR with reduced 2Fe-2S^NqrF^ and 2Fe-2S^NqrD/E^ (Fig. 4a,b). The reduction of 2Fe-2S^NqrD/E^ is linked to formation of a Na^+^ binding site on the cytoplasmic side. In contrast, Na^+^ ions could not fully access the cytoplasmic pocket with the oxidized 2Fe-2S^NqrD/E^ (Supplementary Fig 7). Pairs of hydrophobic residues, that could function as gates for cation movement, are found above and below the Na^+^ binding site: Leu26^NqrD^-Leu115^NqrE^ and Leu107^NqrD^-Leu23^NqrE^ (Fig. 4f). These residues control the access of Na^+^ ions and water and are conserved between NQR and RNF (Supplementary Fig 8).

**Fig 4:**
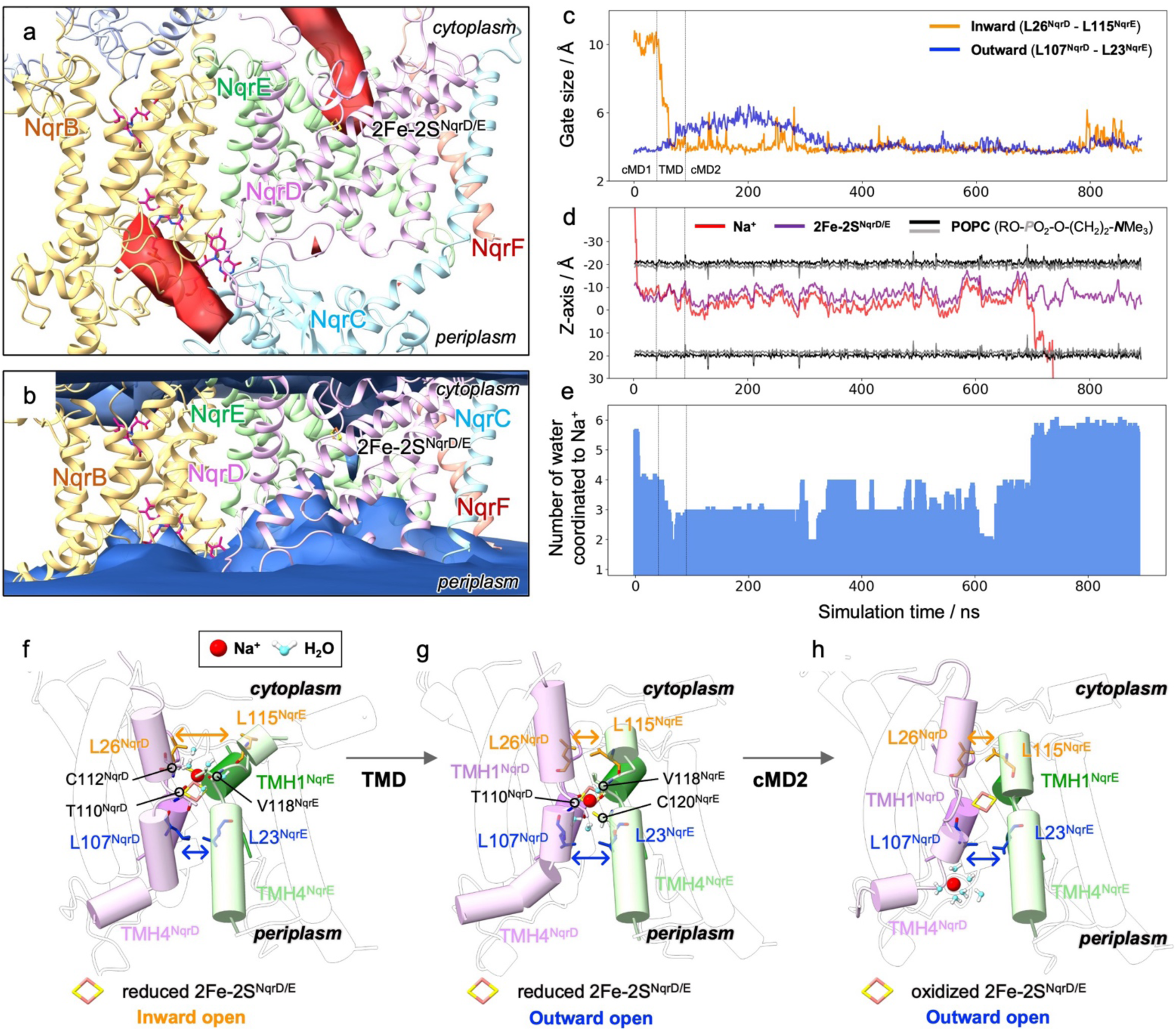
Na^+^ translocation pathway from MD simulation. ***a-b,*** The accessibilities of Na^+^ and water are shown in the structure of wild-type of Na^+^-NQR with reduced 2Fe-2S^NqrF^ and 2Fe-2S^NqrD/E^. ***a,*** The spatial density of Na^+^ is shown in *red*; the relative iso-density surface of 3.5 to the bulk value is shown. ***b,*** The spatial density of water is shown in *cyan*; the relative iso-density surface of 0.3 to the bulk value is shown. ***c-e,*** A first conventional MD (cMD1) using the inward-open model containing reduced 2Fe-2S^NqrF^ and 2Fe-2S^NqrD/E^ based on **WT_ox_** structure is followed by the targeted MD (TMD) inducing the *inward-to-outward* transition of TMHs bundle and the second conventional MD (cMD2) containing oxidized 2Fe-2Ss and reduced FMN^NqrC^. ***c,*** The inward gate and outward gate size are defined by the distances between L26^NqrD^ and L115^NqrE^ (*red*), and between L107^NqrD^ and L23^NqrE^ (*blue*), respectively. The transitions of these gate sizes are shown during the simulation. ***d,*** The z coordinates (perpendicular to the membrane surface) of the Na^+^ being translocated and the 2Fe-2S^NqrD/E^ are shown together with the positions of the membrane positions. ***e,*** Number of water molecules coordinated to the Na^+^ as a function of time in the simulation. ***f-g,*** Representative snapshots during Na^+^ translocation in MD simulation. The amino acid residues that coordinate Na^+^ ion in the inward and outward gates are indicated in *red* and *blue* stick models, respectively. Na^+^ bound in the inward-open and outward-open conformations are shown in ***f*** and ***g***, respectively. The released of Na^+^ to the periplasm is shown in ***h***.

Because the spontaneous *inward-to-outward* transition of the TMH^NqrCDEF^ bundle was not observed within the time scale of simulations, we employed targeted MD (TMD) simulation to induce this conformational transition^41^. During the TMD simulation, an external force was applied to make the conformation of NqrD/E move toward the *outward-open* state. During the TMD and subsequent equilibrium simulations, the inside gate closed while the outside gate opened (Fig. 4c,f,g, and Extended Data Fig. 4). The conformational transition shifted the bound Na^+^ to the outward side of the binding site relative to 2Fe-2S^NqrD/E^ (Fig. 4d,g). With the oxidation of 2Fe-2S^NqrD/E^, the bound Na^+^ was eventually released to the outer side of the membrane (Fig. 4h). Na^+^ translocation was observed in two additional independent trajectories following the TMD simulation (Supplementary Fig. 9). It is noteworthy that the Na^+^ ion was partially hydrated during the translocation with two or more coordinated water molecules (Fig. 4e). This may be due to the lack of acidic residues in the binding site that would strongly bind the ion and dehydrate it as in some transporter proteins^42^. Instead, the binding site is made up of backbone atoms of Thr110^NqrD^, Cys112^NqrD^, Val118^NqrE^, and Cys120^NqrE^ (Fig. 4f−h and Extended Data Fig 5).

## DISCUSSION

### The overall reaction mechanism of Na^+^-NQR

Since the first publication of a structural model of Na^+^-NQR, it has been clear from the spatial arrangement of the redox cofactors that structural rearrangements would have to play a critical role in the enzyme and that understanding these conformational changes would be essential to elucidating the mechanism of redox-driven Na^+^ pumping. In this study, we have identified at least five distinct conformational states of Na^+^-NQR, which may reflect key reaction steps of the enzyme. Based on the multiple conformations of Na^+^-NQR revealed by the current cryo-EM work and previous studies^9^, together with MD simulations, we propose an overall reaction mechanism for the enzyme, which progresses through six key steps (Fig. 5 and Supplementary Fig. 10).

***Step 1 (Oxidized state):*** In the absence of NADH, all cofactors—except for RBF, which is present as a stable neutral semiquinone^15^—are fully oxidized. In this form, the dimeric NqrD/E subunits are in an *inward-open* state. The hydrophilic domain of NqrF remains flexible and does not adopt a distinct structure; however, the range of motion of NqrF does not bring it into proximity with the NqrD/E subunits.
***Step 2 (NADH binding):*** When NADH binds to the NqrF subunit, it donates two electrons. One electron is transferred to the 2Fe-2S^NqrF^, increasing the flexibility of the ferredoxin-like domain of NqrF, allowing it to approach NqrD/E.
***Step 3 (Na^+^ uptake):*** The highly flexible ferredoxin-like domain of NqrF approaches the cytoplasmic pocket formed by the dimeric NqrD/E. This rearrangement allows electron transfer from 2Fe-2S^NqrF^ to 2Fe-2S^NqrD/E^. The negative charge on the reduced 2Fe-2S^NqrD/E^ enables accommodation of hydrated Na^+^ between inward and outward gates, which are made up of conserved hydrophobic amino acids.
***Step 4 (Transition to outward-open conformation and Na^+^ release):*** The NqrD/E subunits, upon reduction of 2Fe-2S^NqrD/E^ and binding of Na^+^, undergo a conformational switch to the *outward-open* state, exposing the bound Na^+^ to the periplasmic side. This transition increases the flexibility of hydrophilic domain of NqrC, enabling its approach toward NqrD/E, while the flexibility of NqrF decreases, preventing electron backflow into 2Fe-2S^NqrF^. This Na^+^- and redox-dependent repositioning of NqrC allows electron transfer from 2Fe-2S^NqrD/E^ to FMN^NqrC^. Simultaneously, the bound Na⁺ loses affinity and is released to the periplasm following reoxidation of 2Fe-2S^NqrD/E^.
***Step 5 (return to inward-open conformation):*** This reoxidation also induces a conformational relaxation of the NqrD/E subunits, restoring them to the *inward-open* state and allowing NqrC to return to the NqrB side. This movement facilitates electron transfer from FMN^NqrC^ to RBF^NqrB^ via FMN^NqrB^. Since i) the UQ reduction is a two-electron process while ii) FMNs and RBF undergo single-electron redox reaction^20^, a similar series of reactions (from Steps 2 to 5) repeats to the transfer of a second electron, which is coupled with the translocation of second Na^+^.
***Step 6 (UQ reduction):*** Through two sequential redox cycles from 2Fe-2S^NqrF^ to RBF^NqrB^, UQ accepts two electrons to form UQH_2_, which is then released from the enzyme. Thus, the enzyme returns to the initial oxidized state. Although the structure of Na⁺-NQR with bound UQ has not yet been identified, photoaffinity labeling experiments suggest that the UQ head-ring is positioned within a pocket in the NqrA subunit, approximately 20 Å above the membrane surface. Note that Hau et al. reported a structure of UQ_2_-bound Na^+^-NQR, in which UQ_2_ was modeled inside the inhibitor binding pocket in the *N*-terminal region of NqrB^6^. However, the density map of UQ_2_ molecule in the pocket is too weak to definitely model UQ_2_ molecule. It is, therefore, premature to conclude that UQs accept electrons from RBF in the inhibitor binding pocket in NqrB.

**Fig. 5:**
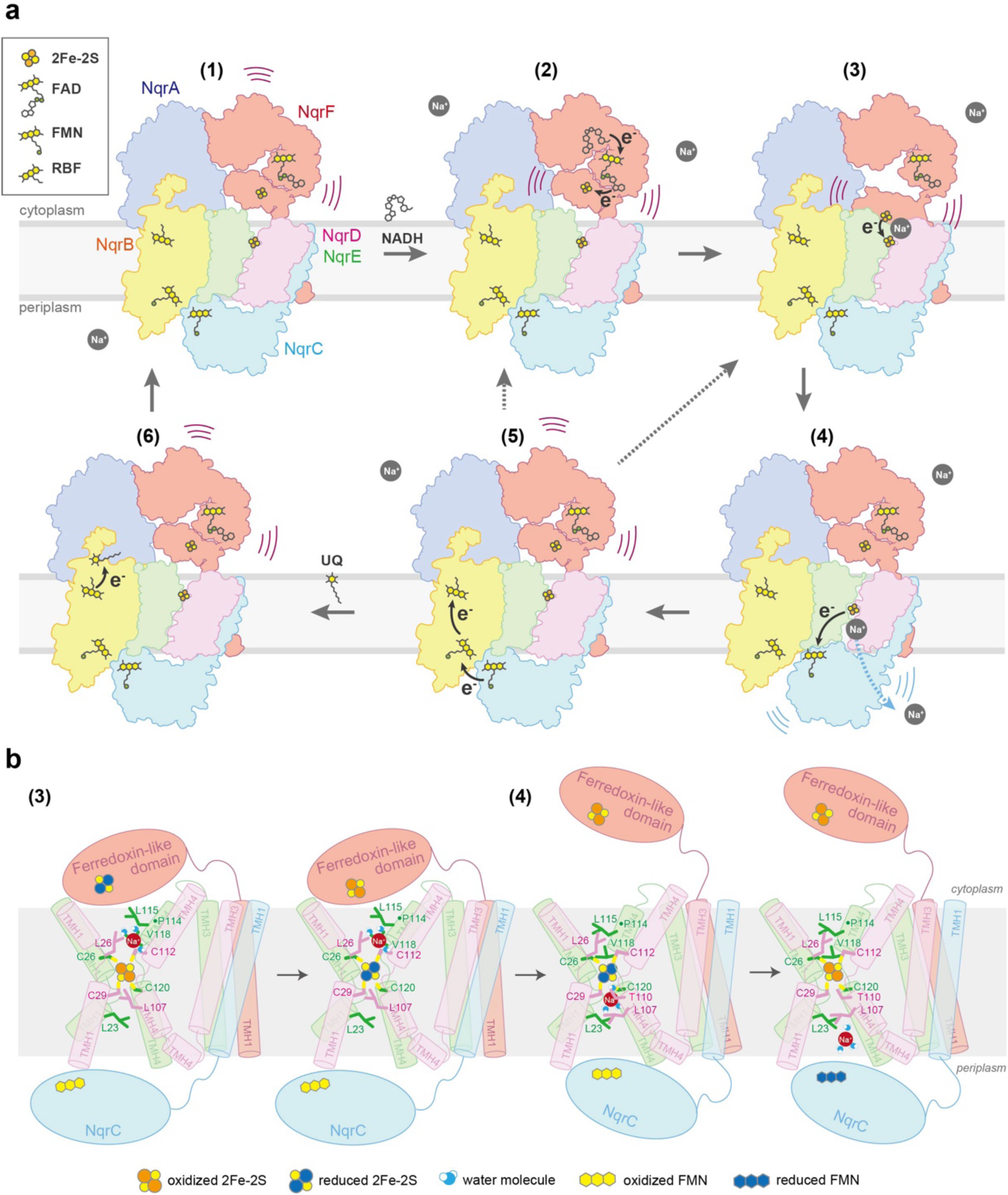
The model of conformational changes. ***a,*** The cascade of conformational changes. One electron flow is shown as *black arrows*. Step 1: oxidized state, step 2: NADH binding, step 3: Na^+^ uptake, Step 4: transition to outward-open conformation, step 5: Na^+^ release and return to inward-open conformation, Step 6: UQ reduction. Since UQ requires a two-electron reduction, step 5 or 6 need to go back step 2 or 3. ***b,*** The detailed model of essential TMHs for conformational changes in step 3-4.

### Na^+^-translocating pathway

The mechanism of redox-driven Na⁺ translocation by Na⁺-NQR remains to be fully elucidated. Based on the X-ray crystallographic and cryo-EM models, Fritz and colleagues proposed that cytoplasmic Na⁺ is translocated to the periplasm via a pathway within NqrB, where it is coordinated by residues such as Ile371, Arg372, Pro376, and Tyr378^5,6^. Our MD simulations also identified Na⁺ binding to periplasmic residues of NqrB, such as Thr236 and Ser239 (Fig. 4a,b, Extended Data Fig 5, Supplementary Fig. 7 and 11 and Supplementary Table 3). However, this Na^+^ was consistently observed across all enzyme states identified in this study, indicating that NqrB is unlikely to be directly involved in redox-driven Na⁺ translocation across the membrane. We were unable to identify any continuous ion translocation pathway spanning the transmembrane region of NqrB, but we have found a pathway of this kind at the interface between the NqrD/E subunits. Consistent with the recent study on the *A. woodii* RNF complex^27^, our findings strongly suggest that Na^+^ translocation is mediated exclusively by the NqrD/E subunits, rather than by NqrB.

The redox-dependent *inward-to-outward* transition of the TMH^NqrCDEF^ bundle was not observed in our MD simulations. During structural modeling of Na^+^-NQR in the inward- and outward-open states, we noticed that the conformation of the NqrD/E subunits appeares to undergo a conformational change centered on Prolines in TMH4 of NqrE—a highly conserved residue located near the 2Fe-2S^NqrD/E^ and surrounded by hydrophobic residues (Fig. 5b, and Supplementary Fig. 8). A change in the redox state of 2Fe-2S^NqrD/E^ may be linked to subtle local rearrangement in adjacent amino acid residues by through electrostatic interactions, and these rearrangements could be amplified by conserved proline, leading to larger conformational shifts. A similar proline-mediated switching mechanism has been proposed in the structure of voltage-gated potassium (Kv) chanel^43^.

Finally, we address how Na⁺ is translocated against the sodium motive force (Na⁺ gradient across the membrane). Our findings indicate that the “*inward-to-outward*” transition of the TMH^NqrCDEF^ bundle alternately regulates the flexibility of the NqrF and NqrC subunits to control electron flow. This mechanism likely essential for the redox-driven Na⁺ translocation—namely, that electron does not flow forward without Na⁺ being expelled to the periplasm. For instance, in the *outward-open* conformation with periplasmic Na⁺ bound, electron backflow from 2Fe-2S^NqrD/E^ to 2Fe-2S^NqrF^ would be necessary to trigger the transition to the *inward-open* conformation, that would allow the bound Na⁺ to move back toward the cytoplasm. Nevertheless, as NqrF is firmly stabilized in this state, such electron backflow may not take place.

In summary, the present structural analyses of Na⁺-NQR in multiple distinct conformations, combined with MD simulations, provide critical insights into the coupling mechanism between the electron transfer and Na^+^ translocation. These findings illustrate how structural rearrangements of individual subunits can control Na⁺ translocation and electron transfer, to achieve coupling between the two processes. This study establishes a valuable framework for further mechanistic studies of Na⁺-NQR and antibiotics development targeting the enzyme.

## METHODS

### Na^+^-NQR expression and purification

Expression of *Vibrio cholerae* Na^+^-NQR from a plasmid, in the parent organism, construction of site-specific mutants, and cultivation of *V. cholerae* strains have been described previously (REF)^7^. Na^+^-NQR was purified as previously described. The membrane pellets were solubilized in a buffer containing 50mM NaPi, 300mM NaCl, 5.0mM imidazole, and protease inhibitor cocktail (Sigma-Aldrich), 0.4% (w/v) *n*-dodecyl-*β*-D-maltoside (DDM), pH 8.0, with stirring for 1 h at 4 °C. The mixture was clarified by ultracentrifugation (100,000 g), and the supernatant was mixed with 20 mL of Ni-NTA resin (Qiagen) that had been equilibrated with binding buffer (50mM NaPi, 300mM NaCl, 5.0mM imidazole, and 0.05% DDM). After gentle agitation for 1 h at 4 °C, the suspension was poured into an empty plastic column. The column was washed with 4 volumes of the binding buffer, followed by 5 volumes of a wash buffer (same as binding buffer but containing 10mM imidazole). The protein was eluted from the column using an elution buffer (same as binding buffer but containing 100 mM imidazole). The resulting Na^+^-NQR preparation was applied to a DEAE anion exchange column (HiPrep DEAE FF 16/10, connected to an ÄKTA system, Cytiva) equilibrated with buffer A [50 mM Tris/HCl, 1.0 mM EDTA, 5% (v/v) glycerol, and 0.05% (w/v) DDM, pH 8.0]. The protein was eluted with a linear gradient of buffer B (buffer A containing 2.0 M NaCl), and the fractions containing pure Na^+^-NQR were pooled and concentrated to 15∼20 mg of protein/mL. Subunit assembly was confirmed by SDS-PAGE^44^ and enzyme activity was checked by the inhibitors-sensitive NADH-UQ_1_ oxidoreductase activity measurements at 282 nm (ε = 14.5mM^−1^cm^−1^)^2^. To remove glycerol from samples, the target protein was further separated by size-exclusion chromatography (Superose^TM^ 6 increase 10/300 gel filtration column, Cytiva) with an elution buffer containing 50 mM Tris-HCl, 100 mM NaCl, 1.0 mM EDTA, 0.05% DDM (pH 8.0).

### Cryo-EM acquisition

To reduced DDM concentrations, immediately prior to cryo-EM grid preparation, all of the purified Na^+^-NQR samples were washed with the same elution buffer without detergent (50 mM Tris-HCl, 100 mM NaCl, 1.0 mM EDTA, pH 8.0). To obtain the reduced form of Na^+^-NQR, Na^+^-NQR solution was mixed with an NADH solution to final concentration of 5 mM just before application onto a grid. 2.7 µL of Na^+^-NQR solution at the concentration of 10 mg/mL was applied onto a glow-charged Quantifoil Cu R1.2/1.3. The grid was blotted by Vitrobot IV and vitrified with liquid ethane. Cryo-EM movies were acquired with a Titan Krios electron microscope (Thermo Fisher, USA) equipped with K3 BioQuantum camera (Gatan, United States). Data collection was automatically controlled by SerialEM software at a nominal magnification of 81,000 and the pixel size is 0.88 Å/pix. The total electron dose and frame rate were 60 electron/Å^2^ and 0.1 s, respectively.

### Cryo-EM data processing

The details of image processing steps in each condition are shown in Extended Data Fig. 1. All of seven datasets (**WT_red_**, **WT_red_/ -Na^+^**, **WT_red_ + KR**, **WT_red_ + AD-42**, **NqrB-G141A_red_ + KR**, **NqrB-T236Y_ox_**, **NqrB-T236Y_red_**, **NqrC-T225Y_ox_**, and **NqrC-T225Y_red_**) were processed by cryoSPARC v3.3.1 or v4.0. 6,086, 13,302, 4,881, 5,096, 13,316, 4,440, 8,212, 13,302, and 5,086 movies were used for data analyses, respectively. After motion correction, CTF estimation, and manually curate exposures, the machine-learning-based particle picking by Topaz software extracted 1,391,844, 2,497,690, 2,157,492, 1,072,343, 2,378,419, 1,026,158, 2,496,421, 3,653,368, and 1,384,33 particles from the datasets, respectively. After particle selections with Heterogeneous Refinement and Ab-initio Reconstruction at the class similarity of 0.6, selected better particles were used in Non-uniform Refinement with the optimizing per-particle defocus option. The datasets having weak density regions were classified with 3DV or 3DC, and the components having stronger density were refined by Local refinement, and/or Non-uniform Refinement. Finally, all datasets provided density maps at 3.07-(**WT_red_**,), 2.65-(***up* state of WT_red_/ -Na^+^**), 3.75-(***middle* state of WT_red_/ -Na^+^**), 3.11-(***down* state of WT_red_/ -Na^+^**), 2.90-(**WT_red_ + KR**), 2.74-(**WT_red_ + AD-42**), 2.61-(***stable* state of NqrB-G141A_red_ + KR**), 2.93-(***shifted* state of NqrB-G141A_red_ + KR**), 3.83-(**NqrB-T236Y_ox_**), 3.31-(**NqrB-T236Y_red_**), 2.66-(**NqrC-T225Y_ox_**), and 2.59-(**NqrC-T225Y_red_**) Å resolution, respectively. These resolutions were estimated based on the gold standard criteria (FSC = 0.143). Furthermore, to capture the overall structure of flexible NqrF subunits, non-uniform refined models (**WT_red_/ -Na^+^**, and **NqrB-G141A_red_ + KR**) were reconstructed with 3DFlex Refinement using a mask covering all subunits. The Flex maps provided by 3DFlex reconstruction are referenced for model building.

### Model building and refinement

The atomic models were built based on our previous cryo-EM structures of Na^+^-NQR (PDB ID: 7XK3 - 7). The initial models were fitted into the density maps with the Chain Refine and Space Refine Zone functions of COOT software^45^. The parts containing weak densities were adjusted with rigid body fitting and morphing. The Flex map output from 3DFlex Reconstruction was used to model the heterogeneous conformations with flexibility. Using the phenix_real_space_refine program^46^, the corrected models were refined with secondary structure and Ramachandran restraints. These results were manually validated with COOT. The iteration of model refinement provided the correct models following the geometry. The model validation was evaluated with Molplobity^47^. To avoid over-fitting, the final models were validated with Fourier shell correlation computed from the final density maps. The model statistics are shown in Supplementary Table 1. Images of structures were generated with UCSF chimeraX software^48^. UCSF ChimeraX was used to compute the electrostatic potential, with default parameters and the Coulombic command with a range of −10 to 10 kcal/(mol·e).

### Molecular dynamics simulation

The all-atom MD simulations of the Na^+^-NQR complex were performed using the cryo-EM structure (PDB ID: 7XK3) and the crystal structure (PDB ID: 8ACW) as the initial structures. For modeling of missing residues at the N-termini (for NqrB, NqrC, and NqrD) and C-termini (for NqrB and NqrC), GalxyFill plug in CHARMM-GUI^49^ was used. The cofactors (FAD, 2Fe–2S^NqrF^, 2Fe–2S^NqrD/E^, FMN^NqrC^, FMN^NqrB^, and RBF) were retained as in the PDB structures. The Nqr structure was embedded in a 150 × 150 Å POPC membrane and solvated with TIP3P water and 150 mM sodium and chloride ions with the Membrane Builder plugin in CHARMM-GUI^50,51^. The CHARMM36 force field^52^ was used except for the cofactors whose parameters were adapted from the works of Kaila group^27,53,54^. The total number of atoms is ∼380,000.

The systems were energy minimized with the restraint of non–hydrogen POPC, protein, and cofactor atoms for 10,000 steps. Then, they were equilibrated in three stages: (1) 100 ps equilibration at *T* = 150 K and no pressure control with the restraint of non–hydrogen POPC, protein, and cofactor atoms, (2) 500 ps equilibration at *T* = 310 K and 1 bar pressure with the restraint of non–hydrogen protein and cofactor atoms, (3) 500 ps equilibration at *T* = 310 K and 1 bar pressure with the restraint of protein–Cα and non–hydrogen cofactor atoms. The restraint force constant was 1.0 kcal mol^-1^ Å^-2^ for each atom. The production runs were performed at *T* = 310 K and 1 bar pressure without restraints. Long-range electrostatic interactions were calculated by the particle mesh Ewald method^55^ with a direct space cut off of 12 Å. Langevin dynamics with 1 ps^-1^ damping coefficient was used for temperature control at *T* = 310 K, and the Nosé–Hoover Langevin piston was used for pressure control at 1 bar^56―58^. The integration timestep was set to 2 fs by applying the SHAKE method^59^ to bonds involving hydrogen. Targeted MD (TMD) simulations were also performed using an outward-open conformation of NqrD/E (shifted state of NqrB-G141A_red_ + KR) as the reference. The force constant for TMD force was set to 500 kcal mol^-1^ Å^-2^ and applied to the Cα atoms of NqrD and NqrE. All MD simulations were performed using NAMD 2.12 /2.14/ 3.0^60^.

## Abbreviations

CBB: Coomassie Brilliant Blue
Cryo-EM: cryogenic electron microscope
DDM: *n*-dodecyl *β*-maltoside
EPR: electron paramagnetic resonance
FAD: flavin adenine dinucleotide
FMN: flavin mononucleotide
IC_50_: the molar concentration needed to reduce the control enzymatic activity by 50%
MD: molecular dynamics
NADH: nicotinamide adenine dinucleotide reduced form
NAD^+^: nicotinamide adenine dinucleotide oxidized form
Na^+^-NQR: Na^+^-pumping NADH-ubiquinone oxidoreductase
NTA: nitrilotriacetic acid
PAGE: polyacrylamide gel electrophoresis
RBF: riboflavin
RNF: Rhodobacter nitrogen fixing” protein
RT: room temperature
TMH: transmembrane helix
SDS: sodium dodecyl sulfate
UQ: ubiquinone
UQH_2_: ubiquinol (reduced form of ubiquinone)
WT: wild-type

## Data availability

The cryo-EM maps have been deposited in the EMDB under accession codes, 64518 (**WT_red_**), 63872 (**NqrC-T225Y_ox_**), 64059 (**NqrC-T225Y_red_**), 64060 (**NqrB-T236Y_ox_**), 64061 (**NqrB-T236Y_red_**), 64063 (**WT_red_/ -Na^+^**, “*up*” state), 64064 (**WT_red_/ -Na^+^**, “*middle*” state), 64065 (**WT_red_/ -Na^+^**, “*down*” state), 64062 (**WT_red_ + KR**), 64069 (**WT_red_ + AD-42**), 64066 (**NqrB-G141A_red_ + KR**, “*stable*” state), and 64068 (**NqrB-G141A_red_ + KR**, “*shifted*” state). The atomic models have been deposited in the Protein Data Bank under accession codes, 9UUU (**WT_red_**), 9U5G (**NqrC-T225Y_ox_**), 9UD2 (**NqrC-T225Y_red_**), 9UD3 (**NqrB-T236Y_ox_**), 9UD4 (**NqrB-T236Y_red_**), 9UD6 (**WT_red_/ -Na^+^**, “*up*” state), 9UD8 (**WT_red_/ -Na^+^**, “*middle*” state), 9UD9 (**WT_red_/ -Na^+^**, “*down*” state), 9UD5 (**WT_red_ + KR**), 9UDG (**WT_red_ + AD-42**), 9UDA (**NqrB-G141A_red_ + KR**, “*stable*” state), and 9UDF (**NqrB-G141A_red_ + KR**, “*shifted*” state). The initial model of Na^+^-NQR with and without bound inhibitors for model building is accessible in PDB under accession number 7XK3 (oxidized form of Na^+^-NQR: **WT** ^7^), 7XK6 (oxidized form of Na^+^-NQR with bound AD-42: **WT_ox_ + AD-42**), and 7XK7 (oxidized form of Na^+^-NQR with bound KR: **WT_ox_ + KR**) ^9^. The data that support the findings of this study are available from the corresponding author upon reasonable request.

## Acknowledgements

We thank Dr. Joel E. Morgan (Rensselaer Polytechnique Institute) for the valuable discussion and comments during this work. We acknowledge the Research Support Project for Life Science and Drug Discovery (Basis for Supporting Innovative Drug Discovery and Life Science Research (BINDS)) from AMED under Grant Number JP23ama 121001. The computation was performed at the Research Center for Computational Science, Okazaki, Japan (Project: 24-IMS-C198, 25-IMS-C227). This work was financially supported by JSPS KAKENHI (22KJ1795 to M.I-F, 24K08729 to T.M., 23K23858 and 25H02299 to K.O., 21H02130 to H.M., and 25K01958 to M.M.), the Uehara Memorial Foundation (to J.K.), Takeda Science Foundation (to J.K.), and the National Institutes of Health (R56-AI132580 and R01AI181279 to B.B.).

## Author contributions

M.I-F., J.K., T.M., B.B. H.M., and M.M. designed the research; M.I-F., T.M., and B.B expressed and purified Na^+^-NQR; M.I-F., J.K., and T.M. performed structural determination and built the atomic models; T.S. and O.K. performed MD simulation; M.I-F., T.S., J.K, T.M., K.O, T. K., B.B., H.M. and M.M analyzed data; K.J. and M.M. directed the project and wrote the paper with M I-F., T.M., K.O., B.B. and H.M.

## EXTENDED DATA FIGURES

**Extended Data Fig. 1:**
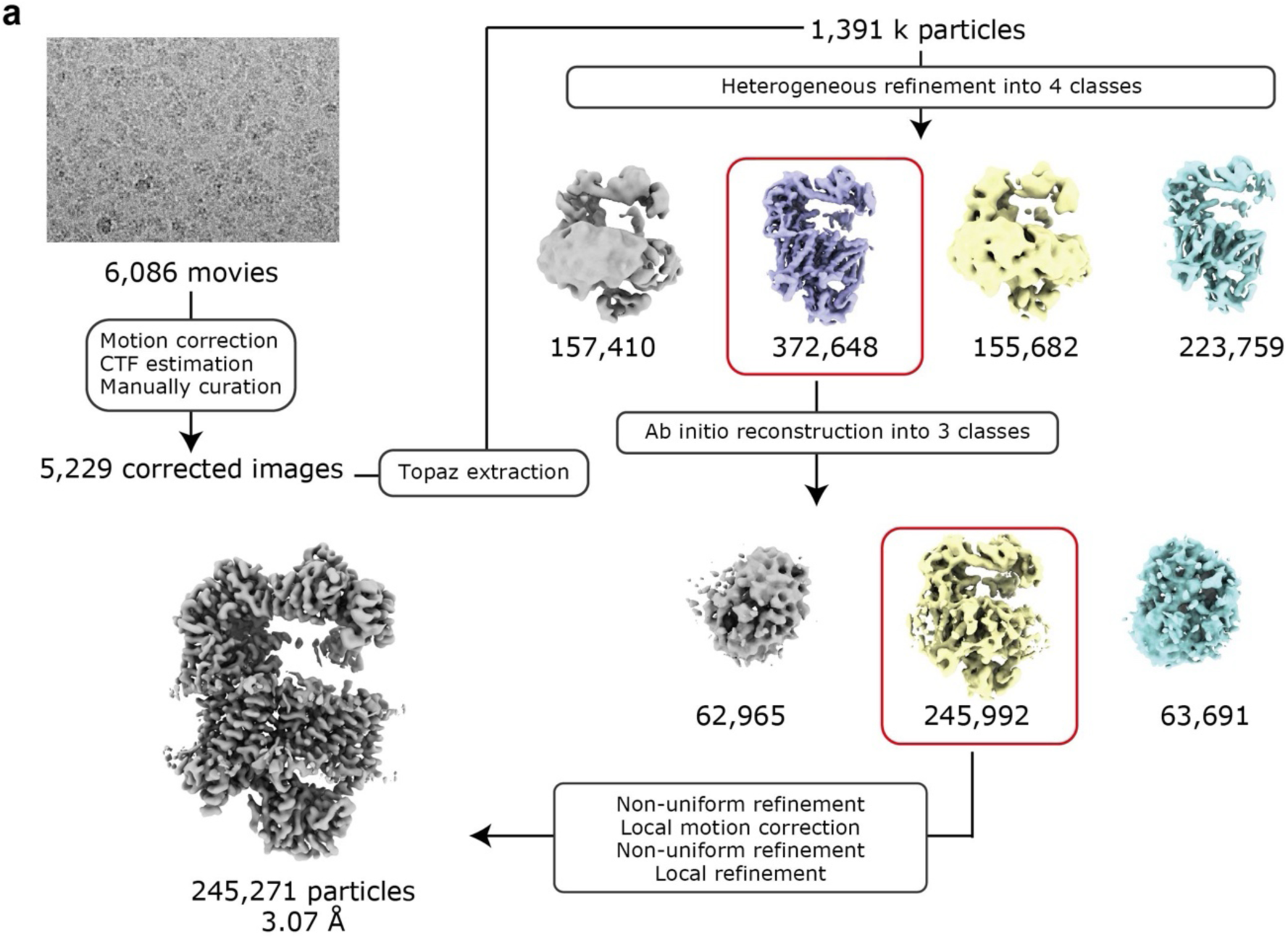
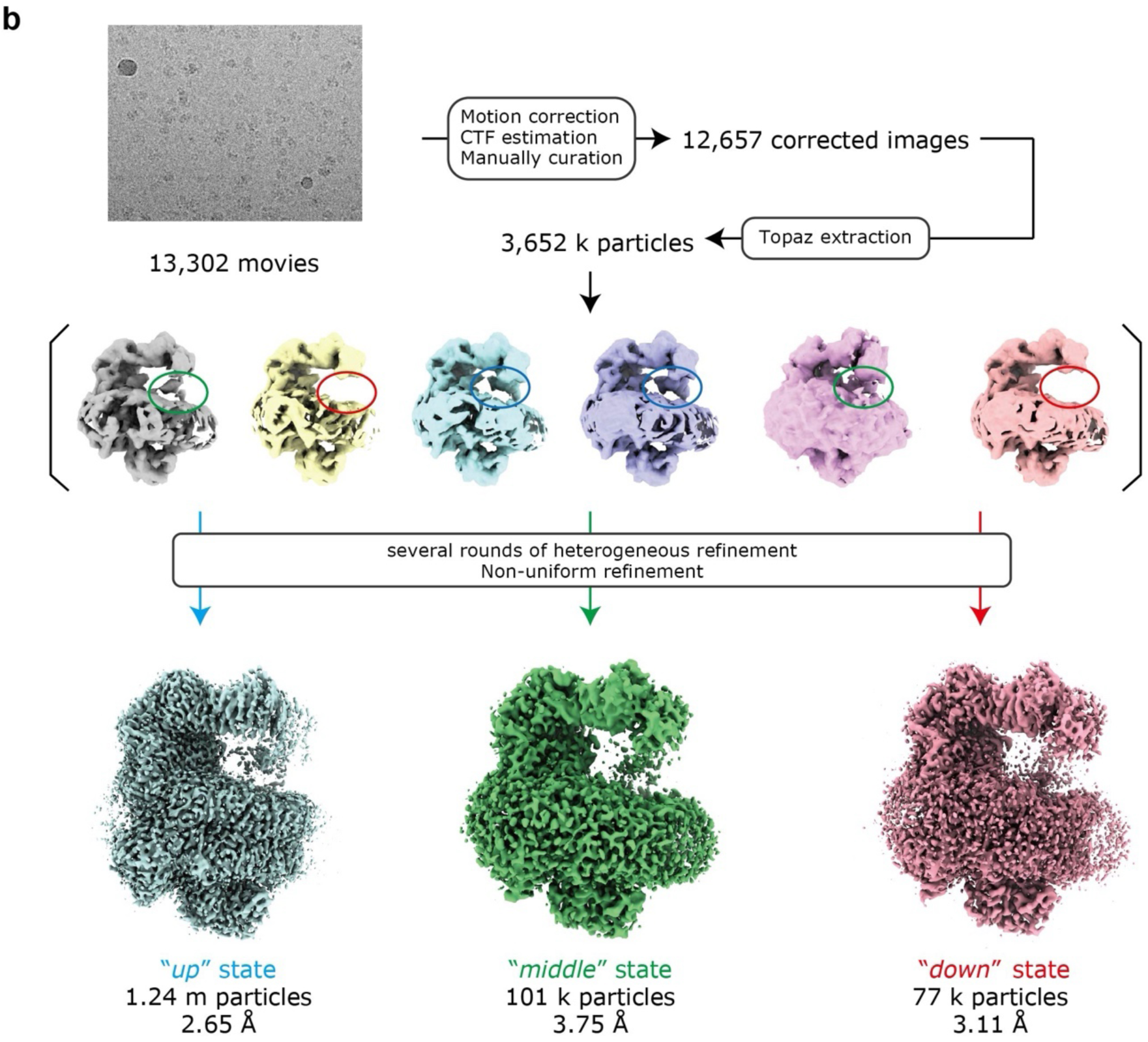
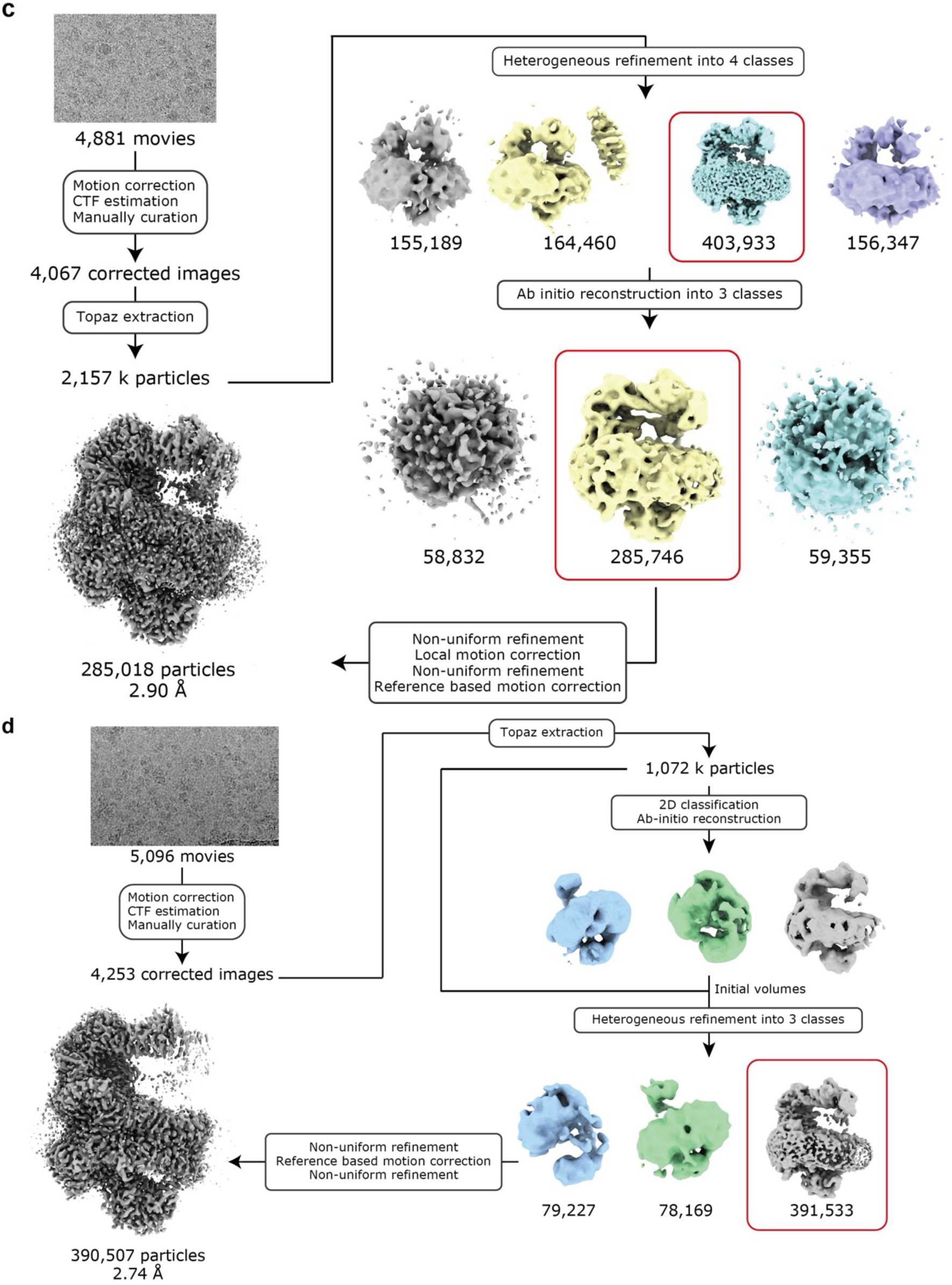
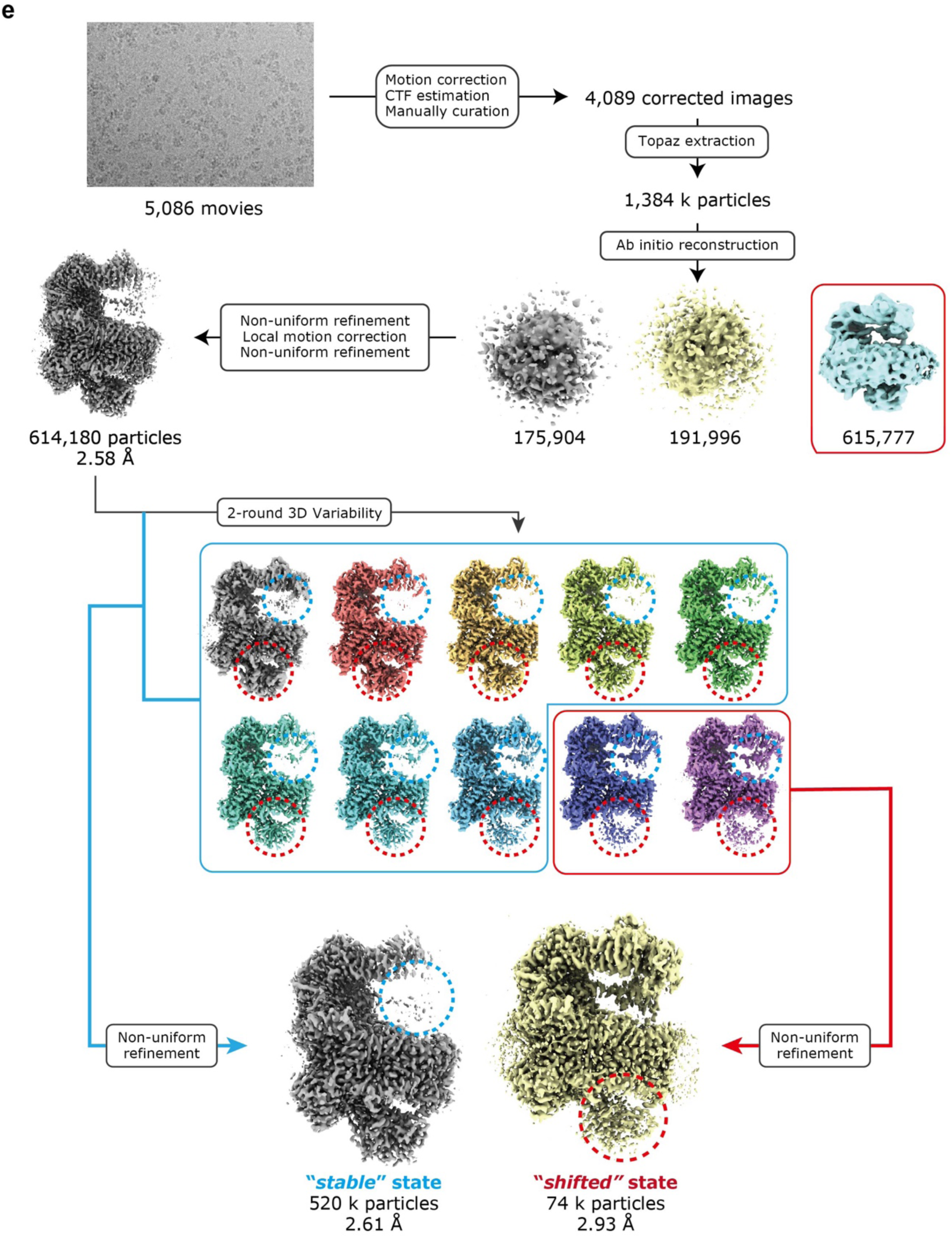
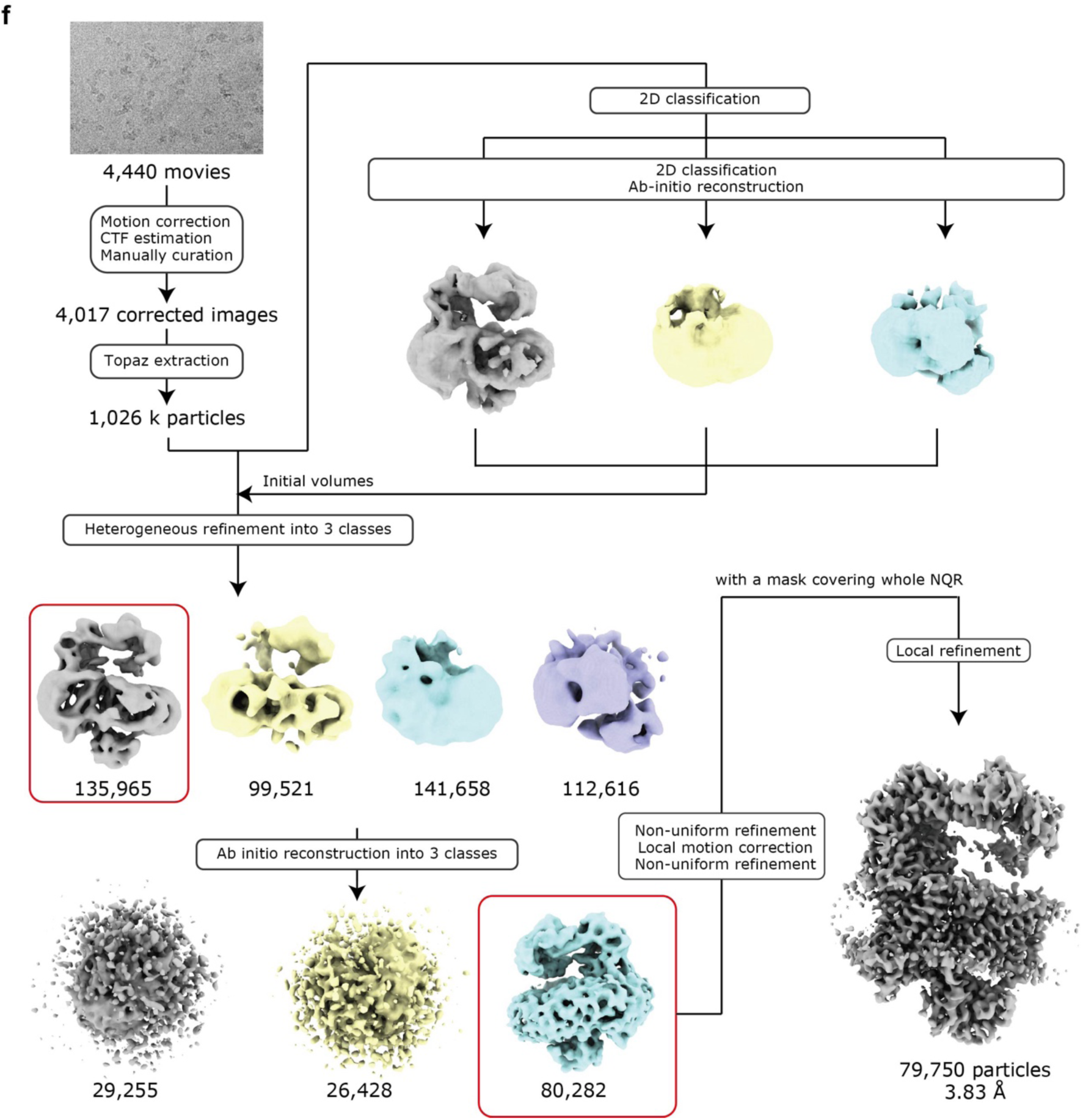
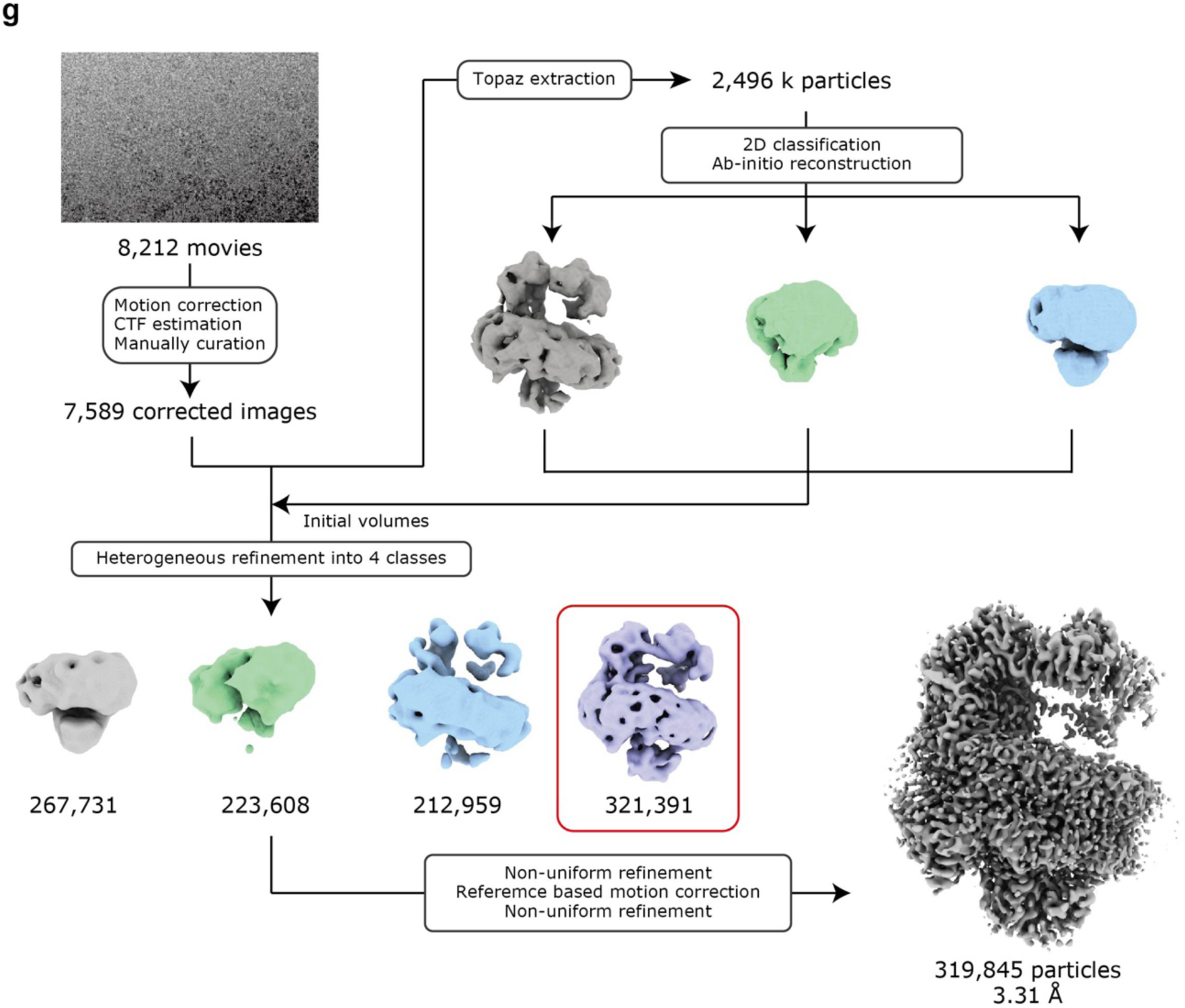
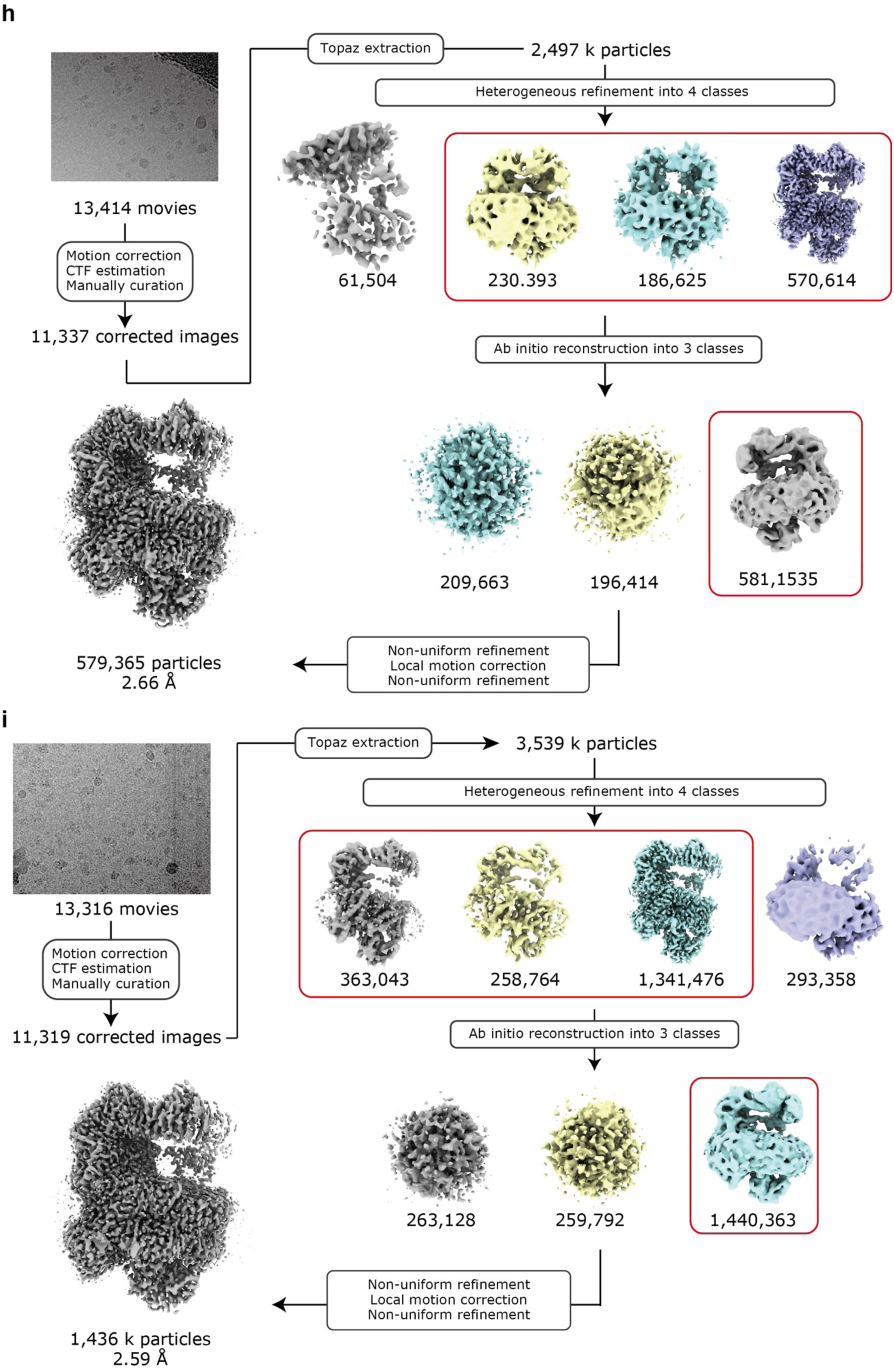
The scheme of single particle analysis with cryoSPARC. ***a:* WT_red_**, ***b***: **WT_red_/ - Na^+^**, ***c***: **WT_red_ + KR**, ***d***: **WT_red_ + AD-42**, ***e***: **NqrB-G141A_red_ + KR**, ***f***: **NqrB-T236Y_ox_**, ***g***: **NqrB-T236Y_red_**, ***h***: **NqrC-T225Y_ox_**, ***i***: **NqrC-T225Y_red_**. The classes surrounded by *red squares* were used in the next step. In ***e***, *red* and *blue circles* indicate the hydrophilic domain of NqrC and the ferredoxin-like domain of NqrF, respectively. The image processing for each dataset was implemented following each flow chart.

**Extended Data Fig 2:**
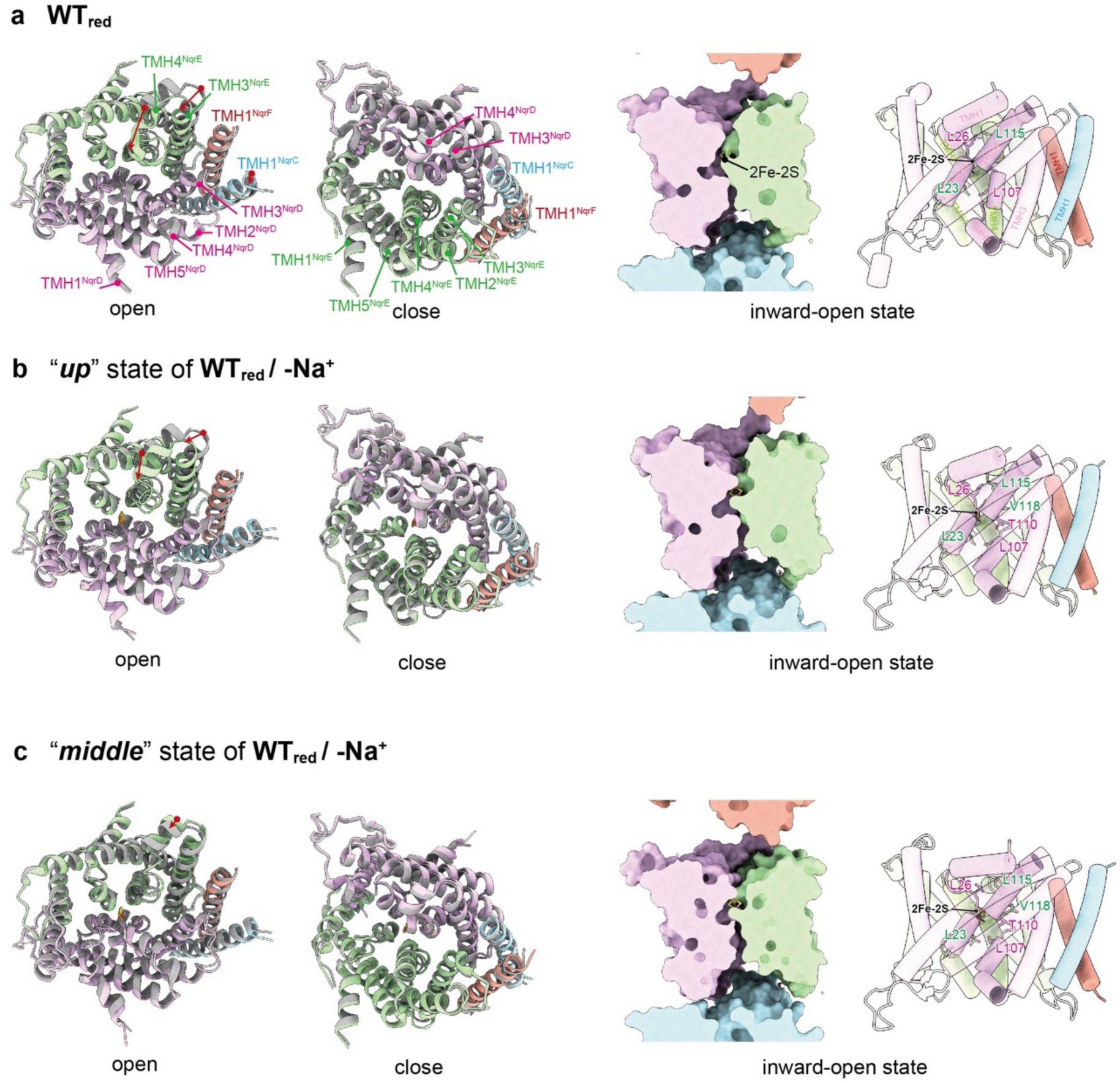
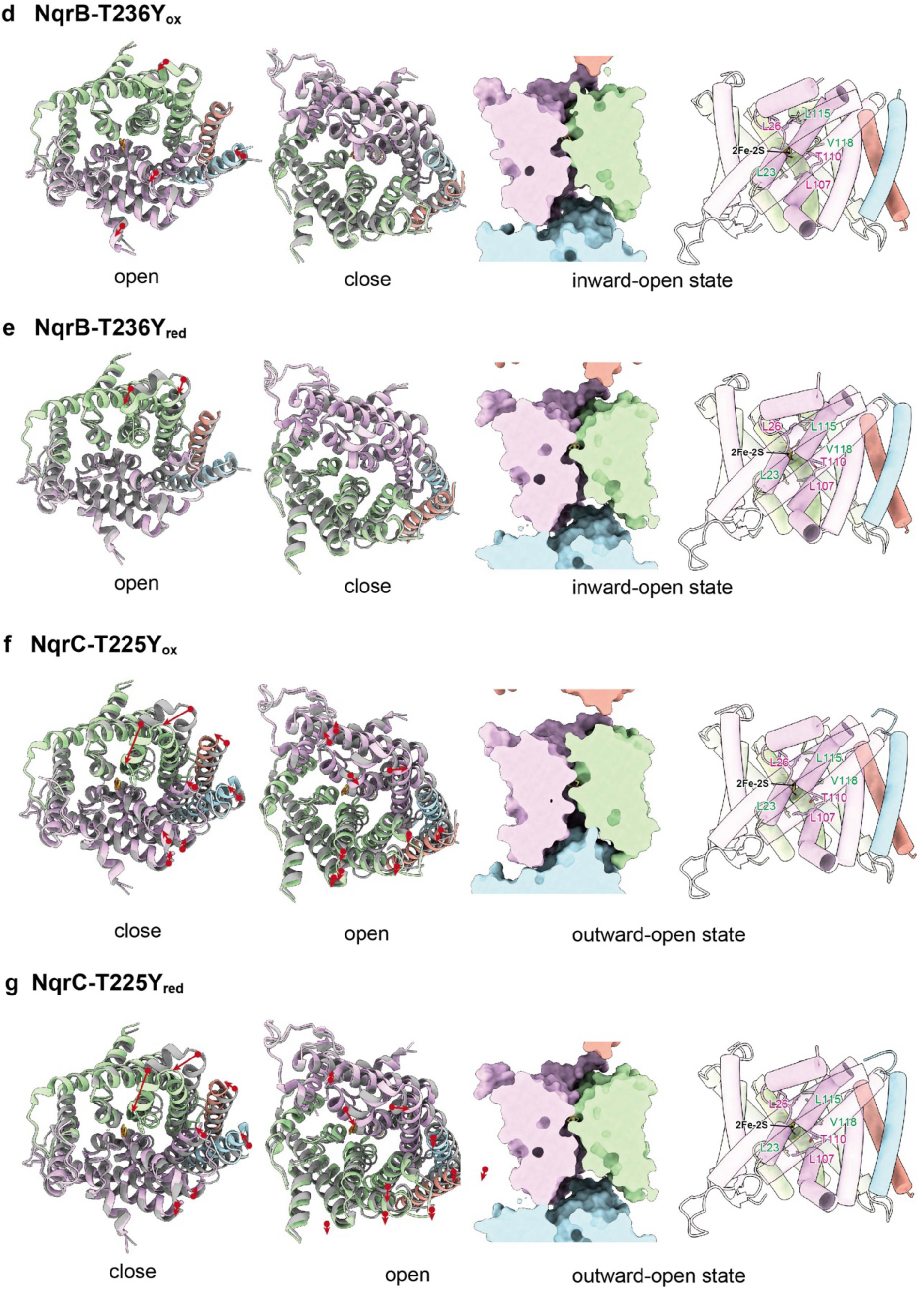
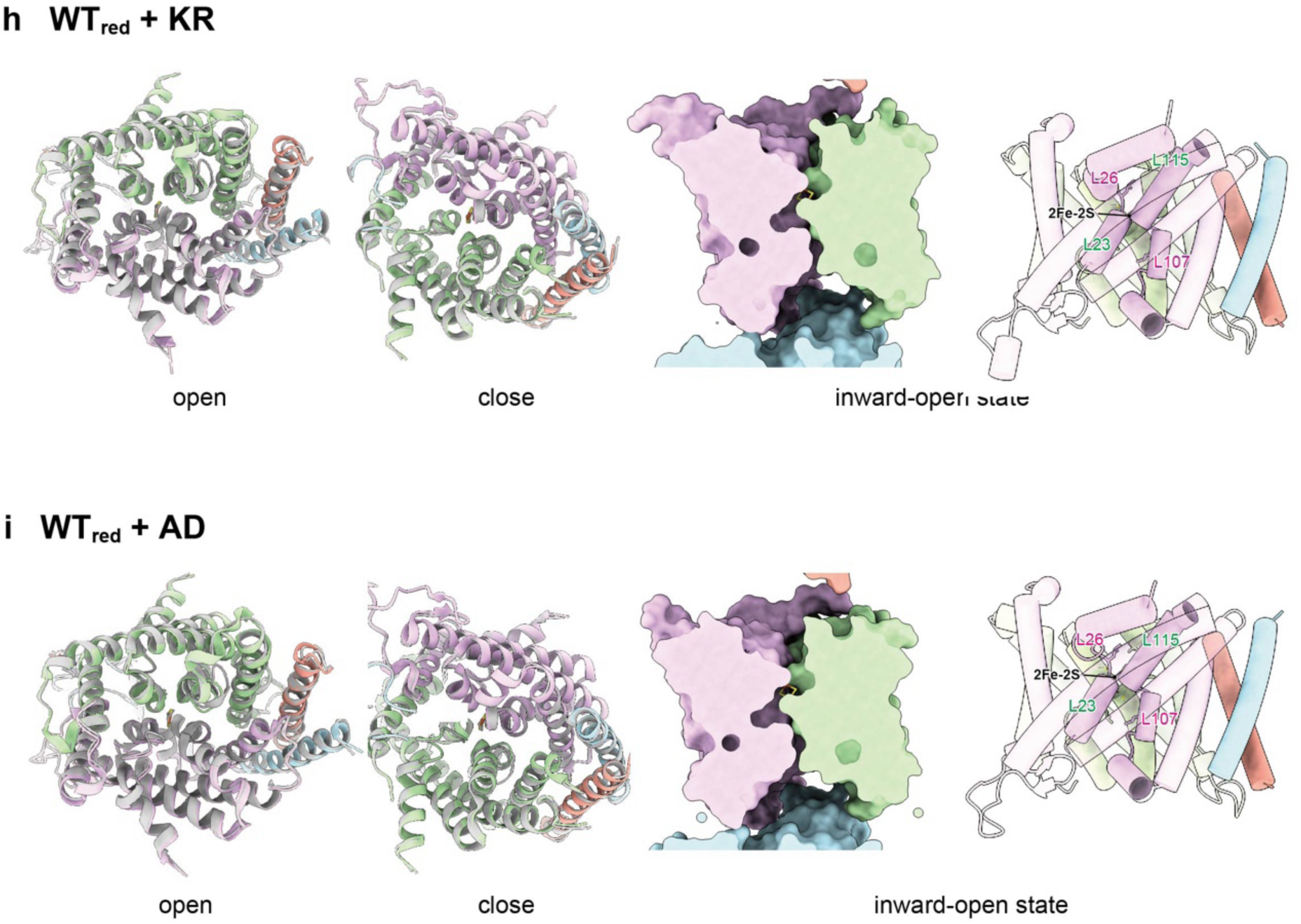
TMHs shifts. The TMHs architecture of NqrCDEF. The *left* and *middle* panels show cytoplasmic views and periplasmic views, respectively. To help visualize the transition, the **WT_ox_** structure is superimposed in *gray*. The *right* panel shows the sliced view. ***a***: **WT_red_**, ***b***: “***up***” state from **WT_red_/-Na^+^**, ***c***: “***middle***” state from **WT_red_/-Na^+^**, ***d***: **NqrB-T236Y_ox_**, ***e***: **NqrB-T236Y_red_**, ***f***: **NqrC-T225Y_ox_**, ***g***: **NqrC-T225Y_red_, *h***: **WT_red_ + KR,** and ***i***: **WT_red_ + AD** are compared with **WT_red_**, respectively.

**Extended Data Fig. 3:**
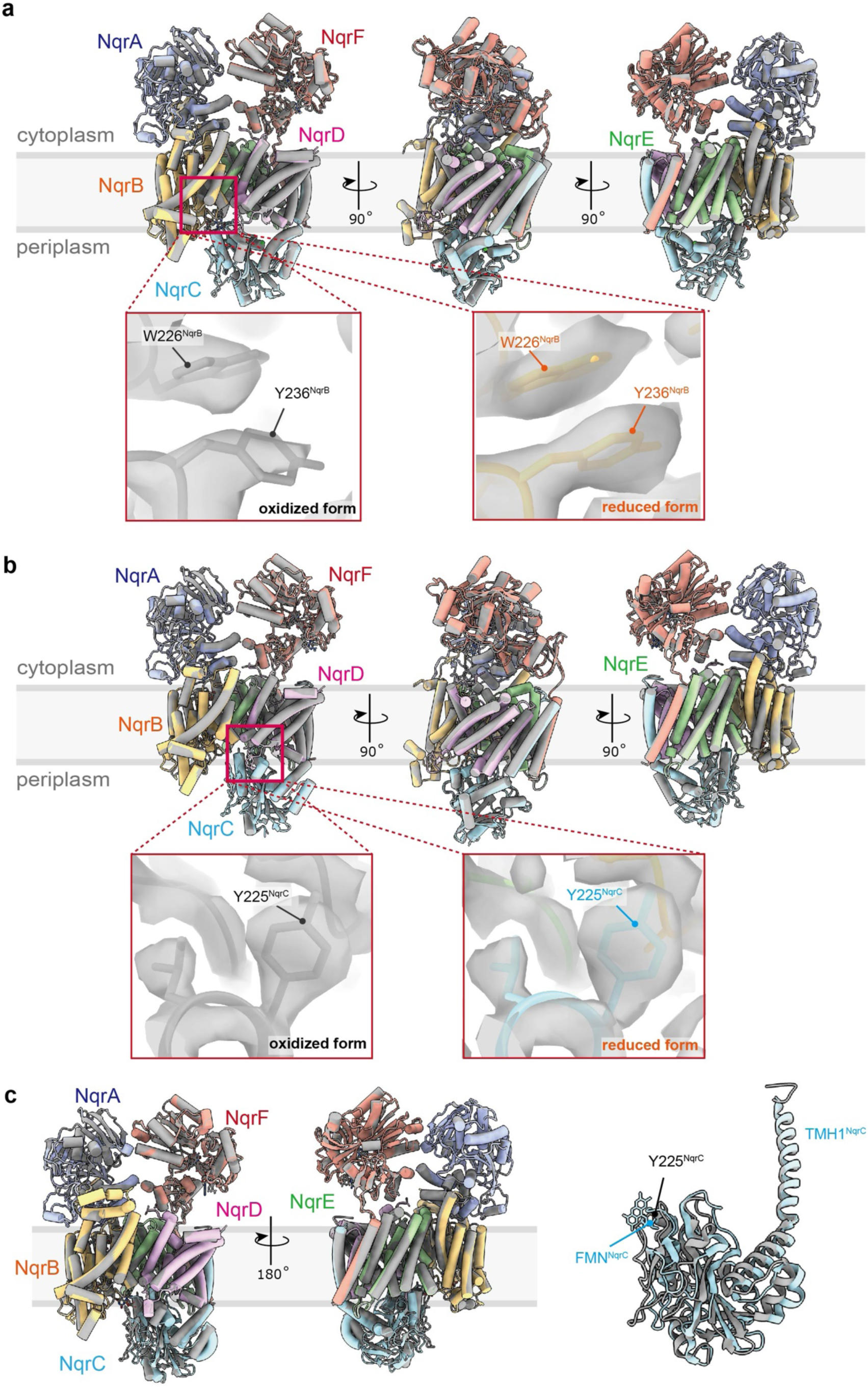
The architecture of FMN deletions. ***a,*** Comparison of **NqrB-T236Y_ox_** (*gray*) and **NqrB-T236Y_red_** (*color*). *Red square* shows the position of the point mutation. NqrB-Y236 in each form is zoomed up with a density map in the bottom pictures. ***b,*** Comparison of **NqrC-T225Y_ox_** (*gray*) and **NqrC-T225Y_red_** (*color*). *Red square* shows the position of the point mutation. NqrC-Y225 in each form is zoomed up with a density map in the bottom pictures. ***c,*** Comparison of **NqrC-T225Y_red_** (*gray*) and the “*shifted*” state of **NqrB-G141A_red_ + KR** (*color*). NqrC is picked up in *right* panel.

**Extended Data Fig. 4:**
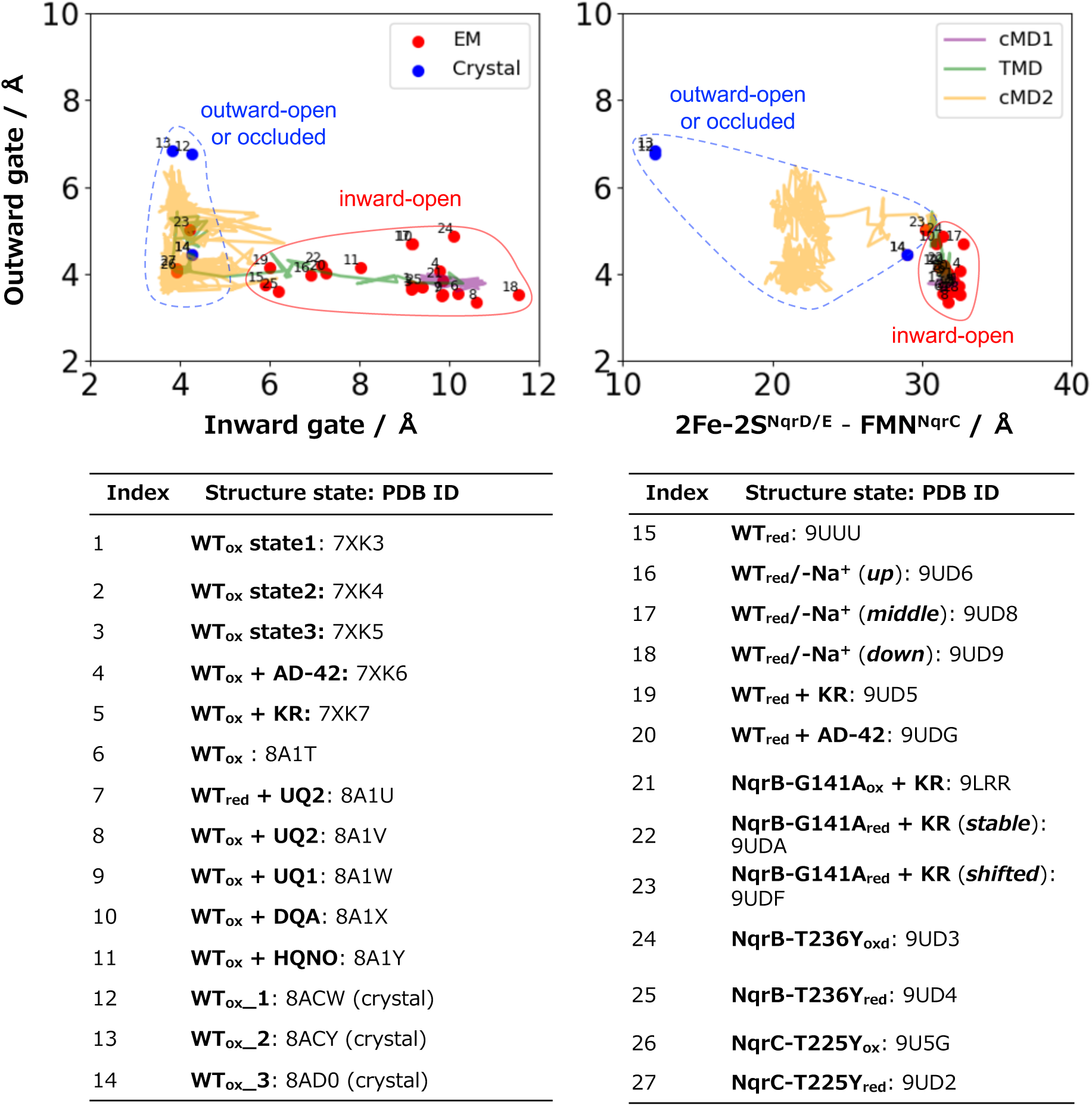
MD simulation trajectory compared to experimental structures. The inward gate and outward gate size are defined by the distances between L26^NqrD^ and L115^NqrE^, and between L107^NqrD^ and L23^NqrE^, respectively (Fig. 4a). The *left* panel shows a 2D plot of the inward gate and the outward gate distances. The *right* panel shows a 2D plot of the cofactors 2Fe-2S^NqrD/E^−FMN^NqrC^ distance and the outward gate distance. The numbers are from the bottom table.

**Extended Data Fig. 5:**
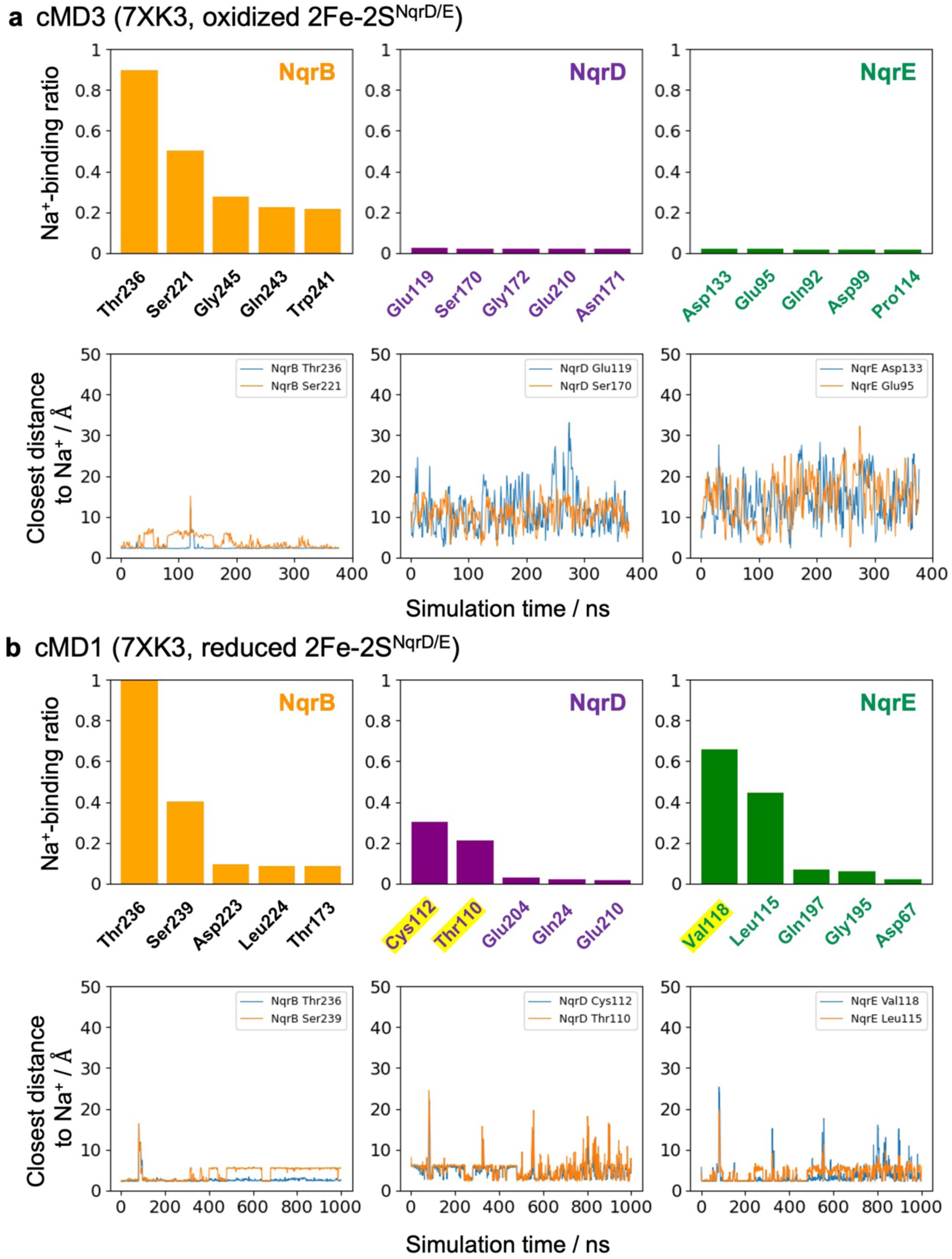
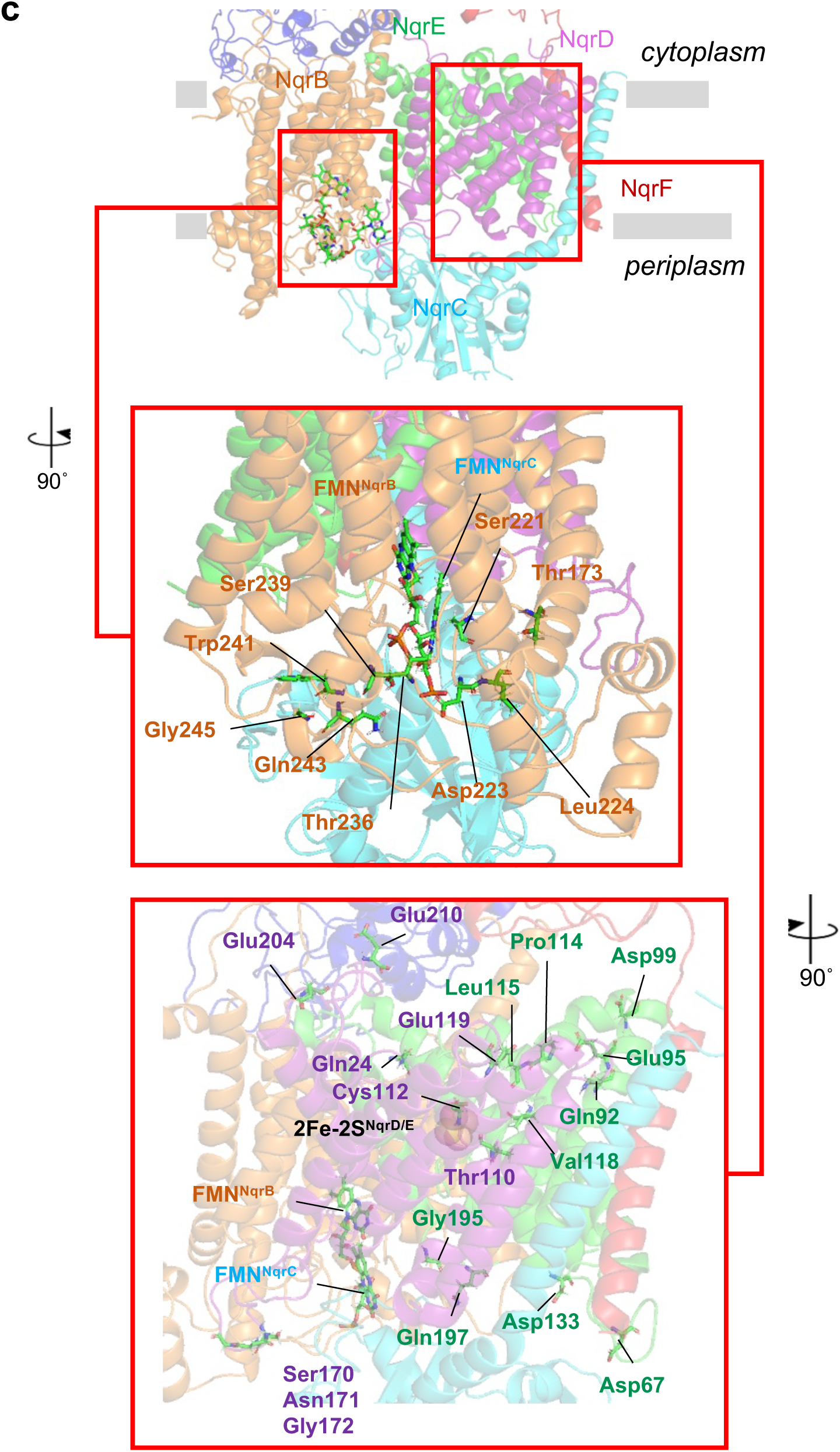
Analysis of amino acid residues that are involved in Na^+^ binding. **a-b,** Simulation is calculated at the scale of 400 ns. Contact ratio corresponds to the ratio of Na^+^-binding frame to total frame. Residues showing > 0.01 of contact ratio and the > 3 Å of distance to were listed up. The *top* panels show the top five residues that have high Na^+^ binding ratios for NqrB, NqrD, and NqrE, respectively. The Na^+^ binding was defined through the distance being less than 3Å. The *bottom* panels show the time course of the closest distances between the residues and Na^+^. ***a***, Analysis of MD simulation with oxidized cofactors (cMD3). ***b***, Analysis of MD simulation with reduced 2Fe-2S^NqrD/E^ cofactor (cMD1). ***c***, The locations of amino acid residues in NqrB and NqrD/E that show a high Na^+^ binding ratio in MD simulations. The *top* panel indicates the relative positions of the two regions containing amino acid residues with a high Na^+^ binding ratio in TMHs. The *middle* panel shows the region around the FMN binding site in NqrB. The *bottom* panel shows the region of TMHs^NqrCDEF^ bundle.

## Notes

### Competing Interest Statement

The authors have declared no competing interest.

### Summary of Updates

I noticed that the ORCID of one of my co-authors (Takehito Seki) is incorrect. I updated his ORCID as follows: Takehito Seki: 0009-0006-1830-8566 (https://orcid.org/0009-0006-1830-8566). There are no changes in the manuscript file or supplementary information.

