## Supplementary Information (Supplementary Figs. 1-11, Supplementary Tables 1-3) for "The Na^+^-pumping mechanism driven by redox reactions in the NADH-quinone oxidoreductase from *Vibrio cholerae* relies on dynamic conformational changes"

[1]*Division of Applied Life Sciences, Graduate School of Agriculture, Kyoto University, Kyoto, Kyoto 606-8502, Japan,* [2] *Center for Biotechnology and Interdisciplinary Studies, Rensselaer Polytechnic Institute, Troy, NY 12180, United States,* [3]*Research Center for Computational Science, Institute for Molecular Science, National Institutes of Natural Sciences, Okazaki, Aichi 444-8585, Japan,* [4]*Graduate Institute for Advanced Studies, SOKENDAI, Okazaki, Aichi 444-8585, Japan,* [5]*Faculty of Applied Biology, Kyoto Institute of Technology, Kyoto, Kyoto 606-8585, Japan,* [6] *Institute for Protein Research, The University of Osaka, Suita, Osaka565-0871, Japan.*

†M.I-F. and T.S. contributed equally to this work.

*Correspondence should be addressed:
Masatoshi Murai and Jun-ichi Kishikawa.

**Supplementary Fig. 1:** Data statistics and resolutions of datasets.
**Supplementary Fig. 2:** 3D Flex analysis of the complete $Na^+$-NQR
**Supplementary Fig. 3:** The binding site and density map of NADH
**Supplementary Fig. 4:** The structures of reduced $Na^+$-NQR wild-type with bound inhibitors
**Supplementary Fig. 5:** Structure and binding form of $Na^+$-NQR inhibitor
**Supplementary Fig. 6:** The structural similarities of each subunit
**Supplementary Fig. 7:** $Na^+$ and water spatial densities from conventional MD simulations
**Supplementary Fig. 8:** Conserved amino acid residues in NqrD and NqrE
**Supplementary Fig 9:** Additional five independent trajectories of cMD2 following TMD
**Supplementary Fig. 10:** The proposed catalytic mechanism of $Na^+$-NQR
**Supplementary Fig. 11:** Water spatial density from conventional MD (cMD1).
**Supplementary Table 1:** Statics of cryo-EM data, refinement, and validation of $Na^+$-NQR.
**Supplementary Table 2:** List of atomistic MD simulations.
**Supplementary Table 3:** $Na^+$ binding site by MD simulation.

a

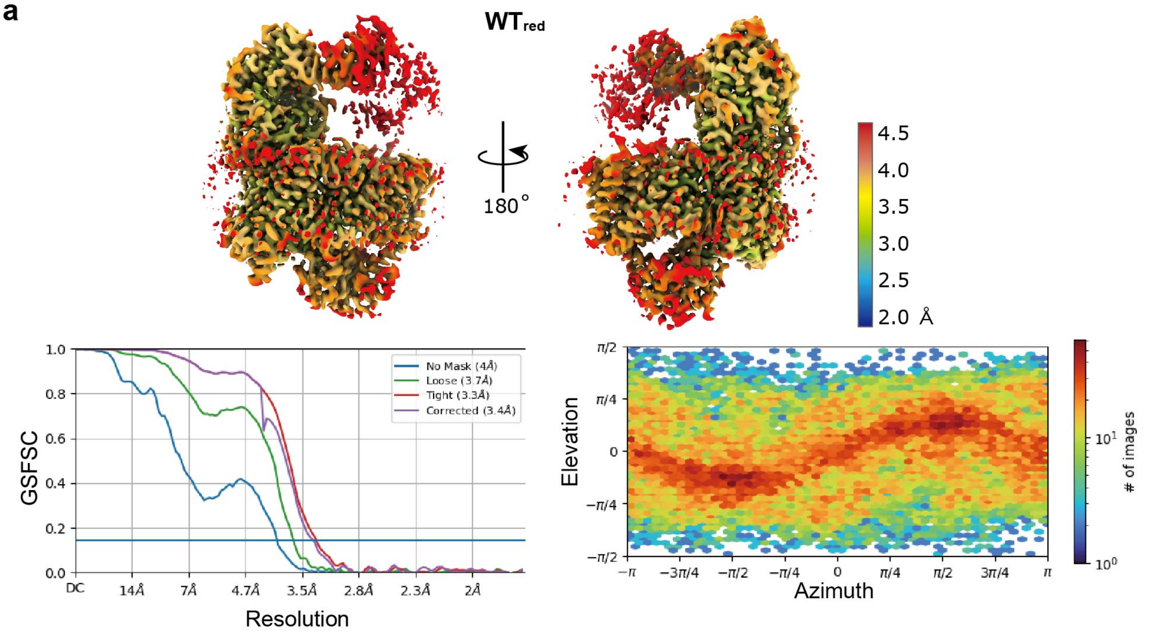


**Supplementary Fig. 1 (continued)**

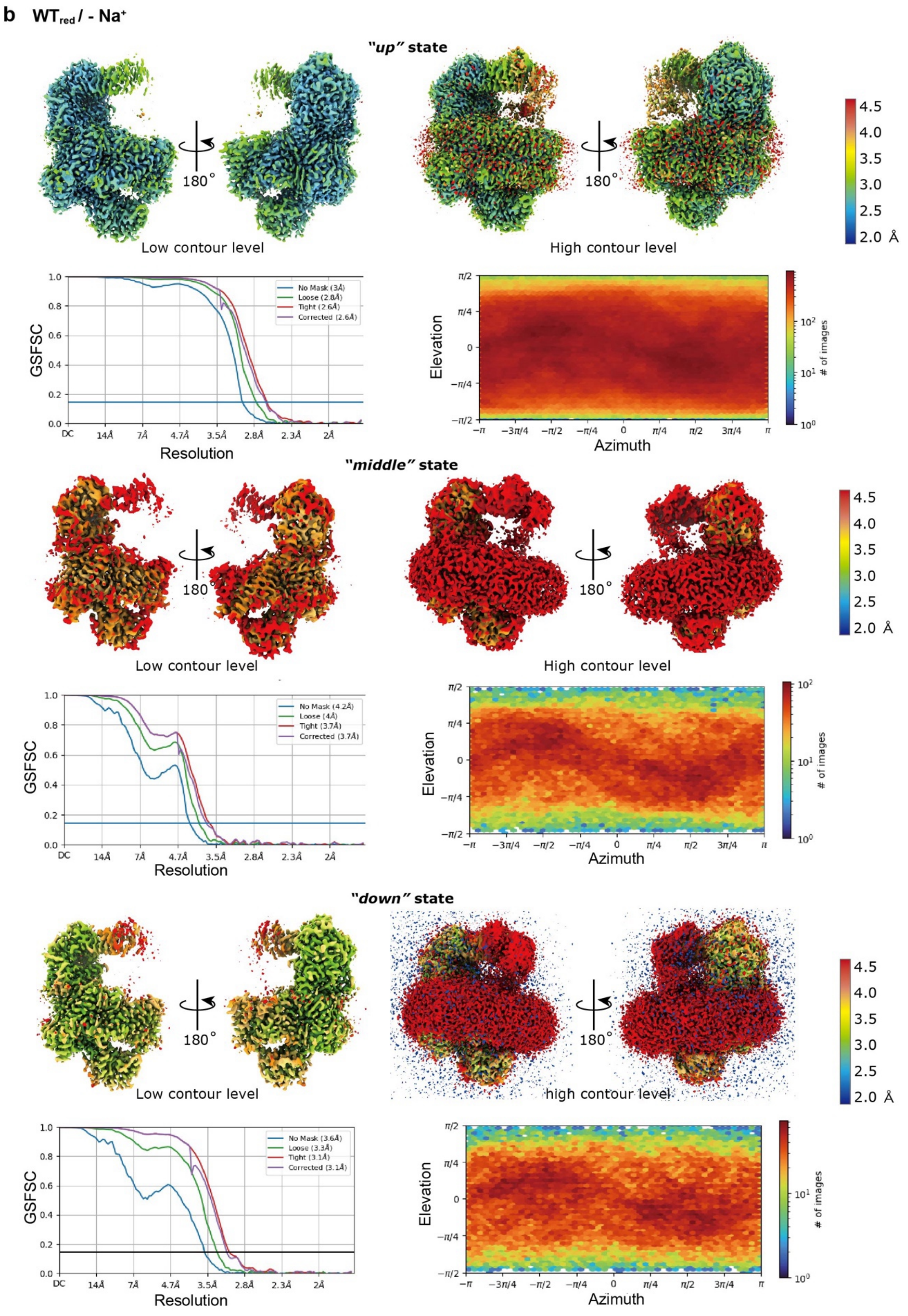


**Supplementary Fig. 1 (continued)**

**c** $WT_{red}$ + KR

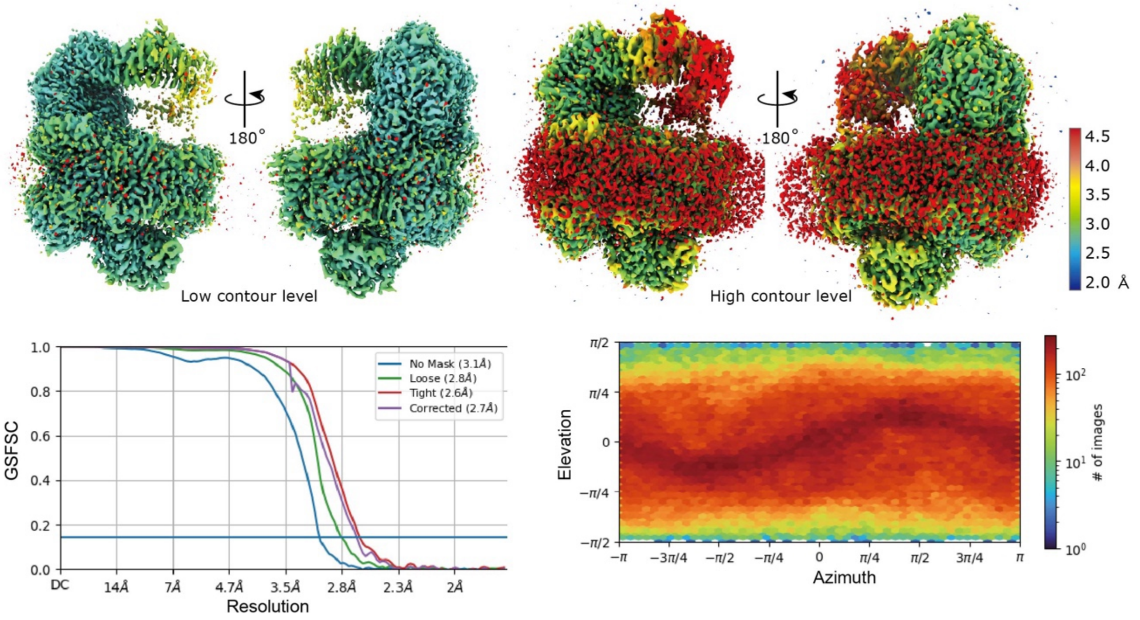


**d** $WT_{red}$ + AD-42

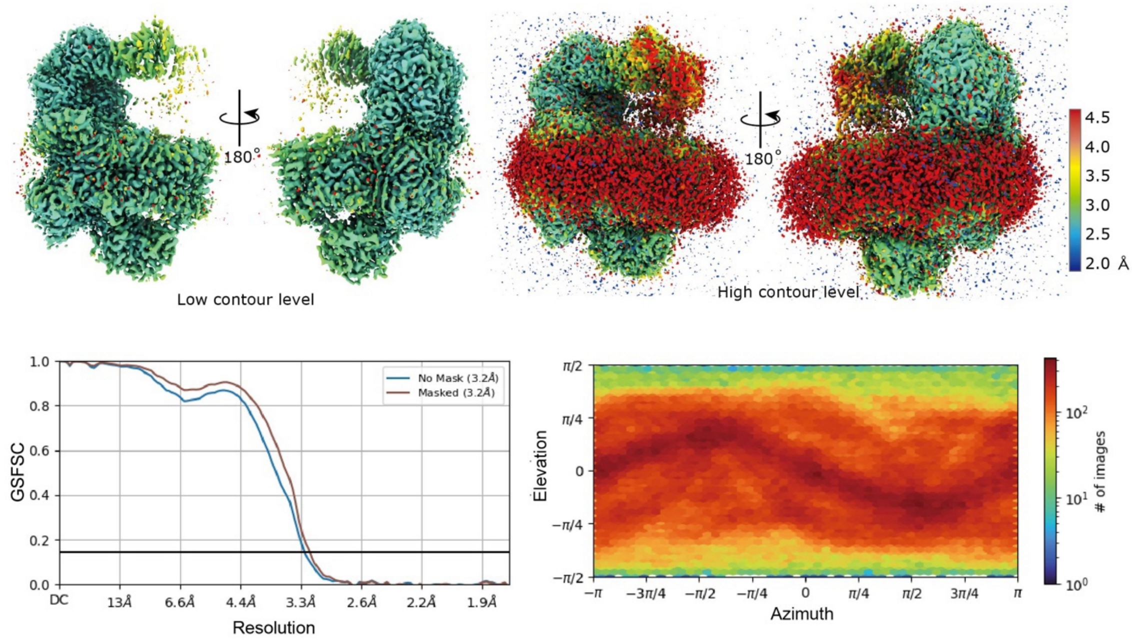


**Supplementary Fig. 1 (continued)**

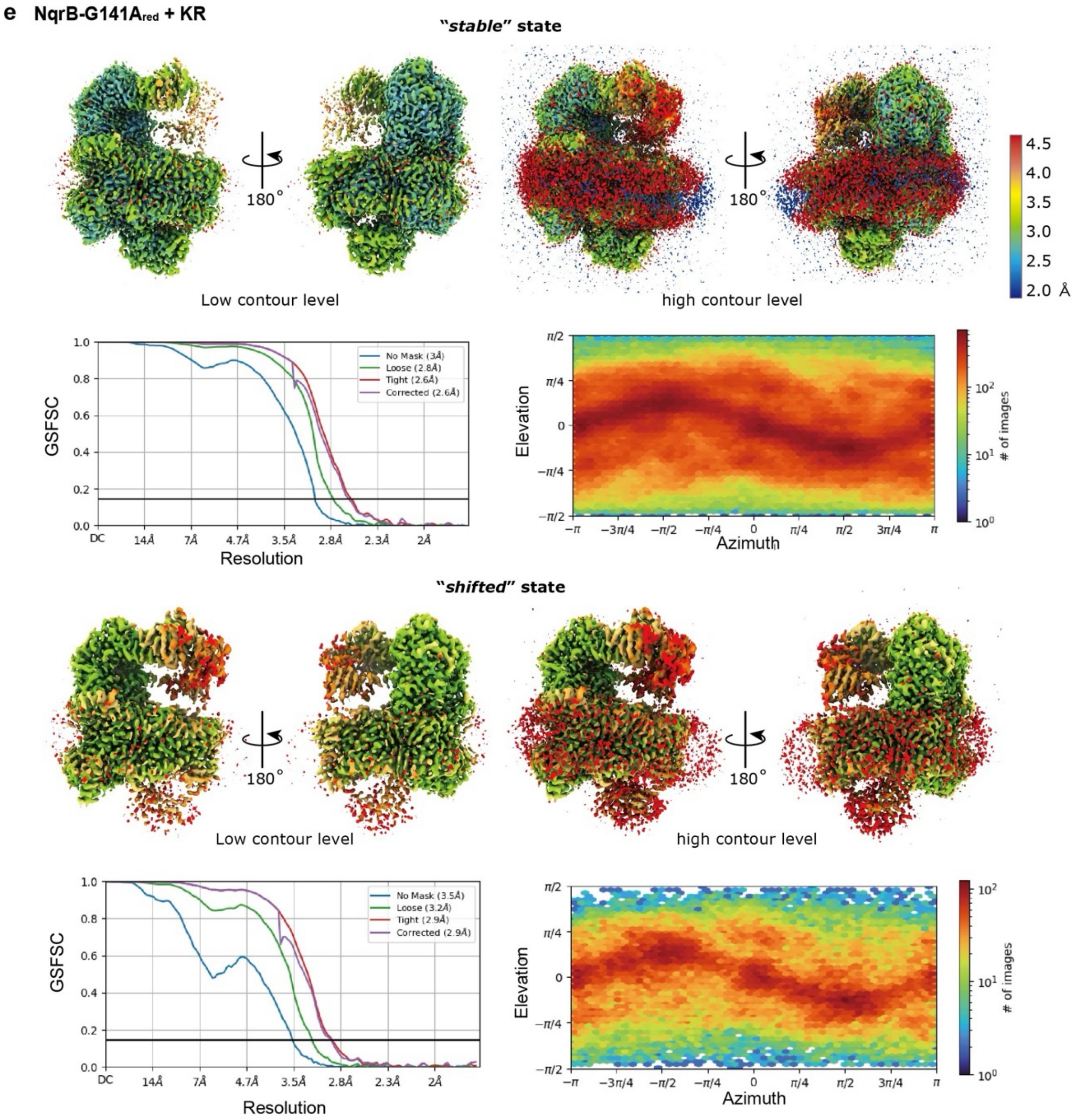


**Supplementary Fig. 1 (continued)**

f

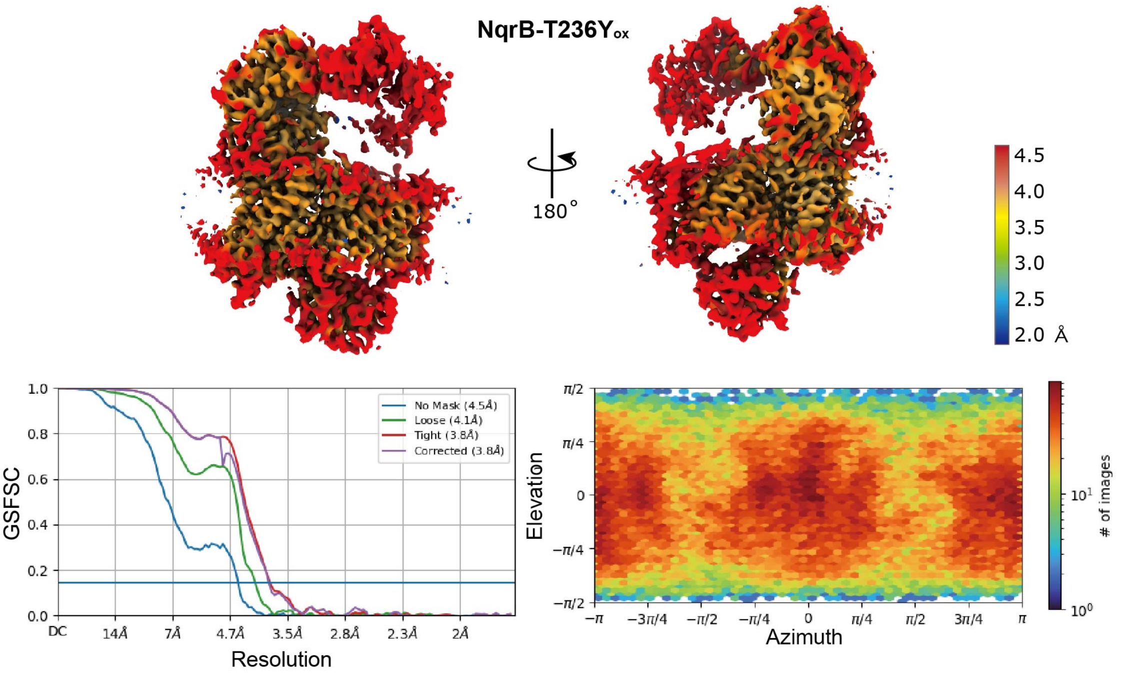


g

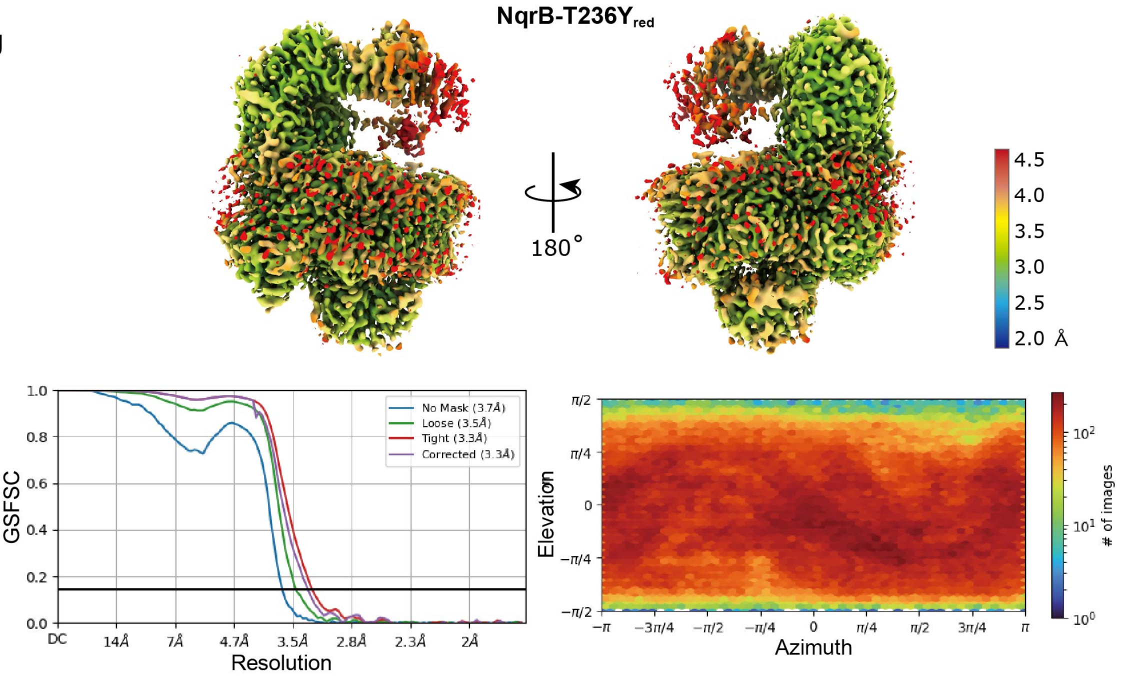


**Supplementary Fig. 1 (continued)**

**h**

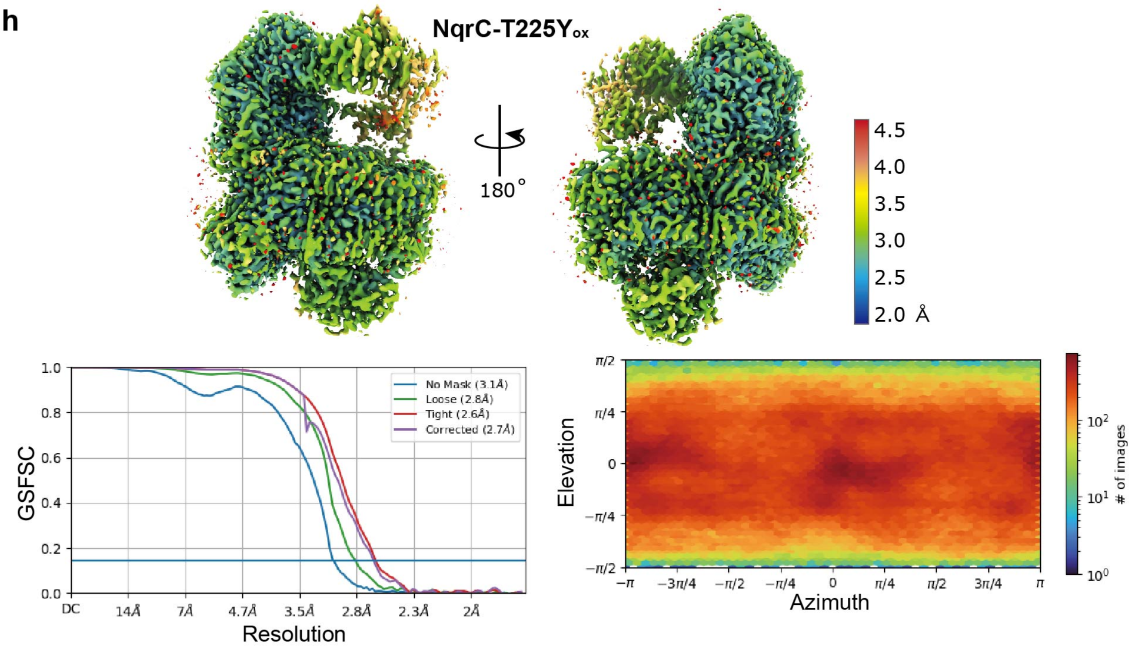


**i**

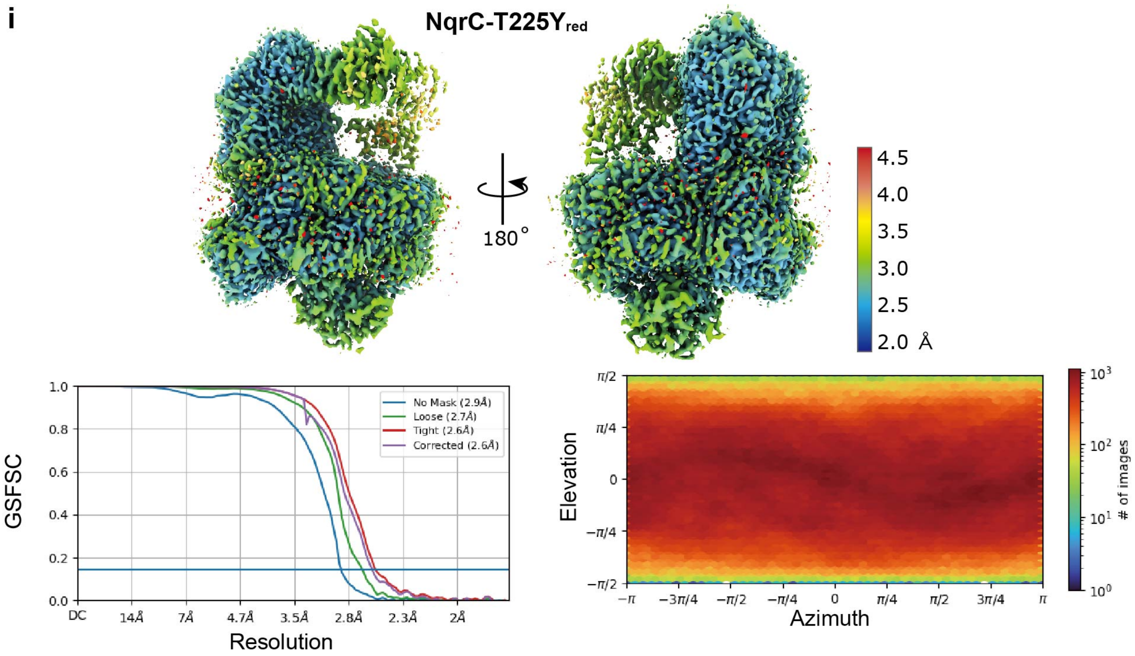


**Supplementary Fig. 1: Data statistics and resolutions of datasets.** The upper panels show the local resolution maps; the lower left panels show the golden standard Fourier shell correlation (GSFSC) curves, and the bottom right panels show 2D distribution histograms for ***a:*** **$WT_{red}$**, ***b:*** **$WT_{red}$/- $Na^+$**, ***c:*** **$WT_{red}$ + KR**, ***d:*** **$WT_{red}$ + AD-42**, ***e:*** **NqrB-G141$A_{red}$ + KR**, ***f:*** **NqrB-T236$Y_{ox}$**, ***g:*** **NqrB-T236$Y_{red}$**, ***h:*** **NqrC-T225$Y_{ox}$**, ***i:*** **NqrC-T225$Y_{red}$**. The image processing for each dataset was implemented following each flow chart. A higher resolution is *blue*, and a lower resolution is *red*.

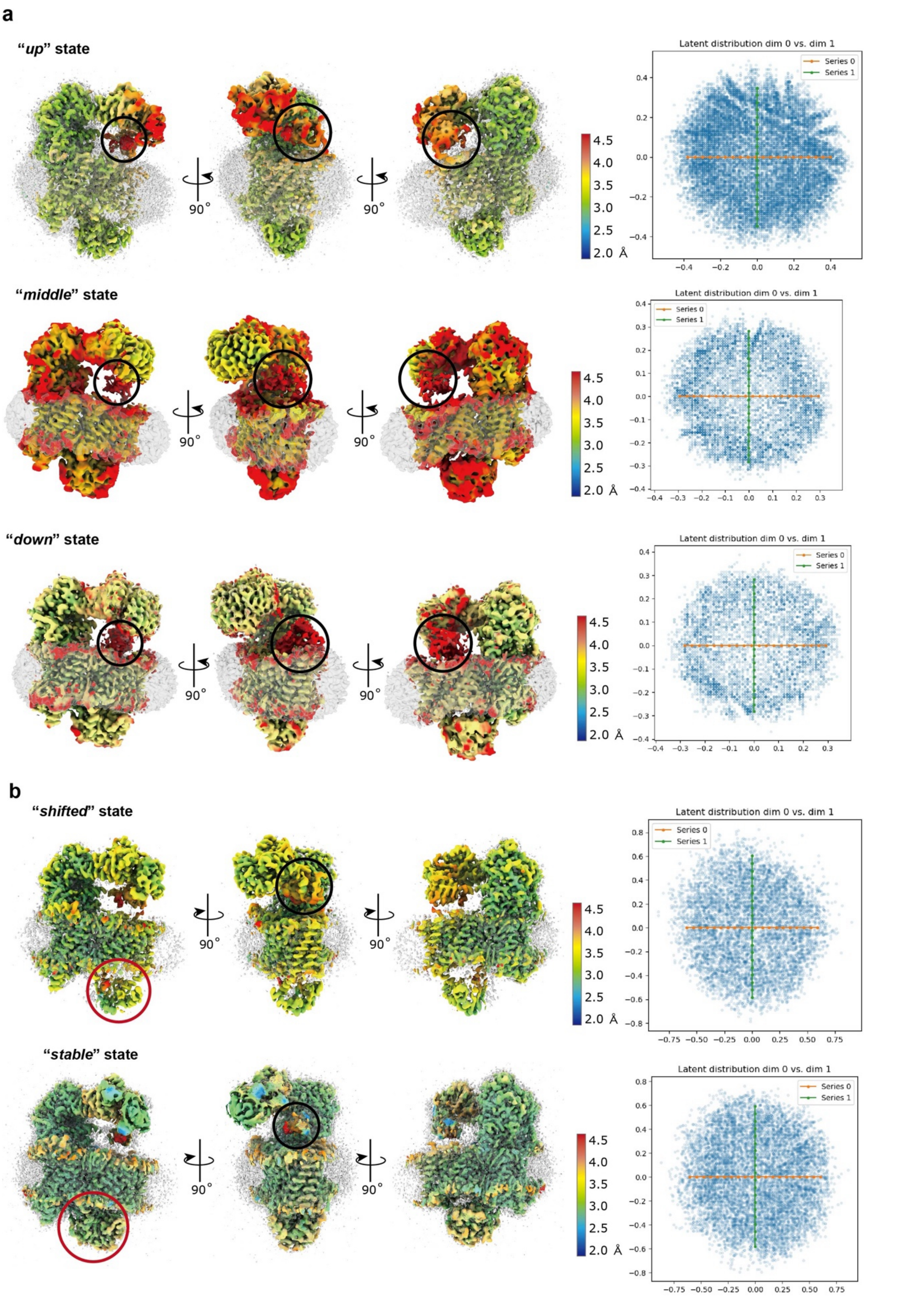


**Supplementary Fig. 2 (continued)**

**Supplementary Fig. 2: 3D Flex analysis of the complete $Na^+$-NQR.** The *left* panel shows a canonical density map from 3DFlex reconstruction colored by local resolution. The *black circle* and *red circle* indicate the ferredoxin-like domain of NqrF and the hydrophilic domain of NqrC, respectively. The *right* panel shows the latent distribution of particles produced by 3DFlex, with the orange line traversing the two major conformations. ***a,*** **$WT_{red}$/- $Na^+$**. ***b,*** **NqrB-G141$A_{red}$ + KR**.

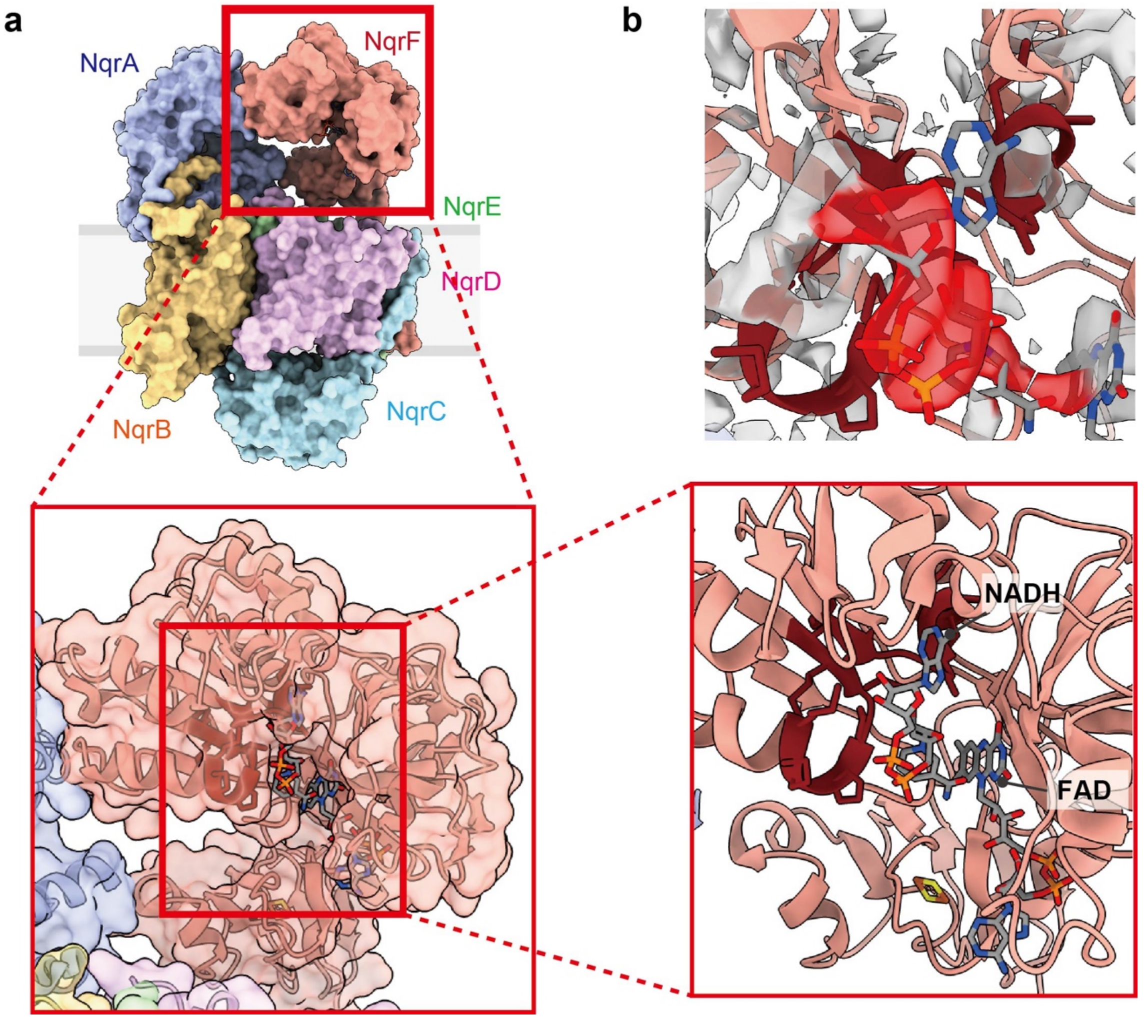


**Supplementary Fig. 3: The binding site and density map of NADH.** ***a**,* The NADH-binding site in a whole **$WT_{red}$**. *Red square* indicates the NADH-binding domain. *Dark red* residues indicate the Rossmann motif of the NADH-binding site. ***b**,* The Fo-Fc density map around NADH in NqrF; the density of NADH is colored *red*.

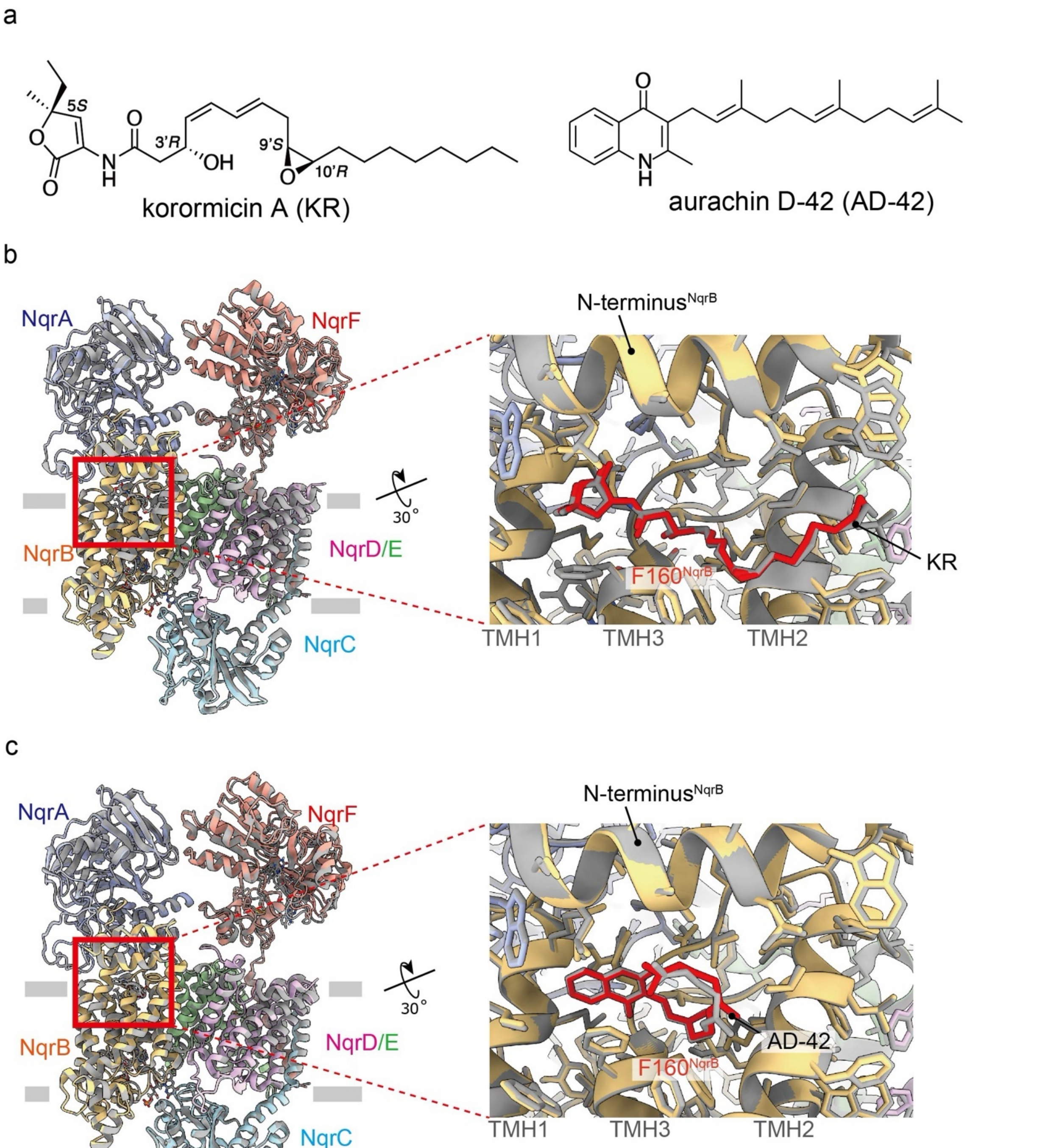


**Supplementary Fig. 4: The structures of reduced Na$^+$-NQR wild-type with bound inhibitors.** ***a,*** Chemical structure of korormicin A (KR) and aurachin D-42 (AD-42). The *gray* and *colored* indicate the structures of $\mathbf{WT_{ox}}$ and $\mathbf{WT_{red}}$ with bound inhibitors, respectively. ***b,*** Comparison of $\mathbf{WT_{ox}}$ **+ KR** and $\mathbf{WT_{red}}$ **+ KR**. ***c,*** Comparison of $\mathbf{WT_{ox}}$ **+ AD-42** vs. $\mathbf{WT_{red}}$ **+ AD-42**. The *colored* and *gray* structures correspond to $\mathbf{WT_{red}}$ and $\mathbf{WT_{ox}}$ with bound inhibitor, respectively.

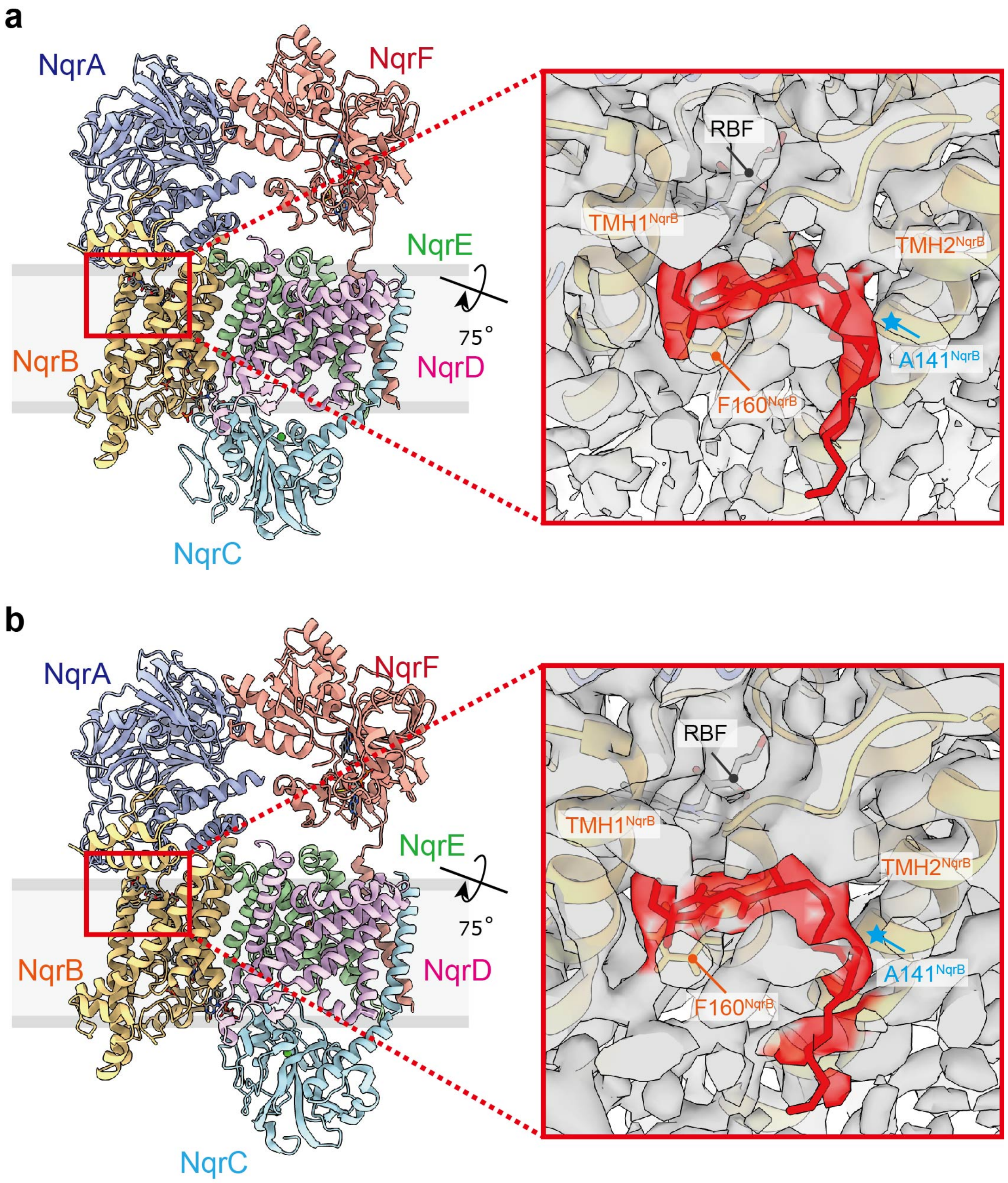


**Supplementary Fig. 5: Structure and binding form of Na$^+$-NQR inhibitor.** The structure and density map of NqrB-G141A$_{red}$ with bound KR. The density and model of KR are shown in *red*. ***a****,* The “***stable****”* state. ***b****,* The “***shifted****” st*ate.

### NqrA

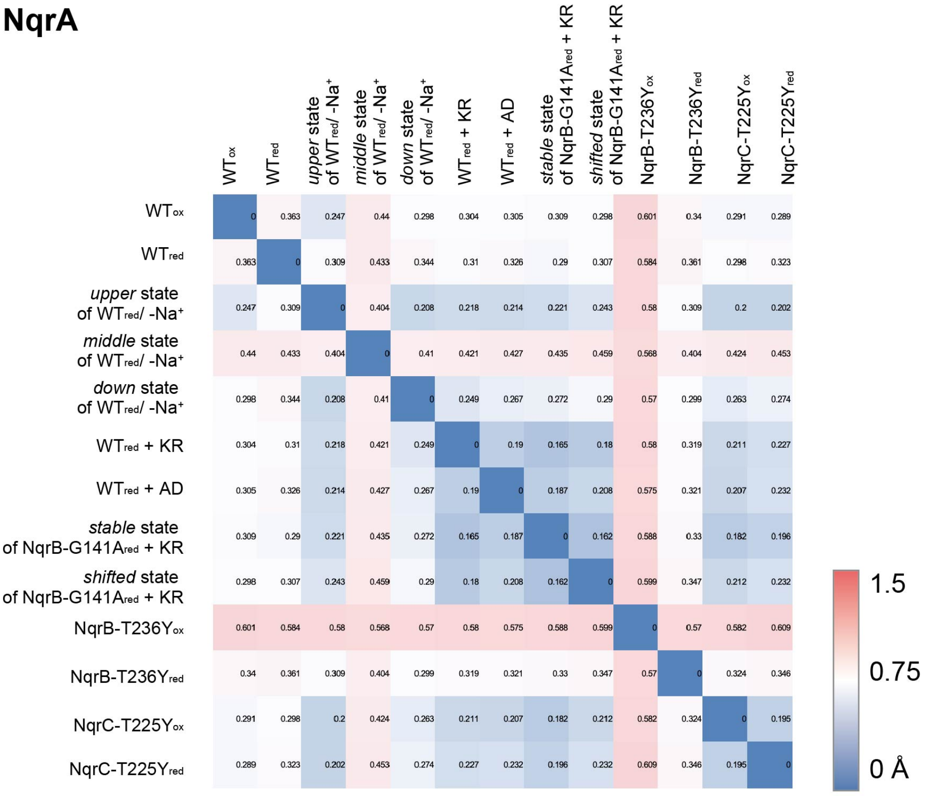


### NqrB

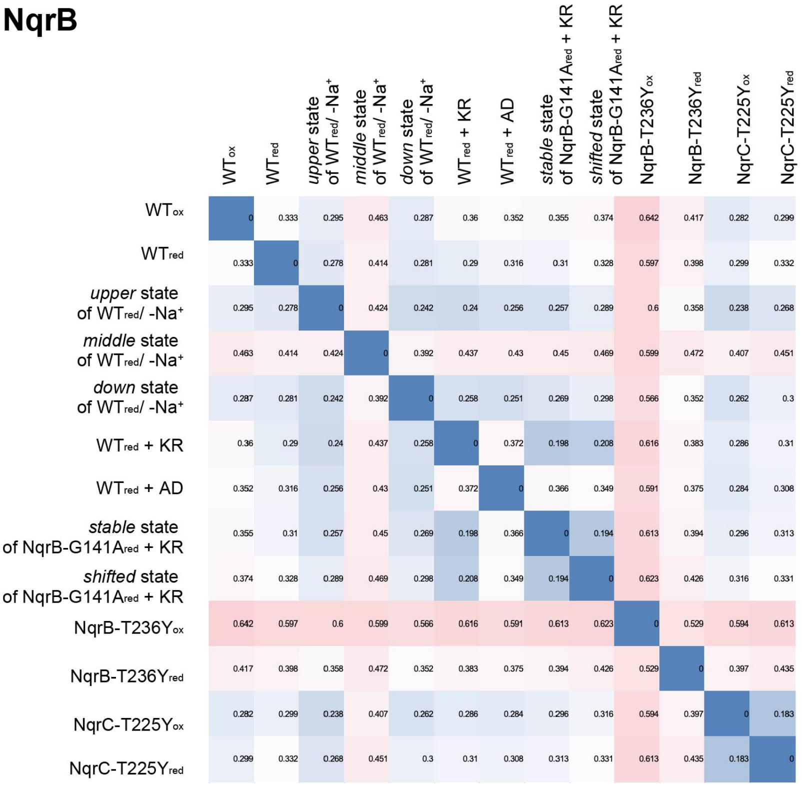


**Supplementary Fig. 6 (continued)**

### NqrC

| | WT$_{ox}$ | WT$_{red}$ | *upper* state of WT$_{red}$/ -Na$^+$ | *middle* state of WT$_{red}$/ -Na$^+$ | *down* state of WT$_{red}$/ -Na$^+$ | WT$_{red}$ + KR | WT$_{red}$ + AD | *stable* state of NqrB-G141A$_{red}$ + KR | *shifted* state of NqrB-G141A$_{red}$ + KR | NqrB-T236Y$_{ox}$ | NqrB-T236Y$_{red}$ | NqrC-T225Y$_{ox}$ | NqrC-T225Y$_{red}$ |
|---|---|---|---|---|---|---|---|---|---|---|---|---|---|
| WT$_{ox}$ | 0 | 0.467 | 0.422 | 0.58 | 0.572 | 0.45 | 0.462 | 0.447 | 1.353 | 0.656 | 0.534 | 0.552 | 0.632 |
| WT$_{red}$ | 0.467 | 0 | 0.423 | 0.57 | 0.595 | 0.423 | 0.422 | 0.376 | 1.424 | 0.673 | 0.564 | 0.654 | 0.699 |
| *upper* state of WT$_{red}$/ -Na$^+$ | 0.422 | 0.423 | 0 | 0.524 | 0.486 | 0.336 | 0.338 | 0.338 | 1.353 | 0.65 | 0.451 | 0.559 | 0.633 |
| *middle* state of WT$_{red}$/ -Na$^+$ | 0.58 | 0.57 | 0.524 | 0 | 0.452 | 0.539 | 0.537 | 0.594 | 1.373 | 0.718 | 0.581 | 0.701 | 0.758 |
| *down* state of WT$_{red}$/ -Na$^+$ | 0.572 | 0.595 | 0.486 | 0.452 | 0 | 0.523 | 0.493 | 0.568 | 1.377 | 0.712 | 0.596 | 0.596 | 0.671 |
| WT$_{red}$ + KR | 0.45 | 0.423 | 0.336 | 0.539 | 0.523 | 0 | 0.33 | 0.33 | 1.367 | 0.679 | 0.513 | 0.58 | 0.643 |
| WT$_{red}$ + AD | 0.462 | 0.422 | 0.338 | 0.537 | 0.493 | 0.33 | 0 | 0.351 | 1.437 | 0.667 | 0.499 | 0.58 | 0.662 |
| *stable* state of NqrB-G141A$_{red}$ + KR | 0.447 | 0.376 | 0.338 | 0.594 | 0.568 | 0.33 | 0.351 | 0 | 1.398 | 0.651 | 0.523 | 0.561 | 0.626 |
| *shifted* state of NqrB-G141A$_{red}$ + KR | 1.353 | 1.424 | 1.353 | 1.373 | 1.377 | 1.367 | 1.437 | 1.398 | 0 | 1.323 | 1.377 | 1.202 | 1.221 |
| NqrB-T236Y$_{ox}$ | 0.656 | 0.673 | 0.65 | 0.718 | 0.712 | 0.679 | 0.667 | 0.651 | 1.323 | 0 | 0.673 | 0.656 | 0.688 |
| NqrB-T236Y$_{red}$ | 0.534 | 0.564 | 0.451 | 0.581 | 0.596 | 0.513 | 0.499 | 0.523 | 1.377 | 0.673 | 0 | 0.641 | 0.721 |
| NqrC-T225Y$_{ox}$ | 0.552 | 0.654 | 0.559 | 0.701 | 0.596 | 0.58 | 0.58 | 0.561 | 1.202 | 0.656 | 0.641 | 0 | 0.474 |
| NqrC-T225Y$_{red}$ | 0.632 | 0.699 | 0.633 | 0.758 | 0.671 | 0.643 | 0.662 | 0.626 | 1.221 | 0.688 | 0.721 | 0.474 | 0 |

### NqrD

| | WT$_{ox}$ | WT$_{red}$ | *upper* state of WT$_{red}$/ -Na$^+$ | *middle* state of WT$_{red}$/ -Na$^+$ | *down* state of WT$_{red}$/ -Na$^+$ | WT$_{red}$ + KR | WT$_{red}$ + AD | *stable* state of NqrB-G141A$_{red}$ + KR | *shifted* state of NqrB-G141A$_{red}$ + KR | NqrB-T236Y$_{ox}$ | NqrB-T236Y$_{red}$ | NqrC-T225Y$_{ox}$ | NqrC-T225Y$_{red}$ |
|---|---|---|---|---|---|---|---|---|---|---|---|---|---|
| WT$_{ox}$ | 0 | 0.403 | 0.389 | 0.588 | 0.506 | 0.408 | 0.377 | 0.431 | 0.703 | 0.556 | 0.458 | 0.777 | 0.77 |
| WT$_{red}$ | 0.403 | 0 | 0.317 | 0.56 | 0.555 | 0.315 | 0.324 | 0.287 | 0.61 | 0.504 | 0.357 | 0.654 | 0.645 |
| *upper* state of WT$_{red}$/ -Na$^+$ | 0.389 | 0.317 | 0 | 0.558 | 0.493 | 0.282 | 0.257 | 0.264 | 0.562 | 0.505 | 0.305 | 0.652 | 0.638 |
| *middle* state of WT$_{red}$/ -Na$^+$ | 0.588 | 0.56 | 0.558 | 0 | 0.52 | 0.592 | 0.557 | 0.561 | 0.753 | 0.649 | 0.614 | 0.844 | 0.86 |
| *down* state of WT$_{red}$/ -Na$^+$ | 0.506 | 0.555 | 0.493 | 0.52 | 0 | 0.511 | 0.436 | 0.526 | 0.727 | 0.633 | 0.548 | 0.811 | 0.772 |
| WT$_{red}$ + KR | 0.408 | 0.315 | 0.282 | 0.592 | 0.511 | 0 | 0.262 | 0.23 | 0.527 | 0.498 | 0.329 | 0.604 | 0.556 |
| WT$_{red}$ + AD | 0.377 | 0.324 | 0.257 | 0.557 | 0.436 | 0.262 | 0 | 0.276 | 0.58 | 0.531 | 0.309 | 0.66 | 0.653 |
| *stable* state of NqrB-G141A$_{red}$ + KR | 0.431 | 0.287 | 0.264 | 0.561 | 0.526 | 0.23 | 0.276 | 0 | 0.51 | 0.497 | 0.355 | 0.594 | 0.567 |
| *shifted* state of NqrB-G141A$_{red}$ + KR | 0.703 | 0.61 | 0.562 | 0.753 | 0.727 | 0.527 | 0.58 | 0.51 | 0 | 0.728 | 0.544 | 0.37 | 0.376 |
| NqrB-T236Y$_{ox}$ | 0.556 | 0.504 | 0.505 | 0.649 | 0.633 | 0.498 | 0.531 | 0.497 | 0.728 | 0 | 0.536 | 0.836 | 0.84 |
| NqrB-T236Y$_{red}$ | 0.458 | 0.357 | 0.305 | 0.614 | 0.548 | 0.329 | 0.309 | 0.355 | 0.544 | 0.536 | 0 | 0.607 | 0.622 |
| NqrC-T225Y$_{ox}$ | 0.777 | 0.654 | 0.652 | 0.844 | 0.811 | 0.604 | 0.66 | 0.594 | 0.37 | 0.836 | 0.607 | 0 | 0.239 |
| NqrC-T225Y$_{red}$ | 0.77 | 0.645 | 0.638 | 0.86 | 0.772 | 0.556 | 0.653 | 0.567 | 0.376 | 0.84 | 0.622 | 0.239 | 0 |

**Supplementary Fig. 6 (continued)**

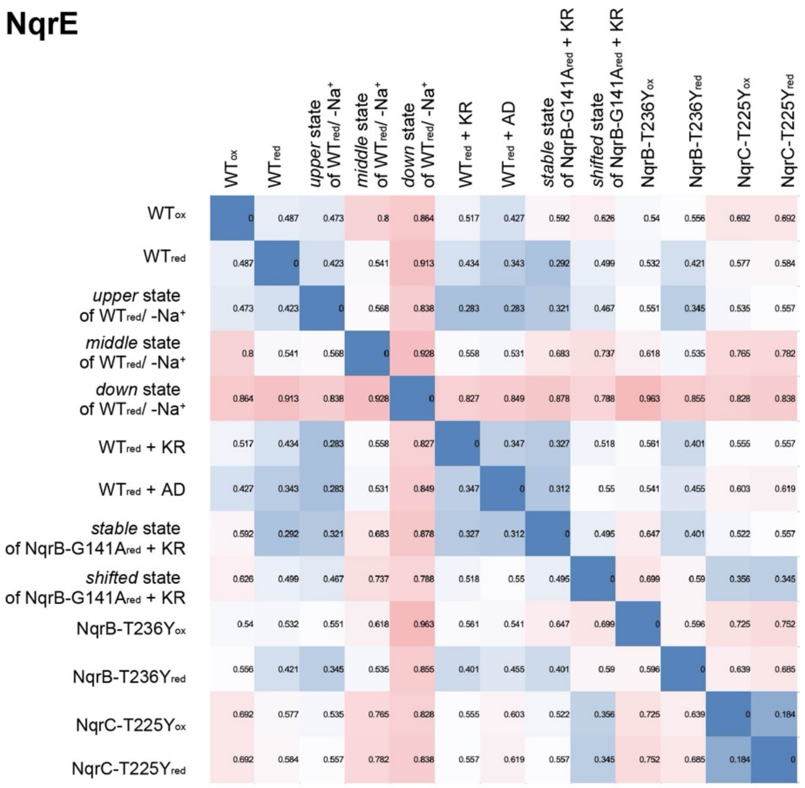


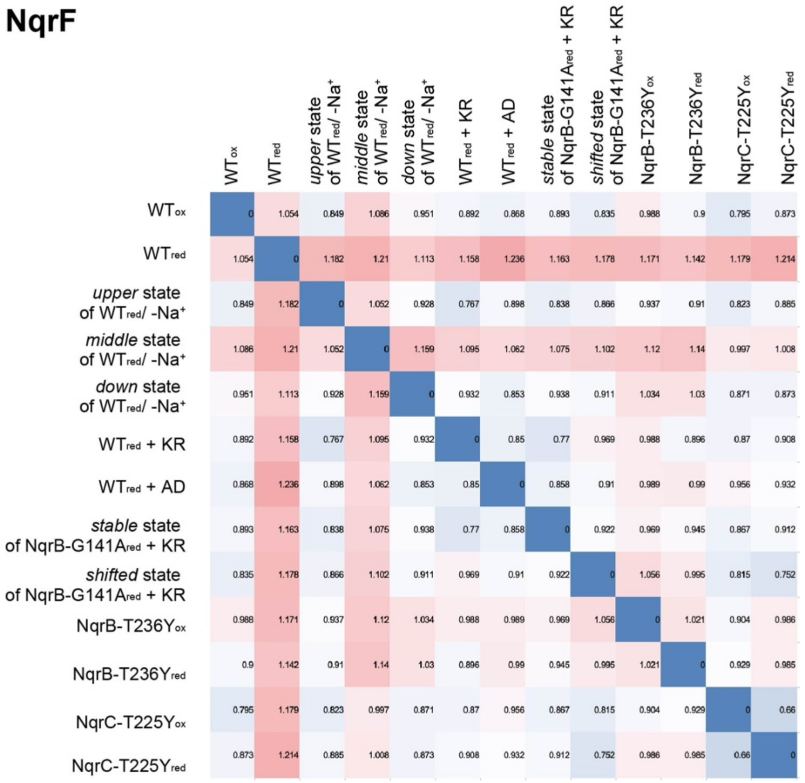


**Supplementary Fig. 6: The structural similarities of each subunit.** The heat maps of NqrA-F subunits based on r.m.s.d value. The value is calculated with UCSF chimera.

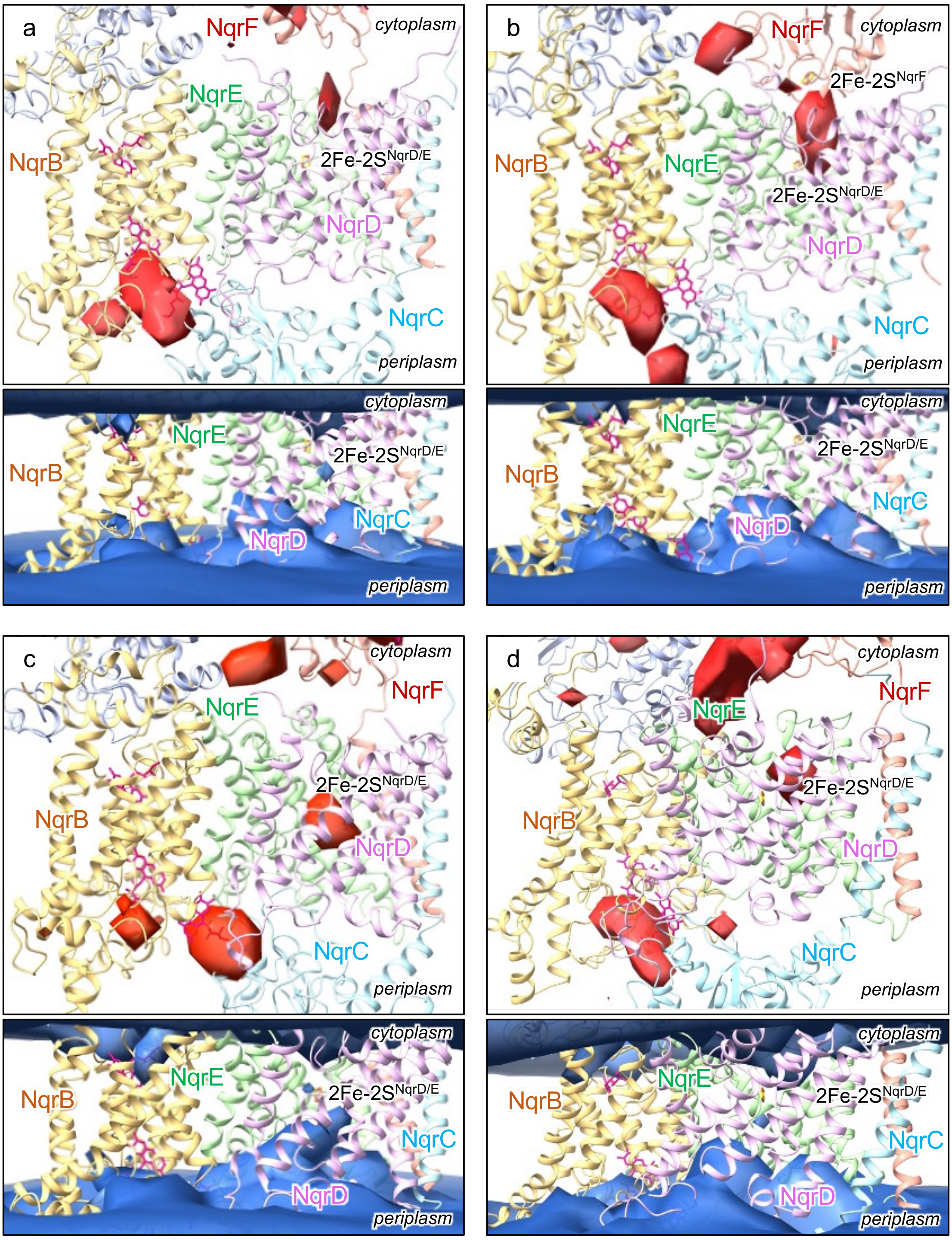


**Supplementary Fig. 7: $Na^+$ and water spatial densities from conventional MD simulations.** Spatial densities from ***a****:* cMD3, ***b****:* cMD4, ***c****:* cMD5, and ***d****:* cMD6 simulations are shown. See **Supplementary Table 2** for details of each simulation. The density of $Na^+$ is shown in red. The relative iso-density surface of 3.5 to the bulk value is shown. The density of water is shown in cyan. The relative iso-density surface of 0.3 to the bulk value is shown.

**a NqrD**

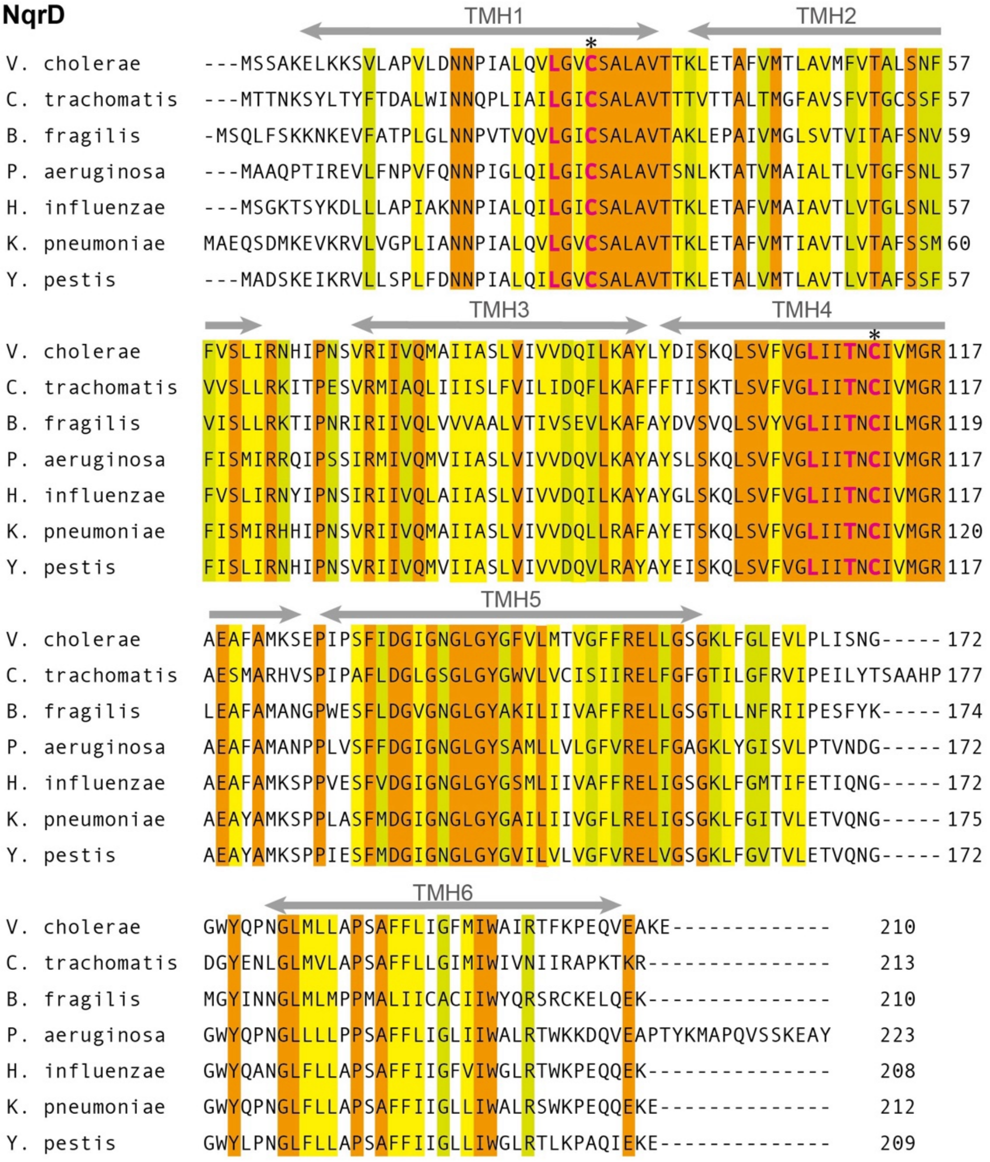


**Supplementary Fig. 8 (continued)**

**b NqrE**

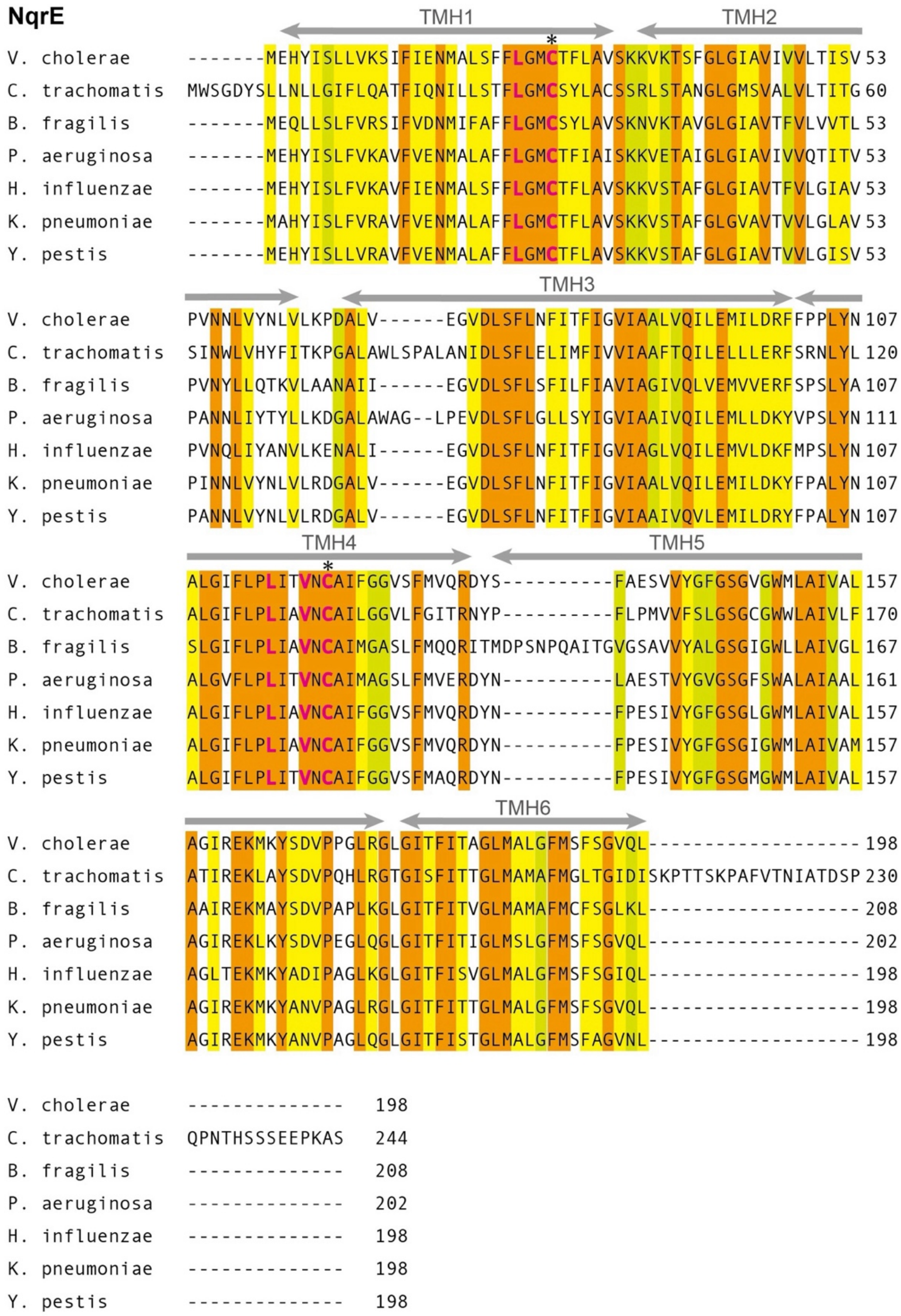


**Supplementary Fig. 8 (continued)**

**c Na⁺-NQR vs RNF**

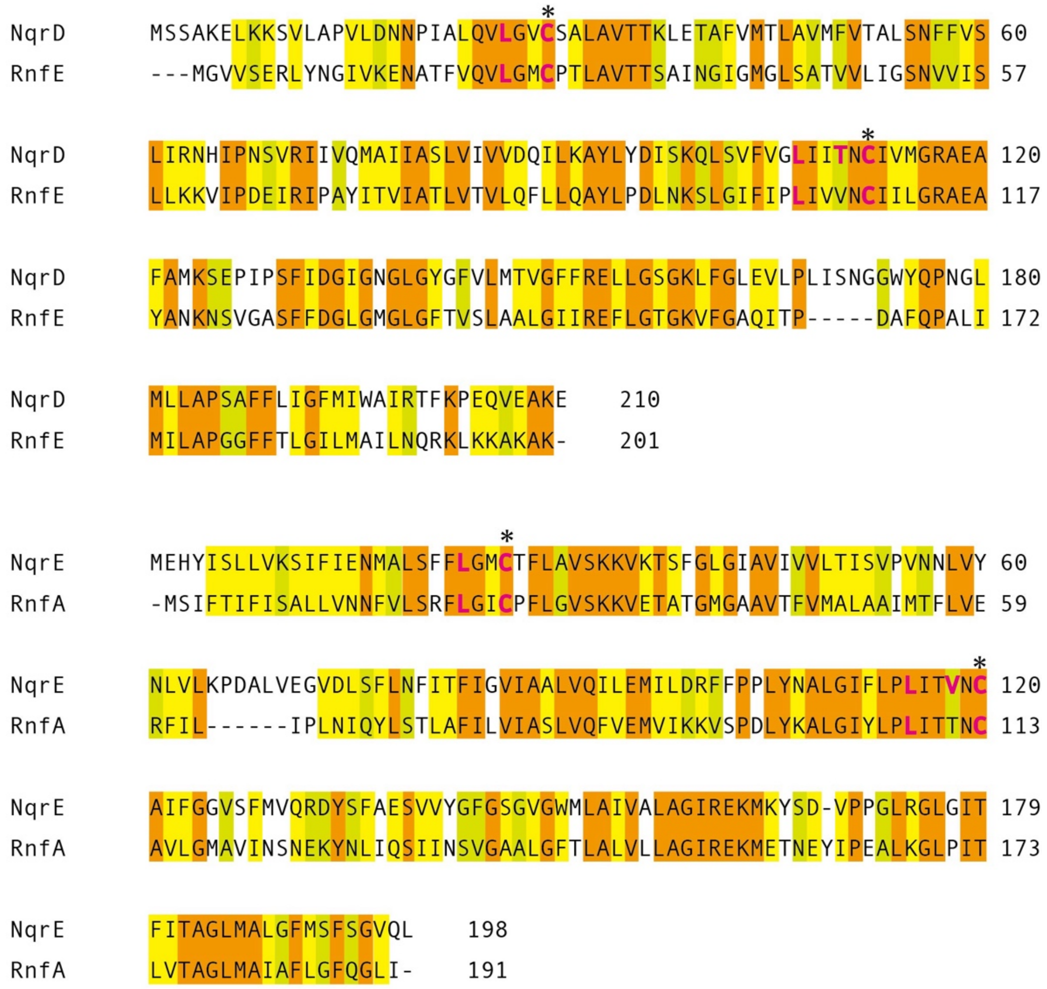


**Supplementary Fig. 8: Conserved amino acid residues in NqrD and NqrE.** This alignment is implemented using crustal omega (https://www.ebi.ac.uk/jdispatcher/msa/clustalo). ***a*** and ***b*** show NqrD and NqrE alignment, respectively. ***c***, The comparison of NqrD/E of $Na^+$-NQR from *V. cholerae* and RnfA/E of RNF from *A. woodii*. The o*range* marker indicates full-conserved residue. The y*ellow* marker indicates strong similarity (scoring > 0.5 in the Gonnet PAM 250 matrix), and the *light green* marker indicates weak similarity (scoring = < 0.5 in the Gonnet PAM 250 matrix). Four cysteines coordinating an iron-sulfur cluster ($Cys20^{NqrD}$, $Cys20^{NqrD}$, $Cys20^{NqrE}$, and $Cys20^{NqrE}$) are denoted by *asterisk* (*). The residues essential for $Na^+$ ion binding (Leu26, Cys29, Leu107, Thr110, and Cys112 of NqrD, and Leu23, Cys26, Leu115, Val118, and Cys120 of NqrE) are denoted by *pink* letters. The alpha helices are shown by *gray* arrows.

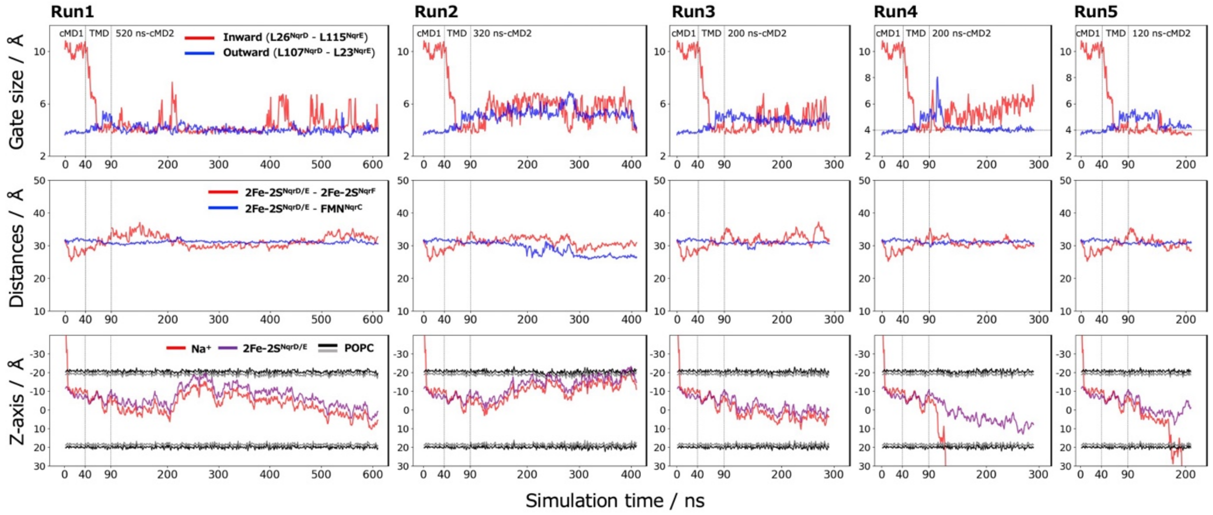


**Supplementary Fig. 9: Additional five independent trajectories of cMD2 following TMD.** *Upper column*: The inward gate and outward gate distances are plotted. *Middle column*: The distances between 2Fe-2S$^{NqrD/E}$ − 2Fe-2S$^{NqrF}$ and 2Fe-2S$^{NqrD/E}$ − FMN$^{NqrC}$ are plotted. *Bottom column*: The z coordinates of the Na$^{+}$ and 2Fe-2S$^{NqrD/E}$ in the NqrD/E transmembrane region are plotted with the membrane positions.

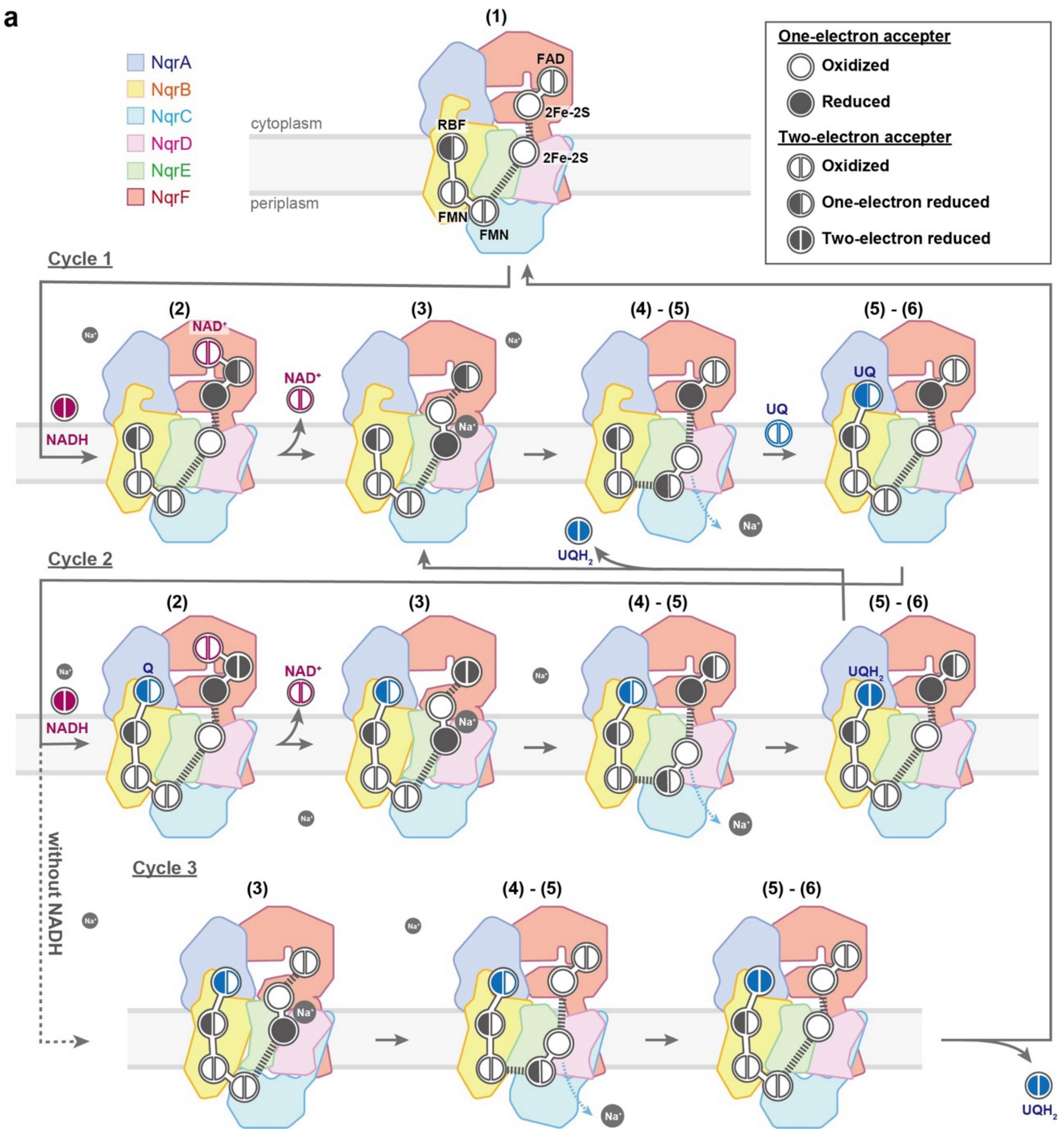


**Supplementary Fig. 10 (continued)**

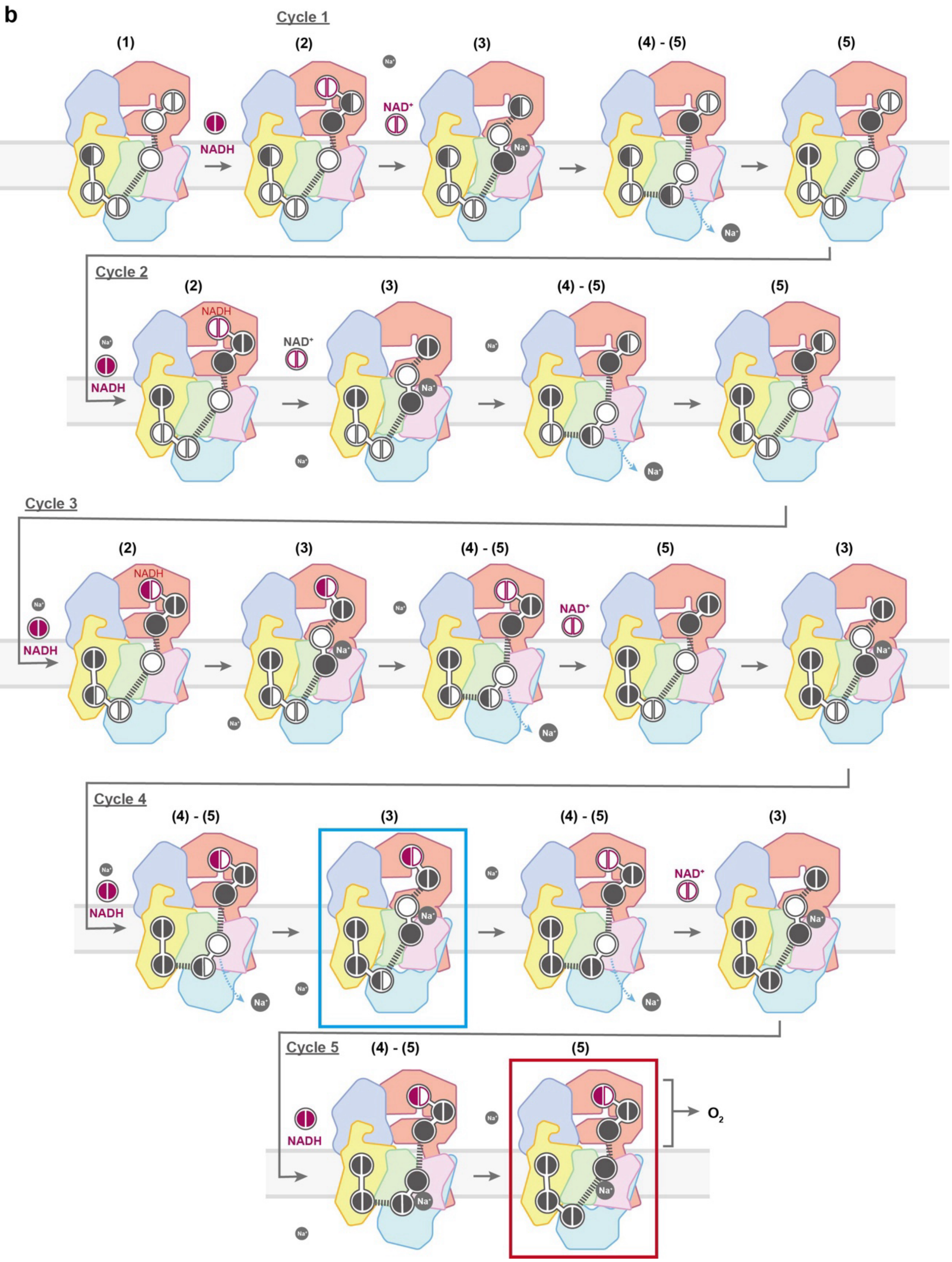


**Supplementary Fig. 10 (continued)**

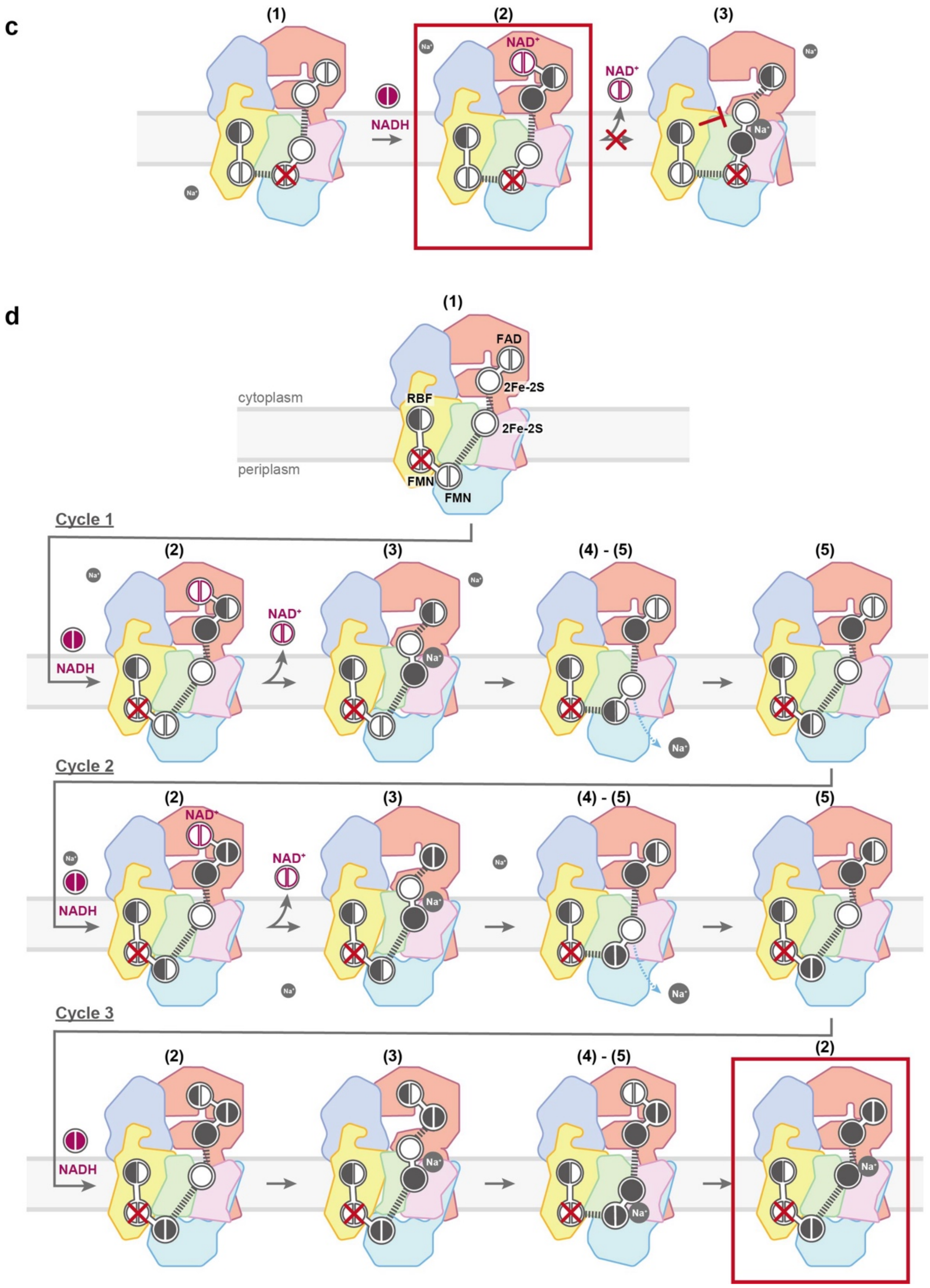


**Supplementary Fig. 10 (continued)**

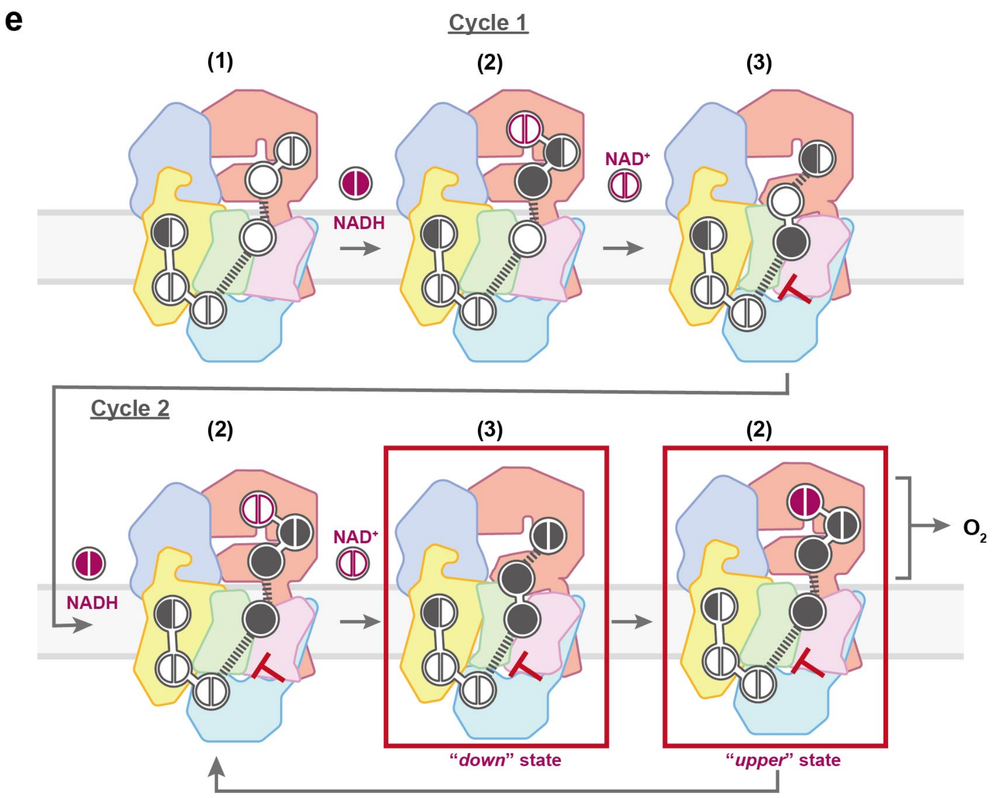


**Supplementary Fig. 10: The proposed catalytic mechanism of $Na^+$-NQR.** One-electron cofactors (e.g. 2Fe-2Ss) are shown as *full circles*; two-electron cofactors (e. g. NADH, FAD, FMNs, RBF, and UQ) are shown as two *semicircles*. *Black filled circles* or *semicircles* indicate reduction by one electron. When cofactors are close enough for electron transfer, the connection is shown as a *solid white bridge*; distances too long for electron transfer are shown with *dashes*. Since oxidized state of RBF is reported to be a stable semiquinone, it is shown as two *semicircles*, one of which is always *black*. All snapshots were assigned to step 1-6 described in the discussion. Something to consider: The transitions between intermediates are marked to indicate the corresponding step number (1-6) from the mechanism described in the discussion section. ***a,*** The reaction of wild-type enzyme with NADH and ubiquinone (UQ). ***b,*** The reaction of wild-type enzyme with NADH in the absence of UQ. The final state (*red square*) is proposed to correspond to the experimentally determined structure of $\mathbf{WT_{red}}$. ***c,*** The reaction of the NqrC-T225 mutant with NAHD in the absence of UQ. The final state (*red square*) is proposed to correspond to the experimentally determined structure of $\mathbf{WT_{red}}$. Starting from intermediate (1) equivalent to $\mathbf{NqrC\text{-}T225Y_{ox}}$, NqrC-T225Y converges into a final state (*red square*), which is proposed to correspond to the experimentally determined structure of $\mathbf{NqrC\text{-}T225Y_{red}}$. ***d,*** The reaction of the NqrB-T236Y mutant with NADH in the absence of UQ. Starting from intermediate (1) equivalent to $\mathbf{NqrB\text{-}T236Y_{ox}}$, NqrB-T236Y converges into a final state (*red square*), which is proposed to correspond to the experimentally determined structure of $\mathbf{NqrB\text{-}T236Y_{red}}$. ***e,*** The reaction of wild-type enzyme with NADH in the absence of UQ and $Na^+$. The reaction schemes that include fully reduced FAD, and two 2Fe-2Ss at step (3) and step (2) enclosed in red *squares* correspond to "***down***" and "***up***" states in $\mathbf{WT_{red}}$**/ -**$\mathbf{Na^+}$. The "***middle***" state is probably present as an intermediate between "***down***" and "***up***" state.

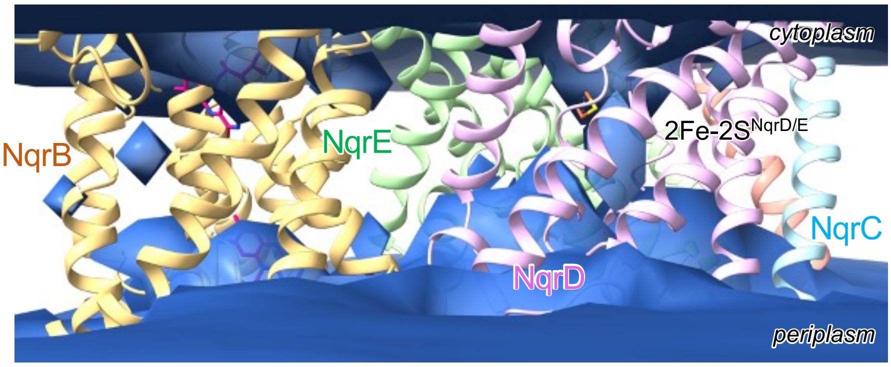


**Supplementary Fig. 11: Water spatial density from conventional MD (cMD1).** The accessibility of water is shown in the model based on $\mathbf{WT_{ox}}$ (PDB ID; 7XK3) containing reduced 2Fe-2S$^{NqrD/E}$. The relative water density with iso-density surface of 0.1 to the bulk value is shown in cyan, as shown in Figure 5b.

**Supplementary Table 1: Statics of cryo-EM data, refinement, and validation of $Na^+$-NQR.**

| | $WT_{red}$ | NqrC-$T225Y_{ox}$ | NqrC-$T225Y_{red}$ |
|---|---|---|---|
| **Data collection and processing** | | | |
| Magnification | 81,000 | 81,000 | 81,000 |
| Voltage / kV | 300 | 300 | 300 |
| Total dose / $e^-$ $Å^{-2}$ | 60 | 60 | 60 |
| Pixel size / Å | 0.88 | 0.88 | 0.88 |
| Defocus range / µm | -0.8 to -2.0 | -0.8 to -2.0 | -0.8 to -2.0 |
| Symmetry imposed | C1 | C1 | C1 |
| Recorded movies | 6,086 | 13,302 | 13,316 |
| Initial particles | 1,391,844 | 2,497,690 | 2,378,419 |
| **States** | | | |
| EMDB ID | EMDB-64518 | EMDB-63872 | EMDB-64059 |
| PDB ID | 9UUU | 9U5G | 9UD2 |
| Final particle | 245,271 | 579,365 | 1,436,839 |
| Resolution ($FSC_{0.143}$) (Å) | 3.1 | 2.7 | 2.6 |
| **Model refinement** | | | |
| Model resolution ($FSC_{0.5}$) (Å) | 3.6 | 2.7 | 2.6 |
| **Composition** | | | |
| Non-hydrogen atoms | 14694 | 14765 | 14763 |
| Residues | 1892 | 1900 | 1900 |
| Ligands | FMN: 2, FES: 2, RBF: 1, FAD: 1, NAI:1, CA: 1 | FMN: 1, FES: 2, RBF: 1, FAD: 1, CA: 1, LMT: 2 | FMN: 1, FES: 2, RBF: 1, FAD: 1, CA: 1, LMT: 2 |
| Waters | 0 | 7 | 5 |
| Bond length (Å) | 0.007 | 0.005 | 0.005 |
| Bond angles (°) | 0.871 | 0.608 | 0.698 |
| Clash score | 5.09 | 3.07 | 3.54 |
| MolProbity score | 1.65 | 1.33 | 1.61 |
| **Ramachandran plot (%)** | | | |
| Favored | 94.36 | 96.50 | 96.56 |
| Allowed | 5.59 | 3.5 | 3.34 |
| Outlier | 0.05 | 0.00 | 0.11 |
| Rotamer outlier (%) | 0 | 0.90 | 1.61 |

**Supplementary Table 1 (continued)**

| | NqrB-T236Y$_{ox}$ | NqrB-T236Y$_{red}$ |
|---|---|---|
| **Data collection and processing** | | |
| Magnification | 81,000 | 81,000 |
| Voltage / kV | 300 | 300 |
| Total dose / $e^{-}$ Å$^{-2}$ | 60 | 60 |
| Pixel size / Å | 0.88 | 0.88 |
| Defocus range / µm | -0.8 to -2.0 | -0.8 to -2.0 |
| Symmetry imposed | C1 | C1 |
| Recorded movies | 4,440 | 8212 |
| Initial particles | 1,026,158 | 2,496,421 |
| **States** | | |
| EMDB ID | EMDB-64060 | EMDB-64061 |
| PDB ID | 9UD3 | 9UD4 |
| Final particle | 79,750 | 319,845 |
| Resolution ($FSC_{0.143}$) (Å) | 3.8 | 3.3 |
| **Model refinement** | | |
| Model resolution ($FSC_{0.5}$) (Å) | 4.1 | 3.4 |
| **Composition** | | |
| Non-hydrogen atoms | 14680 | 14720 |
| Residues | 1894 | 1894 |
| Ligands | FMN: 1, FES: 2, RBF: 1, FAD: 1, PEE:2, CA: 1, LMT: 1 | FMN: 1, FES: 2, RBF: 1, FAD: 1, CA: 1, LMT: 2 |
| Waters | 0 | 5 |
| Bond length (Å) | 0.004 | 0.003 |
| Bond angles (°) | 0.693 | 0.587 |
| Clash score | 9.88 | 6.23 |
| MolProbity score | 2.28 | 1.65 |
| **Ramachandran plot (%)** | | |
| Favored | 94.74 | 96.71 |
| Allowed | 5.26 | 3.24 |
| Outlier | 0.00 | 0.05 |
| Rotamer outlier (%) | 3.37 | 1.36 |

**Supplementary Table 1 (continued)**

| | $WT_{red}$ /- $Na^+$ | | |
|---|---|---|---|
| **Data collection and processing** | | | |
| Magnification | 81,000 | | |
| Voltage / kV | 300 | | |
| Total dose / $e^-$ $Å^{-2}$ | 60 | | |
| Pixel size / Å | 0.88 | | |
| Defocus range / µm | -0.8 to -2.0 | | |
| Symmetry imposed | C1 | | |
| Recorded movies | 13,302 | | |
| Initial particles | 3,653,368 | | |
| **States** | **"*up*" sate** | **"*moderate*" sate** | **"*down*" sate** |
| EMDB ID | EMDB-64063 | EMDB-64064 | EMDB-64065 |
| PDB ID | 9UD6 | 9UD8 | 9UD9 |
| Final particle | 1,244,065 | 100,990 | 77,726 |
| Resolution ($FSC_{0.143}$) (Å) | 2.6 | 3.8 | 3.1 |
| **Model refinement** | | | |
| Model resolution ($FSC_{0.5}$) (Å) | 2.7 | 3.9 | 3.2 |
| **Composition** | | | |
| Non-hydrogen atoms | 14743 | 14706 | 14665 |
| Residues | 1894 | 1894 | 1894 |
| Ligands | FMN: 2, FES: 2, RBF: 1, FAD: 1, CA: 1, LMT: 2 | FMN: 2, FES: 2, RBF: 1, FAD: 1, PEE:3, CA: 1, LMT: 1 | FMN: 2, FES: 2, RBF: 1, FAD: 1, CA: 1 |
| Waters | 2 | 0 | 0 |
| Bond length (Å) | 0.004 | 0.004 | 0.003 |
| Bond angles (°) | 0.566 | 0.700 | 0.639 |
| Clash score | 2.50 | 8.71 | 3.40 |
| MolProbity score | 1.20 | 1.68 | 1.29 |
| **Ramachandran plot (%)** | | | |
| Favored | 97.02 | 97.45 | 97.13 |
| Allowed | 2.98 | 2.55 | 2.82 |
| Outlier | 0.00 | 0.00 | 0.05 |
| Rotamer outlier (%) | 0.58 | 1.36 | 0.84 |

**Supplementary Table 1 (continued)**

| NqrB-G141A$_{red}$ + KR | | |
|---|---|---|
| **Data collection and processing** | | |
| Magnification | 81,000 | |
| Voltage / kV | 300 | |
| Total dose / e$^-$ Å$^{-2}$ | 60 | |
| Pixel size / Å | 0.88 | |
| Defocus range / µm | -0.8 to -2.0 | |
| Symmetry imposed | C1 | |
| Recorded movies | 5,086 | |
| Initial particles | 1,384,337 | |
| **States** | **"*stable*" state** | **"*shifted*" state** |
| EMDB ID | EMDB-64066 | EMDB-64068 |
| PDB ID | 9UDA | 9UDF. |
| Final particle | 520,594 | 74,083 |
| Resolution (FSC$_{0.143}$) (Å) | 2.6 | 2.9 |
| **Model refinement** | | |
| Model resolution (FSC$_{0.5}$) (Å) | 2.8 | 3.1 |
| **Composition** | | |
| Non-hydrogen atoms | 14963 | 15124 |
| Residues | 1921 | 1922 |
| Ligands | IQT (KR):1, FMN: 2, FES: 2, RBF: 1, FAD: 1, CA: 1 | IQT (KR):1, FMN: 2, FES: 2, RBF: 1, FAD: 1, PEE:2, CA: 1, LMT: 2 |
| Waters | 37 | 17 |
| Bond length (Å) | 0.197 | 0.196 |
| Bond angles (°) | 2.777 | 2.764 |
| Clash score | 2.84 | 6.13 |
| MolProbity score | 1.13 | 1.90 |
| **Ramachandran plot (%)** | | |
| Favored | 97.75 | 95.13 |
| Allowed | 2.25 | 4.71 |
| Outlier | 0 | 0.16 |
| Rotamer outlier (%) | 0.38 | 1.98 |

**Supplementary Table 1 (continued)**

| | $WT_{red}$ + KR | $WT_{red}$ + AD |
|---|---|---|
| **Data collection and processing** | | |
| Magnification | 81,000 | 81,000 |
| Voltage / kV | 300 | 300 |
| Total dose / $e^-$ Å$^{-2}$ | 60 | 60 |
| Pixel size / Å | 0.88 | 0.88 |
| Defocus range / µm | -0.8 to -2.0 | -0.8 to -2.0 |
| Symmetry imposed | C1 | C1 |
| Recorded movies | 4,881 | 5,096 |
| Initial particles | 2,157,492 | 1,072,343 |
| **States** | | |
| EMDB ID | EMDB-64062 | EMDB-64069 |
| PDB ID | 9UD5 | 9UDG |
| Final particle | 285,018 | 390,507 |
| Resolution ($FSC_{0.143}$) (Å) | 2.9 | 2.7 (J43) |
| **Model refinement** | | |
| Model resolution ($FSC_{0.5}$) (Å) | 3.0 | 2.7 |
| Composition | | |
| Non-hydrogen atoms | 14496 | 15132 |
| Residues | 1921 | 1919 |
| Ligands | IQT (KR):1, FMN: 2, FES: 2, RBF: 1, FAD: 1, CA: 1, LMT: 2 | ONI (AD):1, FMN: 2, FES: 2, RBF: 1, FAD: 1, PEE:2, CA: 1, LMT: 2 |
| Waters | 37 | 61 |
| Bond length (Å) | 0.003 | 0.004 |
| Bond angles (°) | 0.630 | 0.624 |
| Clash score | 3.14 | 4.59 |
| MolProbity score | 1.26 | 1.40 |
| Ramachandran plot (%) | | |
| Favored | 97.17 | 97.06 |
| Allowed | 2.62 | 2.78 |
| Outlier | 0.21 | 0.16 |
| Rotamer outlier (%) | 0 | 0.38 |

**Supplementary Table 2: List of atomistic MD simulations.**

| Simulation | Initial structure | Reduced cofactor | NqrD/E conformation | | Length / ns |
|---|---|---|---|---|---|
| | | | initial | final | |
| cMD1 | 7XK3 | 2Fe-2S$^{NqrF}$,<br>2Fe-2S$^{NqrD/E}$ | inward | inward | 40 (1,000) |
| TMD | *cMD1* | 2Fe-2S$^{NqrF}$,<br>2Fe-2S$^{NqrD/E}$ | inward | outward | 50 |
| cMD2 | *TMD* | FMN$^{NqrC}$ | outward | (inward) | 800<br>520<br>320<br>200<br>200<br>120 |
| cMD3 | 7XK3 | none | inward | inward | 400 |
| cMD4 | $WT_{red}/-Na^{+}$ (down) | 2Fe-2S$^{NqrF}$,<br>2Fe-2S$^{NqrD/E}$ | inward | inward | 400 |
| cMD5 | NqrB-G141$A_{red}$ +<br>KR (*shifted*) | 2Fe-2S$^{NqrF}$,<br>2Fe-2S$^{NqrD/E}$ | outward | outward | 400 |
| cMD6 | NqrB-G141$A_{red}$ +<br>KR (*stable*) | FAD,<br>2Fe-2S$^{NqrF}$,<br>2Fe-2S$^{NqrD/E}$<br>FMN$^{NqrC}$ | inward | inward | 400 |

**Supplementary Table 3: $Na^+$ binding site by MD simulation.**[a]

| Subunit | Binding site | Contact ratio |
|---|---|---|
| **$WT_{ox}$: full oxidized** | | |
| NqrB | Thr236 | 0.90 |
| | Ser221 | 0.50 |
| | Gly245 | 0.27 |
| | Gln243 | 0.22 |
| | Trp241 | 0.22 |
| | Asp223 | 0.21 |
| | Ser239 | 0.18 |
| | Ala248 | 0.12 |
| NqrD | Glu119 | 0.025 |
| | Ser170 | 0.023 |
| | Gly172 | 0.022 |
| | Glu210 | 0.022 |
| | Asn171 | 0.020 |
| NqrE | Asp133 | -0.023 |
| | Glu95 | 0.020 |
| | Gln92 | 0.017 |
| | Asp99 | 0.016 |
| | Pro114 | 0.016 |
| **$WT_{red}$: reduced $2Fe\text{-}2S^{NqrF}$ and $2Fe\text{-}2S^{NqrD/E}$** | | |
| NqrB | Thr236 | 1.0 |
| | Ser239 | 0.46 |
| | Ser221 | 0.14 |
| | Asp223 | 0.12 |
| | Gln240 | 0.074 |
| NqrD | Cys112 | 0.37 |
| | Thr110 | 0.24 |
| | Glu164 | 0.058 |
| | Lys202 | 0.031 |
| | Arg199 | 0.030 |
| NqrE | Val118 | 0.11 |
| | Thr117 | 0.029 |
| | Leu113 | 0.018 |
| | Val73 | 0.017 |
| | Glu92 | 0.017 |

| | | |
|---|---|---|
| **“down” state of $WT_{red}$/ -$Na^+$: reduced 2Fe-2S$^{NqrF}$ and 2Fe-2S$^{NqrD/E}$** | | |
| NqrB | Thr236 | 1 |
| | Ser239 | 0.88 |
| | Asp223 | 0.18 |
| | His13 | 0.051 |
| | Gly234 | 0.046 |
| NqrD | Cys112 | 0.084 |
| | Thr110 | 0.079 |
| | Val24 | 0.063 |
| | Glu210 | 0.041 |
| | Glu204 | 0.030 |
| | Gly172 | 0.025 |
| NqrE | Val118 | 0.31 |
| | Leu115 | 0.11 |
| | Cys26 | 0.063 |
| | His3 | 0.018 |
| | Leu198 | 0.012 |
| **“shifted” state of NqrB-G141$A_{red}$: reduced FAD, 2Fe-2S$^{NqrF}$, 2Fe-2S$^{NqrD/E}$ and FMN$^{NqrB}$** | | |
| NqrB | Thr236 | 0.94 |
| | Ser239 | 0.34 |
| | Ser221 | 0.078 |
| | Glu106 | 0.046 |
| | His12 | 0.031 |
| NqrD | Thr110 | 0.49 |
| | Glu210 | 0.028 |
| | Cys112 | 0.027 |
| | Glu204 | 0.020 |
| | Gln100 | 0.016 |
| NqrE | Val118 | 0.48 |
| | Cys120 | 0.47 |
| | Asp67 | 0.015 |
| | Glu71 | 0.013 |
| | Glu2 | 0.013 |
| **$WT_{ox}$: reduced 2Fe-2S$^{NqrF}$ and 2Fe-2S$^{NqrD/E}$** | | |
| NqrB | Thr236 | 1.0 |
| | Ser239 | 0.40 |
| | Asp223 | 0.095 |

| | | |
|---|---|---|
| | Leu224 | 0.088 |
| | Thr173 | 0.086 |
| | Thr227 | 0.086 |
| | Ser44 | 0.048 |
| NqrD | Cys112 | 0.30 |
| | Thr110 | 0.21 |
| | Glu204 | 0.029 |
| | Gln24 | 0.021 |
| | Glu210 | 0.019 |
| NqrE | Val118 | 0.66 |
| | Leu115 | 0.45 |
| | Gln197 | 0.067 |
| | Gly195 | 0.060 |
| | Asp67 | 0.021 |
| **"*shifted*" state of NqrB-G141A$_{red}$: reduced 2Fe-2S$^{NqrF}$ and 2Fe-2S$^{NqrD/E}$** | | |
| NqrB | Thr236 | 0.95 |
| | Ser239 | 0.60 |
| | Trp226 | 0.11 |
| | His101 | 0.052 |
| | His12 | 0.042 |
| NqrD | Thr110 | 0.58 |
| | Cys112 | 0.19 |
| | Glu210 | 0.027 |
| | Gly172 | 0.025 |
| | Asn171 | 0.018 |
| NqrE | Val118 | 0.57 |
| | Cys120 | 0.55 |
| | Gln197 | 0.082 |
| | Ala68 | 0.019 |
| | Leu198 | 0.018 |
| **"*stable*" state of NqrB-G141A$_{red}$: reduced FAD, 2Fe-2S$^{NqrF}$, 2Fe-2S$^{NqrD/E}$ and FMN$^{NqrC}$** | | |
| NqrB | Thr236 | 1 |
| | Ser239 | 0.33 |
| | Gly234 | 0.15 |
| | Pro269 | 0.15 |
| | Asp223 | 0.056 |
| NqrD | Chs112 | 0.13 |

| | | |
|---|---|---|
| | Thr110 | 0.086 |
| | Gly172 | 0.026 |
| | Glu204 | 0.026 |
| | Glu210 | 0.025 |
| NqrE | Val118 | 0.85 |
| | Leu115 | 0.72 |
| | Lys33 | 0.055 |
| | Glu172 | 0.039 |
| | Gln197 | 0.028 |
| **"*shifted*" state of NqrB-G141A$_{red}$: reduced 2Fe-2S$^{NqrF}$ and 2Fe-2S$^{NqrD/E}$** | | |
| NqrB | Thr236 | 0.95 |
| | Ser239 | 0.60 |
| | Trp226 | 0.11 |
| | His101 | 0.052 |
| | His12 | 0.042 |
| NqrD | Thr110 | 0.58 |
| | Cys112 | 0.19 |
| | Glu210 | 0.027 |
| | Gly172 | 0.025 |
| | Asn171 | 0.018 |
| NqrE | Val118 | 0.57 |
| | Cys120 | 0.55 |
| | Gln197 | 0.082 |
| | Ala68 | 0.019 |
| | Leu198 | 0.018 |
| **"*stable*" state of NqrB-G141A$_{red}$: reduced FAD, 2Fe-2S$^{NqrF}$, 2Fe-2S$^{NqrD/E}$ and FMN$^{NqrC}$** | | |
| NqrB | Thr236 | 1 |
| | Ser239 | 0.33 |
| | Gly234 | 0.15 |
| | Pro269 | 0.15 |
| | Asp223 | 0.056 |
| NqrD | Chs112 | 0.13 |
| | Thr110 | 0.086 |
| | Gly172 | 0.026 |
| | Glu204 | 0.026 |
| | Glu210 | 0.025 |
| NqrE | Val118 | 0.85 |

| | | |
|---|---|---|
| | Leu115 | 0.72 |
| | Lys33 | 0.055 |
| | Glu172 | 0.039 |
| | Gln197 | 0.028 |

[a] Simulation is calculated at the scale of 400 ns. Contact ratio corresponds to the ratio of $Na^+$-binding frame to total frame. Residues showing > 0.01 of contact ratio and the > 3 Å of distance to were listed up.
